# Limit-pushing overexpression reveals constraints on protein abundance

**DOI:** 10.64898/2026.09.24.754258

**Authors:** Shotaro Namba, Yuki Nishida, Charles Boone, Brenda Andrews, Hisao Moriya

**Affiliations:** Faculty of Environmental, Life, Natural Science and Technology, Okayama University, Okayama, Japan; Graduate School of Environmental, Life, Natural Science and Technology, Okayama University, Okayama, Japan; Donnelly Centre for Cellular and Biomolecular Research and Department of Molecular Genetics, University of Toronto, Toronto, ON, Canada; Department of Molecular Genetics, University of Toronto, Toronto, ON, Canada; RIKEN Center for Sustainable Resource Science, Wako, Saitama 351-0198, Japan; Canadian Institute for Advanced Research (CIFAR), Toronto, ON, Canada

## Abstract

Proteins are often classified as toxic or non-toxic without measuring the abundance reached, leaving constraints on tolerable protein abundance unresolved. We established a limit-pushing approach in *Saccharomyces cerevisiae* combining strong inducible expression with gTOW-mediated high-copy selection to counteract copy-number compensation while measuring protein abundance and growth. Nearly all of approximately 80 chromosome I proteins severely inhibited growth or reduced viability at sufficiently high abundance. We established IE50, the expression level associated with a 50% reduction in growth rate, to quantify their widely varying overexpression tolerance. IE50 was positively associated with predicted structural order and cytoplasmic localization propensity and negatively associated with sulphur content. Single-cell imaging linked higher tolerance to proteins remaining cytoplasmic without becoming aggregation-positive and revealed abundance-dependent changes in localization and organelle morphology. At extreme abundance, Fun12, Nup60, and Pex22 generated distinct large-scale intracellular states through specific sequence regions. These findings establish overexpression toxicity as a quantitative property linked to protein characteristics and reveal both constraints on tolerable abundance and sequence-dependent capacities for intracellular organization.

## Introduction

Protein abundances within cells span several orders of magnitude across different proteins. Because protein abundance directly affects cellular function and fitness, expression level itself is an important trait subject to evolutionary optimization (Hill et al., 2021; Huang et al., 2023; Wang et al., 2012). In general, the optimal abundance of a protein is thought to reflect a balance between the functional benefits provided by the protein and the costs associated with its production and maintenance (Bédard et al., 2022; Cherry, 2010; Dekel & Alon, 2005; Fujita et al., 2025; Venkataraman & Landry, 2026). Indeed, quantitative analyses of expression–fitness relationships have shown that fitness can decrease when protein expression is either too low or too high, resulting in an intermediate optimal expression level (Chou et al., 2014; Hawkins et al., 2020; Keren et al., 2016; Mathis et al., 2021).

The consequences of protein overexpression, however, differ markedly among proteins. Some proteins impair growth even when their expression is increased only modestly, whereas others can accumulate to high levels with little apparent effect on growth (Eguchi et al., 2018; Keren et al., 2016; Tomala & Korona, 2013). At least two types of effects are thought to underlie these differences. One is a general burden on shared cellular resources associated with protein production and maintenance, including transcription, translation, folding, and degradation (Kafri et al., 2016; Stoebel et al., 2008). The other is protein-specific toxicity arising from mechanisms such as stoichiometric imbalance, excessive or promiscuous molecular interactions, inappropriate enzymatic activity, misfolding, and aggregation (Bhattacharyya et al., 2016; Levy et al., 2012; Moriya, 2015; Papp et al., 2003; Vavouri et al., 2009). Even proteins that impose little burden beyond the general cost of their production should eventually reduce fitness when expressed at sufficiently high levels, because cellular resources are finite (Eguchi et al., 2018; Fujita et al., 2025).

If so, classifying proteins simply as “toxic” or “non-toxic” upon overexpression is insufficient. A more informative quantity is the expression limit: how much of each protein a cell can tolerate before its fitness is substantially compromised. Comparing these limits among proteins should help distinguish general expression burden from protein-specific toxicity and reveal the physicochemical and cell-biological properties that determine tolerance to protein overexpression.

The effects of protein overexpression have been studied extensively in the budding yeast *Saccharomyces cerevisiae*, in which gene dosage can be readily manipulated. The development of multicopy plasmids and strong inducible or constitutive expression systems has enabled multiple systematic genome-wide overexpression screens (Arita et al., 2021; Douglas et al., 2012; Gelperin et al., 2005; Makanae et al., 2013; Sopko et al., 2006; Tomala & Korona, 2013; Yoshikawa et al., 2011) (**Figure S1A**). Integrating these studies, 2,891 genes have been reported to inhibit growth when overexpressed in at least one study, whereas no overexpression-associated growth defect has been reported for 3,237 genes (**Figure S1B**). It remains unclear, however, whether this toxic/non-toxic classification reflects intrinsic differences in overexpression tolerance or simply whether each expression system was capable of producing enough protein to reach a toxic level.

Two limitations of conventional overexpression systems make this distinction difficult. First, transcriptional and translational efficiencies impose limits on achievable expression. In addition, when genes are expressed from multicopy plasmids, cells carrying fewer plasmid copies can be selectively enriched when high expression reduces fitness, thereby buffering the effective expression level (Eguchi et al., 2018; Moriya et al., 2006). We previously developed the genetic tug-of-war (gTOW) method to counter this copy-number compensation and drive gene copy number toward its upper limit (Moriya et al., 2006, 2012) (**Figure S1C**). Second, most previous studies did not simultaneously quantify actual protein abundance and its effect on growth. Consequently, it has often been impossible to distinguish whether an apparently non-toxic protein is genuinely tolerated at high abundance or simply fails to reach the abundance at which growth inhibition would occur. A quantitative analysis of overexpression toxicity therefore requires both pushing protein abundance toward its limit and measuring protein abundance and fitness simultaneously (Moriya, 2015).

The unresolved issue is therefore whether proteins have been driven to sufficiently high abundance to reveal their constraints, rather than simply which genes have been classified as toxic. Addressing this issue requires measuring abundance, fitness, and cellular state in the same targets while pushing expression toward its limit. We combined the strong WTC846 inducible expression system (Azizoglu et al., 2021) with gTOW-mediated high-copy selection, building on our previous use of this combination for extreme protein overexpression (Namba & Moriya, 2024). By counteracting selection for reduced plasmid copy number, this approach extends the accessible expression range and allows simultaneous protein-expression and growth measurements to define protein-specific expression–fitness relationships.

We applied this approach to approximately 80 proteins encoded on chromosome I of *S. cerevisiae*, integrating expression–fitness relationships with sequence properties, subcellular localization, and cellular morphology. Nearly all proteins could be driven to levels that caused severe growth inhibition or reduced viability, but the abundance at which growth was impaired differed widely. We established IE50, the protein expression level associated with a 50% reduction in growth rate, as a common quantitative measure of overexpression tolerance. This measure revealed associations with predicted structural order, cytoplasmic localization propensity, and sulphur content.

Single-cell imaging linked high tolerance to proteins remaining broadly cytoplasmic, whereas proteins accumulating in particular cellular structures or classified as aggregation-positive tended to have lower limits. Increasing abundance also altered organelle morphology, and extreme overexpression of Fun12, Nup60, and Pex22 generated distinct large-scale intracellular states that depended on specific sequence regions. Together, these measurements allow us to investigate what limits protein abundance and which intracellular structures and states a protein sequence can generate at high abundance.

## Results

### gTOW-mediated high-copy selection enables lethal protein overexpression

We first examined whether gTOW-mediated high-copy selection could drive diverse proteins to expression levels beyond those attainable using conventional multicopy expression. Under standard multicopy conditions (–Ura), overexpression of a toxic protein is expected to select for cells carrying fewer plasmid copies, thereby reducing its expression level (**Figure 1A**). In contrast, under gTOW conditions (–LeuUra), selection for the *leu2-89* marker counteracts this reduction in plasmid copy number and forces the target gene toward higher copy numbers (**Figure 1B**; **Figure S1C**).

**Figure 1.**
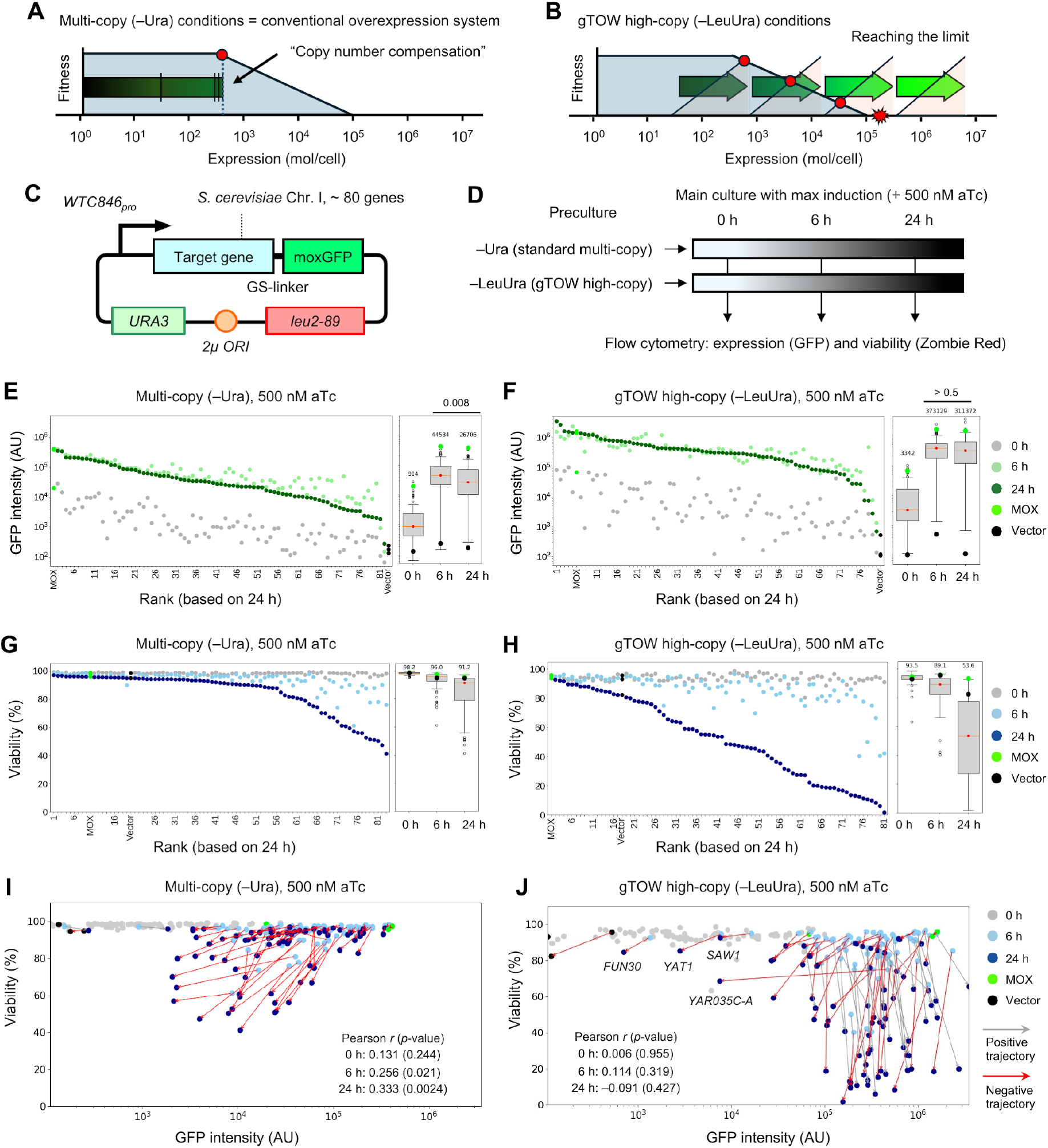
gTOW-mediated high-copy selection suppresses copy-number compensation and enables lethal overexpression. **(A)** Schematic of copy-number compensation under standard multicopy conditions (–Ura). **(B)** Schematic of gTOW-mediated high-copy selection under –LeuUra conditions. The star indicates the expression limit. **(C)** Expression plasmid carrying the WTC846pro-driven target protein–moxGFP fusion cassette, a 2µ origin, *URA3*, and *leu2-89*. **(D)** Flow-cytometry workflow. GFP fluorescence and Zombie Red staining report protein expression and cell viability, respectively, at 0, 6, and 24 h after induction with 500 nM aTc. Individual profiles are shown in **Figure S2**. **(E, F)** Mean live-cell GFP fluorescence (AU) under –Ura **(E)** and –LeuUra **(F)** conditions. Proteins are ranked by fluorescence at 24 h. Points represent the time points in **(D)**; boxplots summarize the distributions across target proteins. MOX, moxGFP alone; Vector, empty vector. The 6- and 24-h measurements were compared using a Wilcoxon signed-rank test. Individual fluorescence distributions are shown in **Figures S3 and S4**, respectively. **(G, H)** Cell viability (%) under –Ura **(G)** and –LeuUra **(H)** conditions. Strains are ranked by viability at 24 h. Time points and controls are as in **(E, F)**; boxplots summarize the distributions across strains. Individual Zombie Red fluorescence distributions are shown in **Figures S5 and S6**, respectively. **(I, J)** Cell viability plotted against mean live-cell GFP fluorescence (AU, logarithmic scale) under –Ura **(I)** and –LeuUra **(J)** conditions. Points represent the time points in **(D)**. Arrows connect measurements for the same target protein from 6 to 24 h. Positive and negative trajectories indicate increases and decreases in mean live-cell GFP fluorescence, respectively, over this interval. Correlations were assessed using Pearson correlation. *FUN30*, *YAR035C-A*, *YAT1*, and *SAW1* are labeled in **(J)**. Changes between 6 and 24 h are compared in **Figures S8A and S8B**, respectively.

We analyzed approximately 80 proteins encoded on chromosome I of *S. cerevisiae* (**Figure 1C**). Each protein was fused to moxGFP, a GFP variant, through a GS linker, allowing its expression to be monitored by GFP fluorescence. Cells were precultured under –Ura (standard multicopy) or –LeuUra (gTOW high-copy) conditions and then transferred to the corresponding medium containing 500 nM aTc. GFP fluorescence and Zombie Red staining were measured by flow cytometry before induction (0 h) and at 6 and 24 h after induction to quantify protein expression in live cells and population viability, respectively (**Figure 1D**; **Figure S2**).

Protein expression was markedly higher under –LeuUra conditions than under –Ura conditions (**Figure 1E, F**; **Figures S3 and S4**). At 6 h, the median fluorescence across the proteins increased from 44,600 AU under –Ura conditions to 373,000 AU under –LeuUra conditions, an approximately 8.4-fold increase. At 24 h, the corresponding values were 26,700 and 311,000 AU, respectively, an approximately 12-fold difference. Under –Ura conditions, expression of many proteins decreased between 6 and 24 h (**Figure 1E**; **Figure S3**). Nevertheless, expression levels were generally positively correlated among the different conditions (*r* > 0.58; **Figure S7A**), indicating that relative differences in protein expressibility were partially preserved.

The increase in protein expression under –LeuUra conditions was accompanied by a substantial loss of cell viability (**Figure 1G, H**; **Figures S5 and S6**). At 24 h, median viability was 91.2% under –Ura conditions but only 53.6% under –LeuUra conditions. Thus, gTOW-mediated high-copy selection pushed the expression of many proteins into a range associated with substantial loss of viability.

We next examined how protein expression and viability changed between 6 and 24 h (**Figure 1I, J**; **Figure S8**). Red arrows indicate decreased mean live-cell GFP fluorescence from 6 to 24 h, whereas gray arrows indicate increased or unchanged fluorescence. Under –Ura conditions, many proteins showed decreases in both expression and viability (**Figure 1I**; **Figure S8A**). This behavior is consistent with copy-number compensation: cells with the highest expression preferentially lose viability, enriching cells with lower plasmid copy numbers and reducing population-level expression. In contrast, under –LeuUra conditions, high expression was maintained for many proteins despite decreasing viability (**Figure 1J**; **Figure S8B**). At 24 h, expression level and viability were not significantly correlated under –LeuUra conditions (*r* = –0.091, *p* = 0.427), indicating that differences in cell death could not simply be explained by differences in the amount of protein expressed.

Four proteins—Fun30, Yar035c-a, Yat1, and Saw1—remained at relatively low expression levels even under maximal induction in –LeuUra medium (**Figure 1J**; **Figures S2 and S4**). Yar035c-a was expressed at 6 h but decreased markedly by 24 h (**Figure S4**), indicating that its abundance was not maintained during prolonged induction.

Excluding these four proteins, all tested proteins reached fluorescence levels corresponding to at least approximately 2% of that of MOX under maximal induction in –LeuUra medium (**Figure S4**). Using the reported moxGFP abundance of approximately 15% of total cellular protein as a reference (Fujita et al., 2025), this corresponds to a rough MOX-equivalent estimate of at least approximately 0.3% of total cellular protein, assuming comparable fluorescence per GFP molecule and without correcting for fusion-protein molecular weight.

Together, these results demonstrate that gTOW-mediated high-copy selection counteracts the reduction in protein expression associated with conventional multicopy overexpression and allows diverse proteins to be pushed into expression regimes associated with severe loss of viability.

### Quantifying protein toxicity by expression–fitness relationships

To quantify the effects of target-protein overexpression on growth, we examined the relationship between protein expression and growth rate. Cells were precultured under –Ura conditions and then transferred to either –Ura or –LeuUra medium, in which expression was induced with 0, 50, 150, or 500 nM aTc. OD595 and GFP fluorescence were continuously monitored using a plate reader, and the resulting time-course data were used to calculate the maximum growth rate (MGR) and maximum fluorescence level (MFL), respectively (**Figure 2A**; **Figures S9 and S11**).

**Figure 2.**
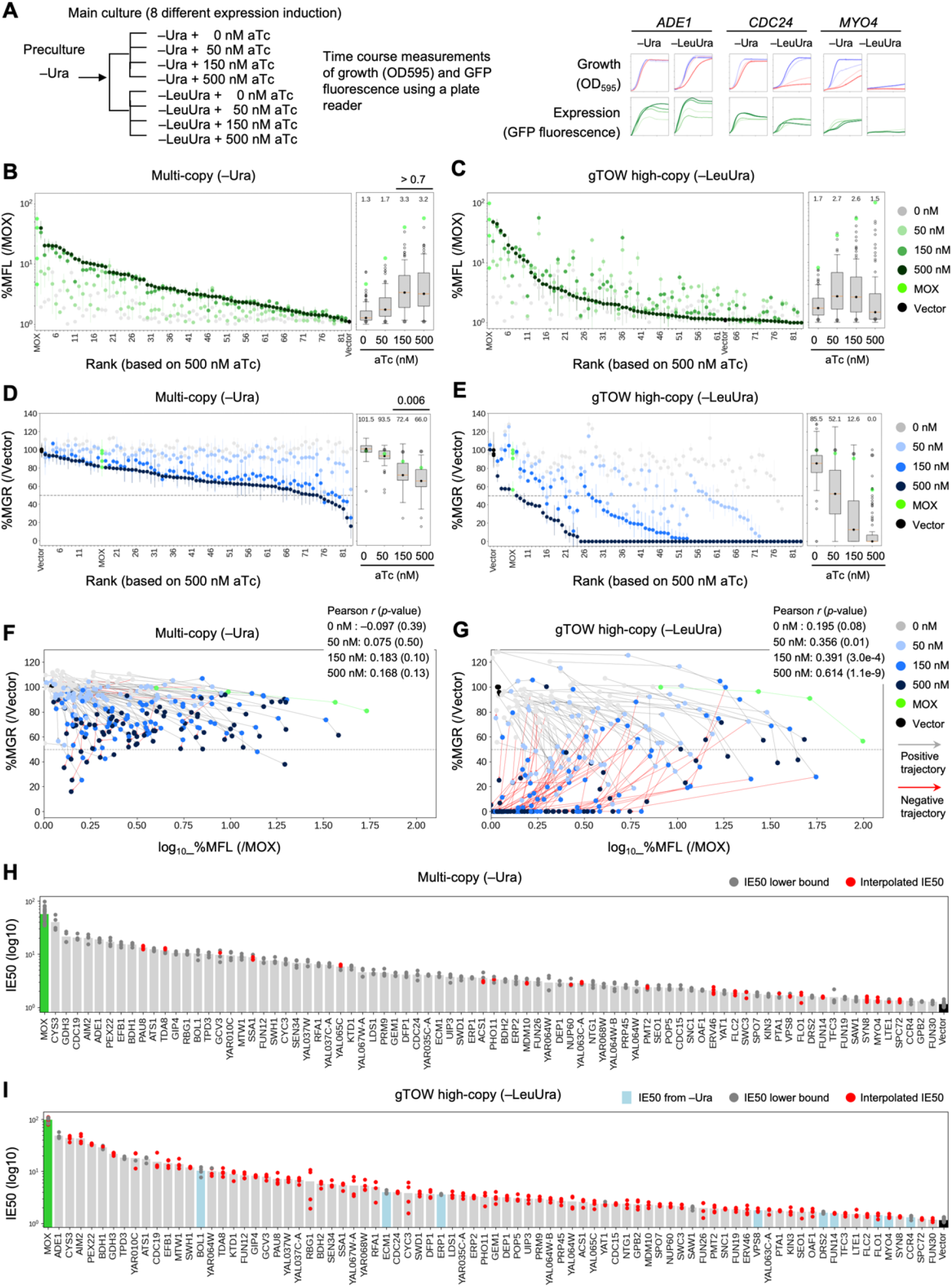
Quantification of protein toxicity based on expression–fitness relationships. **(A)** Plate-reader workflow and representative growth and GFP fluorescence time courses for strains overexpressing *ADE1*, *CDC24*, and *MYO4*. Measurements were obtained under –Ura and –LeuUra conditions with 0, 50, 150, or 500 nM aTc. MGR, maximum growth rate; MFL, maximum fluorescence level. Complete time courses are shown in **Figures S9 and S11**. **(B, C)** MFL expressed as a percentage of the MOX control (%MFL) under –Ura **(B)** and –LeuUra **(C)** conditions. Proteins are ranked by %MFL at 500 nM aTc. Boxplots summarize the distributions across target proteins at each aTc concentration in **(A)**. MOX, moxGFP alone; Vector, empty vector. Comparisons among induction conditions are shown in **Figure S10**. **(D, E)** MGR expressed as a percentage of the Vector control (%MGR) under –Ura **(D)** and –LeuUra **(E)** conditions. Induction conditions and controls are as in **(B, C)**; boxplots summarize the distributions across target proteins. The dashed line marks %MGR = 50%. Comparisons among induction conditions are shown in **Figure S12**. **(F, G)** Expression–growth trajectories under –Ura **(F)** and –LeuUra **(G)** conditions. Each point represents log10 %MFL and %MGR for one target protein at one aTc concentration. Arrows connect measurements for the same protein in order of increasing aTc concentration (0, 50, 150, and 500 nM). Positive and negative trajectories indicate increases and decreases in %MFL, respectively, between successive induction levels. Correlations were assessed using Pearson correlation. Individual protein trajectories are shown in **Figure S13**. **(H, I)** IE50 values of individual target proteins under standard multicopy conditions **(H)** and gTOW high-copy conditions **(I)**. “Interpolated IE50” denotes estimates obtained by interpolation, “IE50 lower bound” denotes the maximum observed MFL when growth inhibition did not reach 50%, and “IE50 from –Ura” denotes –Ura estimates used when an estimate could not be obtained under –LeuUra conditions. MOX and Vector are shown as controls. The *y*-axis is logarithmic. Values under the two conditions are compared in **Figure S14**.

Normalized MFL values under –Ura and –LeuUra conditions are shown in **Figure 2B, C**. Under –Ura conditions, MFL increased with increasing aTc concentration but approached saturation at approximately 150 nM, with no significant difference between 150 and 500 nM aTc (**Figure 2B**, *p* > 0.7). Under –LeuUra conditions, MFL initially increased with induction but the measured MFL values decreased at 150 and 500 nM aTc (**Figure 2C**; **Figure S9**). Comparison of protein-expression ranks across conditions revealed positive correlations between most conditions, whereas the correlation was almost lost between 0 and 500 nM aTc under –LeuUra conditions (*r* = 0.10; **Figure S10**).

We next compared relative MGR values across induction conditions (**Figure 2D, E**; **Figure S12**). Under –Ura conditions, MGR progressively decreased with increasing aTc concentration, with a significant further decrease from 150 to 500 nM (**Figure 2D**, *p* = 0.006). In contrast, under –LeuUra conditions, the number of strains classified as non-growing increased sharply with induction strength. At 500 nM aTc, more than 90% of the strains were classified as non-growing based on a final OD595 of ≤0.5 (**Figure 2E**; **Figure S11**). For these cultures, MGR was set to zero and the observed MFL value was retained. The decrease in measured MFL at 150 and 500 nM aTc under –LeuUra conditions coincided with the increasing prevalence of low-density cultures, for which plate-reader fluorescence could not be quantified reliably (**Figure 2C**). MGR values were more strongly correlated between conditions with similar induction strengths, whereas the correlations decreased as the difference in induction strength increased (**Figure S12**).

For each protein, we plotted MFL against MGR across the four induction levels to visualize its expression–fitness relationship (**Figure 2F, G**; **Figure S13**). Under –Ura conditions, increasing induction generally increased MFL and reduced MGR, but the correlation between MFL and MGR at each induction level remained weak. The Pearson correlation coefficients at 150 and 500 nM aTc were *r* = 0.183 and 0.168, respectively. Moreover, only approximately 10% of the target proteins showed a reduction in MGR to 50% or less (**Figure 2F**). Under –LeuUra conditions, increasing induction substantially reduced MGR for many proteins, with more than 90% of the strains classified as non-growing at 500 nM aTc (**Figure 2G**; **Figure S13**).

These measurements allowed us to quantify the expression level at which each protein substantially impaired growth. We therefore defined the 50% inhibitory expression level (IE50) as the protein expression level at which MGR decreased to 50% of the vector-control growth rate. When MGR crossed the 50% threshold with increasing induction, IE50 was estimated by linear interpolation of log10(MFL) between the two points spanning the threshold. When MGR did not fall below 50%, the highest observed MFL was used as a lower-bound estimate of IE50. When an IE50 estimate could not be obtained under –LeuUra conditions, the corresponding –Ura estimate was used as a fallback, if available. This procedure provided a common metric for comparing the amount of each protein that could accumulate before growth was severely compromised (**Figure 2H, I**).

IE50 values obtained under –Ura and –LeuUra conditions were strongly correlated (*r* = 0.93; **Figure S14A**), indicating that the relative ordering of proteins was largely preserved between the two conditions. However, proteins with higher IE50 values tended to show larger –LeuUra/–Ura ratios (*r* = 0.47; **Figure S14B**). This relationship is consistent with copy-number compensation limiting the expression levels attainable under –Ura conditions (**Figure 1A**), whereas gTOW high-copy selection extends the measurable expression range (**Figure 1B**).

Comparison of the correlation structure among expression, growth, viability, and IE50 measurements showed that IE50 was strongly associated with expression-related measurements but exhibited a distinct relationship with growth and viability (**Figure S15**). IE50 was also significantly correlated with GFP fluorescence measured by flow cytometry, although this relationship was stronger under standard multicopy conditions (–Ura; Pearson *r* = 0.782, *R*² = 0.611) than under gTOW high-copy conditions (–LeuUra; *r* = 0.532, *R*² = 0.283; **Figure S16**). Thus, IE50 measured under standard multicopy conditions is strongly influenced by protein expressibility, whereas gTOW high-copy selection reveals larger differences among proteins that cannot be explained by expressibility alone. We therefore next investigated the protein properties underlying these differences.

### Subcellular localization, structural order, and sulphur content constrain protein overexpression

We next used IE50 to identify protein properties associated with tolerance to overexpression. We analyzed approximately 70 parameters encompassing gene-expression-related features, amino-acid composition, physicochemical properties, subcellular localization, and predicted structural properties. Some of these parameters were correlated with one another (**Figure S17A, B**).

We compared Pearson and Spearman correlations between each parameter and IE50 under standard multicopy (–Ura) and gTOW high-copy (–LeuUra) conditions (**Figure 3A**; **Figure S17C, D**). Overall, the correlation coefficients were well correlated between the two conditions (Pearson: *r* = 0.78; Spearman: *r* = 0.62), although some parameters showed condition-specific associations. At an FDR threshold of 0.05, 10 and 4 parameters showed significant Pearson correlations under –Ura and –LeuUra conditions, respectively. For Spearman correlations, 17 and 18 parameters were significant under –Ura and –LeuUra conditions, respectively (**Figure S17C, D**).

**Figure 3.**
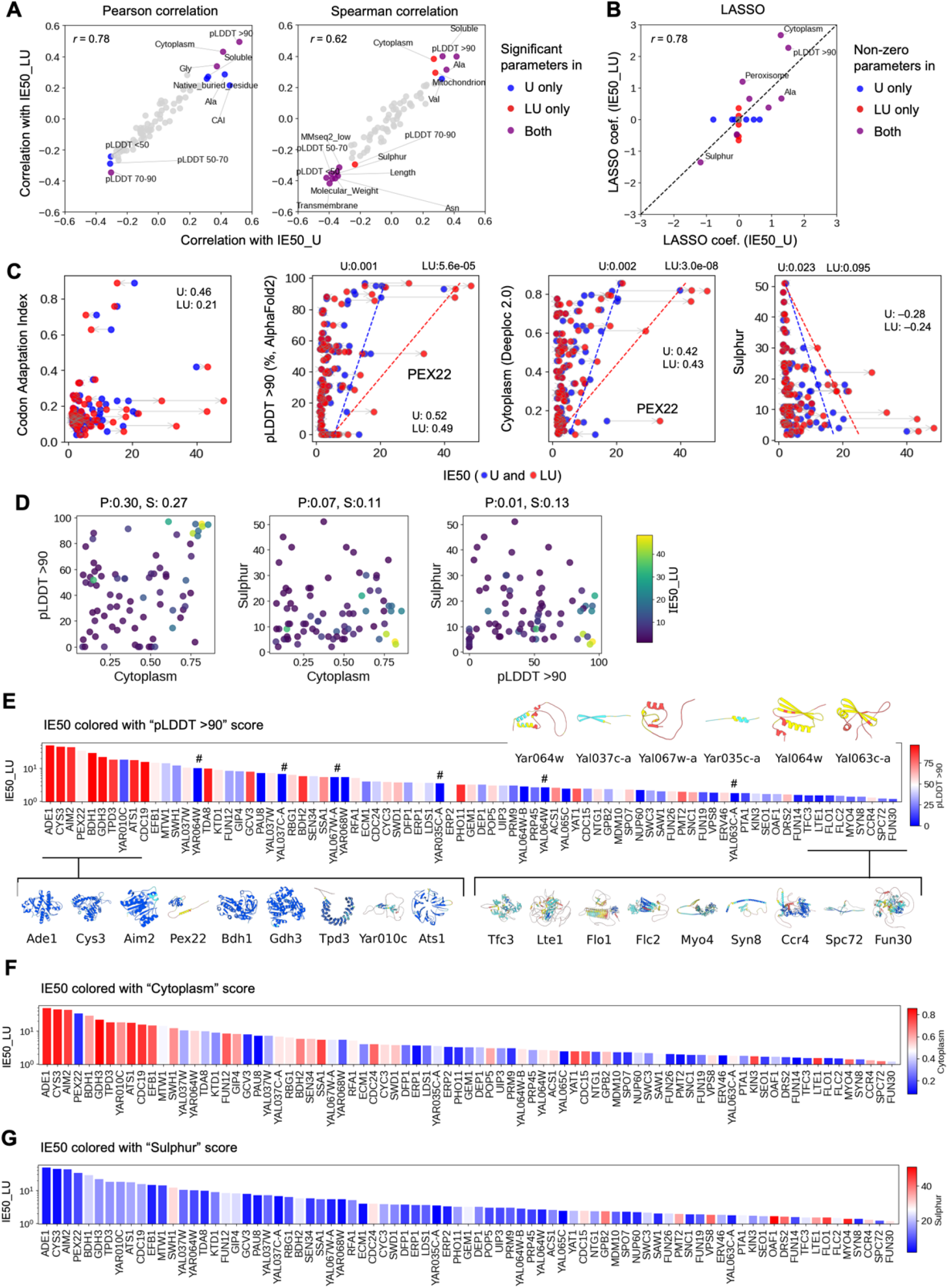
Subcellular localization, structural order, and sulphur content constrain protein overexpression limits. **(A)** Correlation coefficients between protein parameters and IE50 under –Ura (*x*-axis) and –LeuUra (*y*-axis) conditions: Pearson (left) and Spearman (right). Parameter classes indicate significance under –Ura only, –LeuUra only, or both conditions (Benjamini–Hochberg-adjusted FDR < 0.05). Complete correlation results are shown in **Figure S17**. **(B)** LASSO regression coefficients under –Ura (*x*-axis) and –LeuUra (*y*-axis) conditions. Parameter classes indicate non-zero coefficients under –Ura only, –LeuUra only, or both conditions. The dashed line marks *y* = *x*. The three-parameter regression model and residual analyses are shown in **Figure S18**. **(C)** IE50 plotted against representative protein parameters. Lines connect values for the same protein under –Ura and –LeuUra conditions. Sulphur content denotes the total number of cysteine and methionine residues per protein. For pLDDT > 90, Cytoplasm score, and sulphur content, dashed lines show the fitted conditional 90th percentile of IE50; *p* values refer to the quantile-regression slopes. *PEX22* is labeled. Relationships between IE50 and all analyzed protein parameters are shown in **Figure S19**. **(D)** Pairwise relationships among pLDDT > 90, Cytoplasm score, and sulphur content. Each point represents a target protein, with its IE50 under –LeuUra conditions also indicated. P, Pearson; S, Spearman. Correlations with other experimental measurements are shown in **Figure S20**. **(E–G)** Target proteins are ordered by decreasing IE50 under –LeuUra conditions in all three panels. The panels show the fractions of residues with AlphaFold2 pLDDT > 90 (E), DeepLoc 2.0 Cytoplasm scores (F), and sulphur contents (G). Representative predicted structures are shown in **(E)**; # marks emerging genes. Predicted structures of all target proteins are shown in **Figure S21**. Detailed relationships between protein parameters and GFP fluorescence, viability, MGR, and MFL are shown in **Figures S22–S25**, respectively.

To identify major predictors of IE50 while accounting for redundancy among parameters, We performed least absolute shrinkage and selection operator (LASSO) regression (**Figure 3B**). Nine parameters had non-zero coefficients under both conditions, whereas six were specific to –Ura and six to –LeuUra conditions. Among the strongest associations, the fraction of residues with AlphaFold2 pLDDT > 90 (indicating very high local prediction confidence) and the Cytoplasm score predicted by DeepLoc 2.0 were positively associated with IE50, whereas sulphur content was negatively associated with IE50. Sulphur content was strongly correlated with protein length and molecular weight and may therefore partly reflect these properties, although it retained a large coefficient in the LASSO analysis.

A regression model incorporating Cytoplasm score, the fraction of residues with pLDDT > 90, and sulphur content explained approximately 40–50% of the variation in IE50 (–Ura: *R*² = 0.46; –LeuUra: *R*² = 0.41; **Figure S18A, B**). Analysis of the residuals from this model identified no additional parameters with significant Pearson or Spearman correlations with the residuals (**Figure S18C, D**). Together, these results support Cytoplasm score, the fraction of residues with pLDDT > 90, and sulphur content as major predictors of IE50 among the protein parameters examined, although substantial variation remains unexplained.

Representative relationships are shown in **Figure 3C**. The codon adaptation index (CAI) was significantly associated with IE50 under –Ura conditions, but this relationship became weaker under –LeuUra conditions. The fraction of residues with pLDDT > 90 was positively correlated with IE50 under both conditions (–Ura: Pearson *r* = 0.52; –LeuUra: *r* = 0.49). Cytoplasm score and sulphur content also showed positive and negative Spearman correlations, respectively, with IE50 under –LeuUra conditions (**Figure 3A**).

Notably, the expansion of the IE50 distribution from –Ura to –LeuUra conditions was concentrated in particular regions of these parameter spaces (**Figure 3C**). Quantile regression at the 0.9 quantile showed that the upper boundary of IE50 under –LeuUra conditions increased significantly with increasing Cytoplasm score and fraction of residues with pLDDT > 90 (Cytoplasm: *p* = 3.0 × 10⁻⁸; pLDDT > 90: *p* = 5.6 × 10⁻⁵). Sulphur content showed the opposite trend, although it did not reach statistical significance (*p* = 0.095). These results suggest that these properties are associated not only with average IE50 but also with the upper range of protein expression that cells can tolerate.

Among the three parameters, the fraction of residues with pLDDT > 90 and Cytoplasm score showed a weak positive correlation (Pearson *r* = 0.30; Spearman *ρ* = 0.27), whereas no clear correlations were observed between the other pairs (**Figure 3D**).

When proteins were ordered by decreasing IE50 under –LeuUra conditions, proteins with a high fraction of residues with pLDDT > 90 tended to have higher IE50 values (**Figure 3E**). However, a subset of proteins exhibited moderate IE50 despite having low fractions of residues with pLDDT > 90; this group included proteins encoded by evolutionarily young genes, potentially arising *de novo* from ancestral noncoding sequences (Vakirlis et al., 2020; **Figure 3E**, #). AlphaFold2-predicted structures of all target proteins, including these proteins, are shown in **Figure S21**. Similarly, proteins with high IE50 frequently had high Cytoplasm scores (**Figure 3F**). In contrast, proteins with high sulphur content were largely absent from the high-IE50 range (**Figure 3G**).

We further examined how the protein properties associated with IE50 were related to individual experimental measurements. Comprehensive comparison of approximately 70 protein parameters with expression levels, growth rates, and viability measured by flow cytometry and plate-reader assays showed that the strength and significance of these associations varied with both the measurement and expression condition (**Figure S20**). Analyses of GFP fluorescence, viability, MGR, and MFL across individual induction conditions likewise showed that associations with multiple protein properties changed with expression conditions (**Figures S22–S25**). Thus, the relationships with IE50 identified in **Figure 3** do not appear to arise from a single expression condition or experimental readout, but instead reflect how protein abundance and its cellular consequences change across the expression range.

Pex22 was a notable exception to the general trends. Despite its high IE50, Pex22 had both a low fraction of residues with pLDDT > 90 and a low Cytoplasm score (**Figure 3C**). Thus, Pex22 represents a protein whose high overexpression tolerance is not readily explained by the major properties identified above. The intracellular structures formed by Pex22 at high abundance are examined in a later section.

Together, these analyses identify cytoplasmic localization, predicted structural order, and sulphur-related protein properties as major features associated with the amount of protein that cells can tolerate.

### Subcellular localization is associated with protein expression limits and organelle morphology during overexpression

Our results thus far suggested that proteins localized to the cytoplasm, or those with a low propensity to localize to specific intracellular structures, may be tolerated at higher expression levels. We therefore directly examined how subcellular localization changes as protein expression increases.

Protein expression was induced with four aTc concentrations (0, 50, 150, and 500 nM) under standard multicopy (–Ura) and gTOW high-copy (–LeuUra) conditions, and the localization of GFP-fused proteins was examined after 6 h by high-content imaging (**Figure 4A**; **Figure S26**). Individual cells were segmented from the acquired images, and approximately 5 × 10⁵ single-cell GFP images were subjected to machine-learning-based localization classification (**Figure S26**). The classifier was trained using approximately 100,000 labeled images and assigned cells to eight localization classes: Cytoplasm, Aggregate, Nucleus, ER, Mitochondria, Nuclear membrane, Puncta/Vesicle, and Plasma membrane (**Figure 4B**; **Figures S26 and S27**). Images failing quality-control criteria, including poor image quality and insufficient fluorescence, were excluded (**Figure S27**). The classifier showed high classification accuracy across the localization classes (**Figure S28A**).

**Figure 4.**
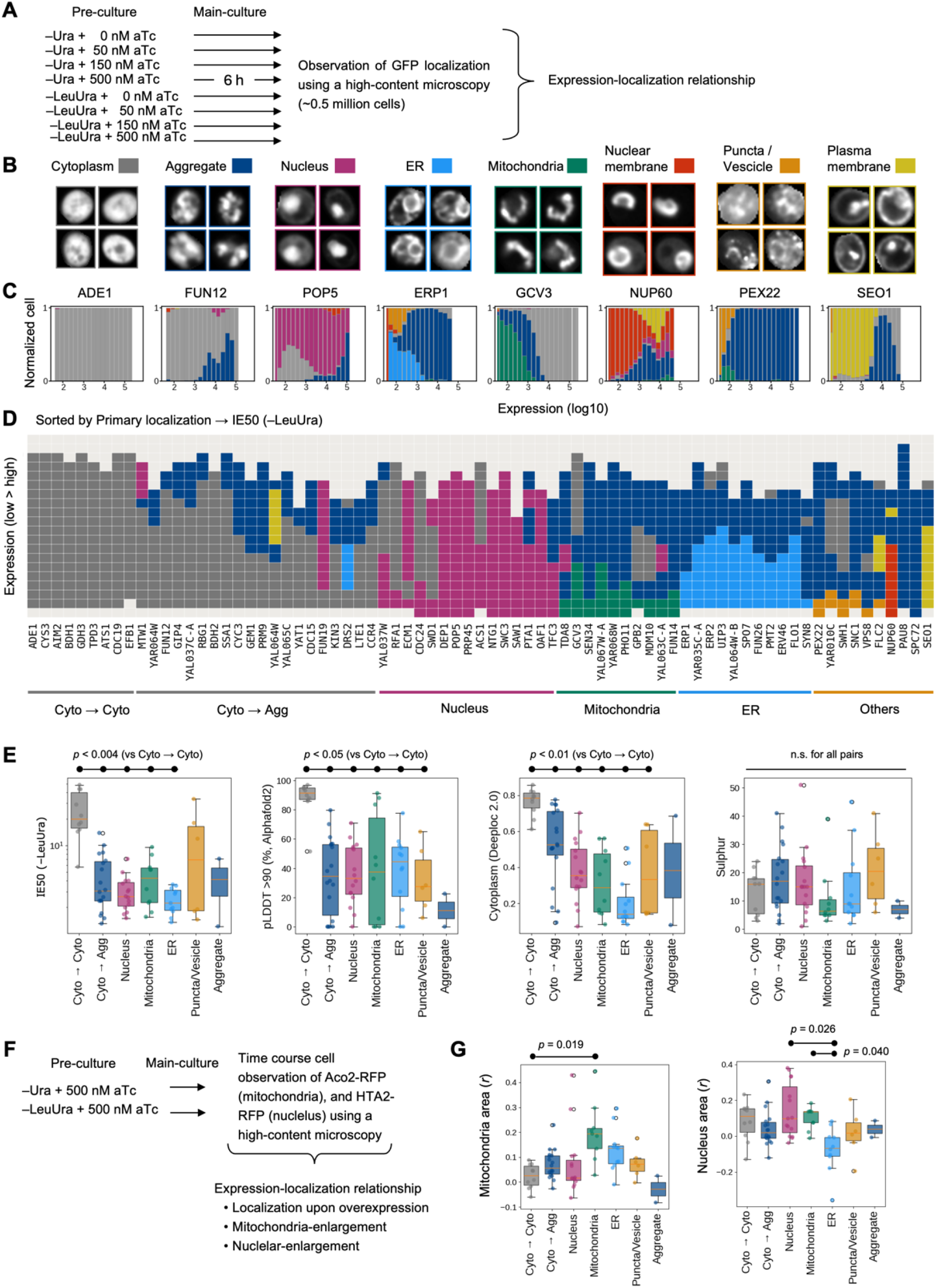
Subcellular localization is associated with protein expression limits and organelle morphology during overexpression. **(A)** High-content imaging workflow for localization analysis after 6 h of culture under –Ura or –LeuUra conditions with 0, 50, 150, or 500 nM aTc. The full workflow is shown in **Figure S26**. **(B)** Representative images of the eight localization classes. Additional examples and excluded quality-control classes are shown in **Figure S27**; classifier performance is shown in **Figure S28A**. **(C)** Fractions of cells assigned to each localization class across 20 log10 GFP fluorescence bins for representative proteins. Classes are defined in **(B)**. Distributions for all target proteins are shown in **Figure S29**. **(D)** Dominant localization class in each GFP fluorescence bin for all target proteins. Columns represent proteins, grouped by low-expression localization and ordered within each group by IE50 under –LeuUra conditions. Major localization-transition classes are indicated below. Individual distributions and corresponding single-cell images are shown in **Figures S29 and S30**, respectively. **(E)** IE50 under –LeuUra conditions, fraction of residues with pLDDT > 90, DeepLoc 2.0 Cytoplasm score, and sulphur content, grouped by the localization patterns in **(D)**. Pairwise comparisons were performed using two-sided Mann–Whitney U tests with Holm correction across the 15 comparisons within each parameter. The Aggregate group was excluded from testing because it contained only two proteins. **(F)** Organelle-morphology imaging workflow under –Ura or –LeuUra conditions with 500 nM aTc. Aco2-RFP and Hta2-RFP label mitochondria and nuclei, respectively. Morphological-feature correlations are shown in **Figure S31**; individual images and measurements are shown in **Figures S32 and S33**. **(G)** Pearson correlation coefficients between single-cell GFP fluorescence intensity and mean mitochondrial area (left) or nuclear area (right), grouped by expression-dependent localization pattern. Each point represents one target protein. *p* values were calculated using two-sided Mann–Whitney U tests with Holm correction across the 15 pairwise comparisons within each panel. The Aggregate group was excluded from testing because it contained only two proteins. Individual measurements are shown in **Figures S32 and S33**.

We next examined how protein localization changed with expression level by grouping individual cells into 20 equal-width bins of log10-transformed GFP fluorescence (**Figure 4C**; **Figure S29**). Ade1 remained predominantly cytoplasmic across a broad expression range, whereas Fun12 was increasingly assigned to the Aggregate class at higher expression levels. Pop5 retained its nuclear localization even at high expression, whereas Gcv3 shifted from mitochondrial localization toward aggregation. We summarized these expression-dependent localization profiles across all target proteins, ordered by their predominant localization at low expression and IE50 under –LeuUra conditions (**Figure 4D**; **Figure S30**).

Two comparisons supported the interpretation of these imaging data. First, experimentally observed localization at low expression generally agreed with the highest-scoring compartment predicted by DeepLoc 2.0 (**Figure S28B**). Second, maximum GFP fluorescence measured by imaging correlated strongly with flow-cytometry measurements at 6 h after induction with 500 nM aTc, IE50 under –LeuUra conditions, and plate-reader MFL at 500 nM aTc (Spearman’s *ρ* = 0.824, 0.746, and 0.781, respectively; **Figure S28C**).

Localization changes during overexpression differed markedly according to low-expression localization (**Figure 4D**; **Figures S29 and S30**). Cytoplasmic proteins generally either remained cytoplasmic or shifted toward aggregation as expression increased. Nuclear proteins tended to retain nuclear localization, whereas mitochondrial and ER proteins frequently showed aggregation at high expression levels. Consistent with this pattern, the occurrence of aggregation was significantly associated with low-expression localization (*χ*² test, *p* = 7.9 × 10⁻⁶).

We next compared these localization patterns with protein expression limits (**Figure 4E**). Proteins that remained cytoplasmic (Cyto→Cyto) had significantly higher IE50 values under –LeuUra conditions than proteins in the Cyto→Agg, Nucleus, Mitochondria, and ER groups (Holm-adjusted *p* < 0.004 for all four comparisons). The Cyto→Cyto group also had higher fractions of residues with pLDDT > 90 and higher Cytoplasm scores than these four groups and the Puncta/Vesicle group, whereas sulphur content did not differ significantly among the tested groups (**Figure 4E**). These findings link experimentally observed cytoplasmic localization to higher expression tolerance, consistent with the association between Cytoplasm score and IE50 identified in **Figure 3**.

Proteins for which Aggregate became the dominant localization class in at least one expression bin during overexpression had lower IE50 values than proteins for which this transition was not observed (Wilcoxon rank-sum test, *p* = 0.0084). The probability of aggregation was significantly associated with IE50 (logistic regression, *β* = −0.088, *p* = 0.009). In contrast, the expression level at which aggregation was first observed was not significantly correlated with IE50 (Spearman’s *ρ* = 0.16, *p* = 0.255). Together, these results indicate that proteins classified as aggregation-positive tend to have lower IE50 values, while the relationship between aggregation onset and expression limits remains unresolved.

We further examined whether overexpression affected the morphology of the intracellular structures to which proteins localize. Mitochondria and nuclei were labeled with Aco2-RFP and Hta2-RFP, respectively, and their morphology was monitored over time by high-content imaging following induction with 500 nM aTc under standard multicopy (–Ura) and gTOW high-copy (–LeuUra) conditions (**Figure 4F**). Multiple morphological features were extracted from mitochondrial and nuclear images. Because many area- and shape-related features were strongly correlated, Mean_Mito_AreaShape_Area and Mean_Nuc_AreaShape_Area were used as representative measures of mitochondrial and nuclear size, respectively (**Figure S31**).

Analysis of cells overexpressing individual target proteins revealed substantial protein-dependent variation in mitochondrial area and number and in nuclear area and compactness (**Figures S32 and S33**). For each target protein, we calculated the Pearson correlation between single-cell GFP fluorescence intensity and mean mitochondrial radius or nuclear area and compared these coefficients across localization groups. Correlations with mitochondrial radius were more positive in the Mitochondria group than in the Cyto→Cyto group (Holm-adjusted *p* = 0.019). In the ER group, correlations between GFP fluorescence intensity and nuclear area tended to be negative, indicating that higher expression was associated with smaller nuclear area. These correlations were significantly lower than those in the Nucleus and Mitochondria groups (Holm-adjusted *p* = 0.026 and 0.040, respectively; **Figure 4G**). These results indicate that increasing protein expression is associated not only with changes in protein localization but also with organelle morphology. The association between ER-localized protein expression and nuclear area further suggests that these effects may extend beyond the organelle of localization, potentially reflecting functional coupling between organelles.

Together, these results show that protein expression limits are closely associated with subcellular localization. Proteins that localize to specific intracellular structures or tend to aggregate during overexpression generally have lower expression limits, whereas proteins that remain cytoplasmic tolerate substantially higher expression levels.

### Protein overexpression drives cells into previously unexplored intracellular states

Finally, we examined the intracellular structures that emerged under strong protein overexpression. Fluorescence patterns ranged from broadly distributed signals to discrete foci, large intracellular bodies, and ring-like patterns (**Figure 5A**; **Figure S34**). We selected Fun12, Nup60, and Pex22 for further characterization and mapped the protein regions contributing to their distinct phenotypes.

**Figure 5.**
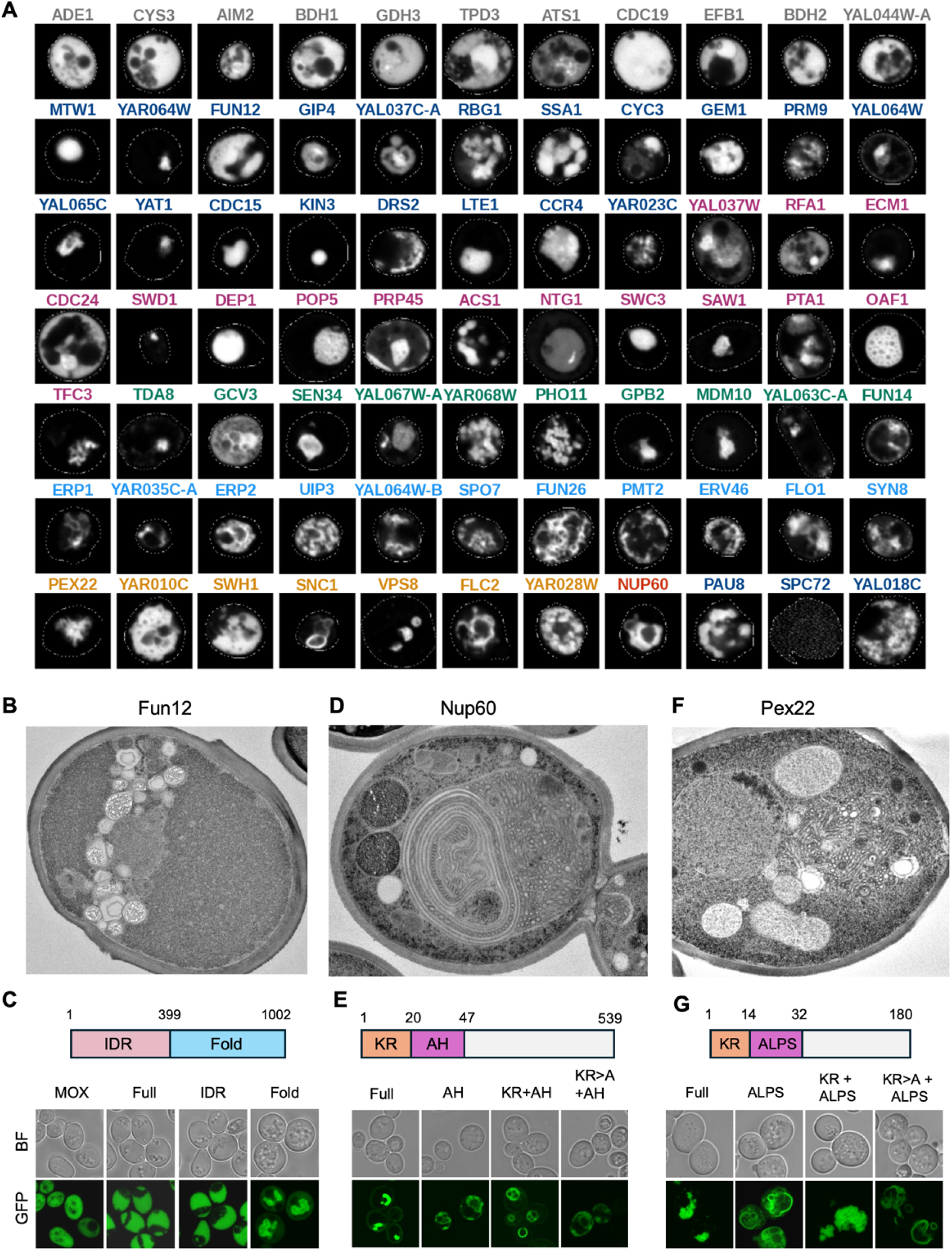
Strong protein overexpression induces distinct intracellular states driven by specific protein regions. **(A)** Representative GFP fluorescence images of target proteins under strong overexpression conditions. Proteins are arranged according to their major localization patterns, using the localization classes defined in Figure 4. Additional representative images of all target proteins are shown in **Figure S34**. **(B)** Transmission electron microscopy (TEM) image of a cell overexpressing Fun12. Additional fluorescence and TEM images are shown in **Figure S35**. TEM images in **(B)**, **(D)**, and **(F)** are displayed at individually adjusted magnifications and are not shown at a common scale. **(C)** Fun12 domain organization and representative bright-field (BF) and GFP fluorescence images of cells overexpressing full-length Fun12 or the indicated fragments. IDR, intrinsically disordered region (residues 1–399); folded region, residues 400–1002. MOX is the control. FRAP and IE50 measurements are shown in **Figure S35**. **(D)** TEM image of a cell overexpressing Nup60. Additional images and serial sections are shown in **Figure S36**. **(E)** Nup60 domain organization and representative BF and GFP fluorescence images of cells overexpressing full-length Nup60 or the indicated fragments and mutants. KR, Lys/Arg-rich region (residues 1–20); AH, amphipathic helix (residues 27–47); KR>A, replacement of Lys and Arg in the KR-rich region with alanine. Additional localization images and IE50 measurements are shown in **Figure S36**. **(F)** TEM image of a cell overexpressing Pex22. Additional images are shown in **Figure S37**. **(G)** Pex22 domain organization and representative BF and GFP fluorescence images of cells overexpressing full-length Pex22 or the indicated fragments and mutants. KR, Lys/Arg-rich region (residues 1–14); ALPS, putative amphipathic lipid packing sensor motif (residues 15–32). KR>A is defined in **(E)**. Additional localization images and IE50 measurements are shown in **Figure S37**.

Although Fun12 was assigned to the Aggregate class by high-content imaging (**Figure 4C**), confocal microscopy revealed broadly distributed, relatively homogeneous cytoplasmic fluorescence rather than discrete aggregates. Organelles were displaced toward one side of the cell, forming a densely clustered, band-like region (**Figure 5A, C**; **Figure S35A**). TEM likewise showed asymmetric organelle clustering and a relatively homogeneous region occupying much of the remaining cytoplasm (**Figure 5B**; **Figure S35B, C**). Overexpression of the N-terminal, charged-residue-rich intrinsically disordered region (IDR) reproduced the cytoplasmic phenotype of full-length Fun12, whereas the C-terminal folded region formed discrete aggregates (**Figure 5C**). In fluorescence recovery after photobleaching (FRAP) experiments, full-length Fun12 and the IDR showed slower recovery than MOX, whereas the folded-region aggregates showed little recovery (**Figure S35D, E**). The IDR also had a higher IE50 than full-length Fun12, whereas the folded region had a lower IE50 (**Figure S35F**). These results identify the IDR as sufficient to reproduce the dynamic cytoplasmic state and suggest that it limits the formation of discrete aggregates by full-length Fun12.

Nup60 overexpression produced extensive, closely apposed membranes around the nucleus (**Figure 5D**; **Figure S36A**). Previous work reported nuclear-envelope deformation and tubular membrane extensions upon overexpression of Nup60 or its N-terminal region (Mészáros et al., 2015). Here, serial TEM sections supported the presence of multiple membrane layers rather than cross-sections of a single tube (**Figure S36B**). We next examined the N-terminal amphipathic helix (AH), which mediates membrane binding, and its adjacent lysine- and arginine-rich (KR-rich) region (Cibulka et al., 2022; Mészáros et al., 2015). An AH-containing fragment associated with membranes but did not reproduce the nuclear-envelope localization and membrane structures observed with full-length Nup60. Including the KR-rich region restored these phenotypes, whereas reducing its positive charge altered them (**Figure 5E**; **Figure S36C**). Constructs displaying nuclear-envelope localization and abnormal membrane structures had lower IE50 values than constructs lacking these properties (**Figure S36D**). Thus, the KR-rich region contributes to nuclear-envelope targeting and membrane remodeling, which were associated with lower overexpression tolerance.

Pex22 overexpression produced a different membrane phenotype: TEM revealed densely packed tubular membrane profiles occupying a large fraction of the cell, while fluorescence microscopy showed an extensive Pex22-GFP-positive region containing numerous small signal-depleted spaces (**Figure 5F, G**; **Figure S37A**). A fragment containing the putative amphipathic lipid packing sensor (ALPS) motif associated with membranes but did not efficiently reproduce the full-length phenotype. Inclusion of the adjacent KR-rich region produced structures more similar to those formed by full-length Pex22, whereas replacing its lysine and arginine residues with alanine weakened this phenotype (**Figure 5G**; **Figure S37B**). These results implicate both the putative ALPS motif and the adjacent positively charged region in structure formation. Unlike Nup60, however, Pex22 constructs showed no clear correspondence between membrane morphology and IE50 (**Figure S37C**).

Together, these analyses show that extreme overexpression reveals distinct, sequence-dependent capacities for intracellular organization. The Fun12 IDR supported a dynamic cytoplasmic state, whereas membrane-associated regions and adjacent KR-rich sequences contributed to the structures formed by Nup60 and Pex22. The relationship between these phenotypes and expression tolerance differed among proteins, indicating that large-scale structural remodeling does not uniformly predict growth inhibition.

## Discussion

In this study, we showed that protein overexpression toxicity is better understood as a difference in the abundance at which proteins become harmful than as a distinction between toxic and non-toxic proteins. In previous systematic screens, overexpression-associated growth inhibition had not been reported for more than half of yeast proteins (**Figure S1A, B**). Yet, when expression was pushed sufficiently high, nearly all chromosome I proteins examined here caused severe growth inhibition or reduced viability (**Figure 1**). Our results therefore show that an apparently non-toxic phenotype can reflect insufficient expression rather than an absence of toxicity. To quantify these differences, we established IE50 as a protein-expression counterpart to IC50: whereas IC50 denotes the concentration of a substance that inhibits a specified activity by 50%, IE50 denotes the protein expression level at which growth rate is reduced by 50% (**Figure 2**). This brings the principle that “the dose makes the poison” to protein overexpression: the relevant distinction is how much of each protein a cell can tolerate.

By placing proteins with widely different tolerances on a common continuous scale, IE50 allowed us to identify protein properties associated with toxicity through systematic analysis. Remarkably, analysis of only approximately 80 proteins revealed three major properties associated with overexpression tolerance: predicted structural order, cytoplasmic localization propensity, and sulphur content (**Figure 3**). Earlier large-scale studies had already associated dosage sensitivity with properties such as intrinsic disorder and interaction potential (Vavouri et al., 2009; Makanae et al., 2013; Tomala & Korona, 2013). Our contribution is to connect protein properties to experimentally measured expression thresholds, including for proteins whose toxicity was not apparent within conventional expression ranges. The emergence of these associations from a relatively small, functionally diverse set demonstrates the power of abundance-based measurements to reveal constraints on protein overexpression.

The association with pLDDT provides a new link between predicted protein structure and overexpression tolerance. Several highly tolerated proteins had compact, confidently predicted folds (**Figure 3E**), suggesting that adopting a well-defined structure helps limit toxicity. This interpretation is consistent with the hypothesis that selection against misfolding favours robust protein sequences at high expression levels (Drummond et al., 2005; Geiler-Samerotte et al., 2011). Conversely, several proteins with low IE50 contained extensive low-confidence regions consistent with intrinsic disorder (**Figure 3E**), supporting a role for promiscuous interactions involving disordered regions (Vavouri et al., 2009). Sequence-based disorder predictions, however, were not significantly associated with IE50 in our analysis (**Figure 3**; **Figure S17**). The pLDDT-based measure may therefore capture structural features relevant to toxicity that are not adequately represented by sequence-based disorder scores.

Evolutionarily young proteins were informative exceptions: several tolerated substantial expression despite having few confidently predicted structural regions (**Figure 3E**). These observations suggest two possible evolutionary trajectories from relatively benign young proteins: acquisition of well-defined structures that increases tolerance, or acquisition of interaction-prone regions that increases toxicity at high abundance. This model links the emergence of new proteins to the evolution of their expression constraints and complements evidence that newly emerged yeast sequences can acquire adaptive functions (Vakirlis et al., 2020).

The association with sulphur content adds amino acid composition to these constraints (**Figure 3G**). It is consistent with our previous finding that excess cysteine-containing proteins can disrupt protein processing and cause cytotoxicity (Moriya, 2020; Namba et al., 2022). Together, these associations help explain why overexpression toxicity cannot be reduced to a common cost of protein production. General biosynthetic costs can set the upper limit for highly tolerated proteins (Kafri et al., 2016; Eguchi et al., 2018; Fujita et al., 2025), whereas the structural, compositional, and localization properties of other proteins cause growth inhibition well below that ceiling.

Our imaging analysis identified intracellular localization as one such constraint. To our knowledge, this study provides the first systematic analysis linking measured protein abundance to both localization changes and organelle morphology across a diverse set of overexpressed proteins (**Figure 4**). Previous systematic imaging had revealed cellular phenotypes caused by overexpression (Sopko et al., 2006); introducing a measured abundance axis allowed us to resolve how these phenotypes develop as protein levels increase. High tolerance was associated with remaining cytoplasmic without entering an aggregation-dominated state (**Figure 4E**). This extends reporter-based evidence that localization processes can limit overexpression (Kintaka et al., 2016; Kintaka et al., 2020) to a diverse set of native proteins. The spatial organization of the cell is therefore an important determinant of how much excess protein it can accommodate.

The structures accommodating excess proteins also changed with increasing abundance (**Figure 4G**; **Figures S32 and S33**). Moreover, the negative association between expression and nuclear area for ER-localized proteins suggests that these effects can extend beyond the compartment containing the overexpressed protein (**Figure 4G**). Abundance–structure relationships thus offer a way to investigate both the capacity of individual compartments to accommodate proteins and how changes in one organelle affect another.

Protein aggregation has long been implicated as a major mechanism of protein toxicity (Bucciantini et al., 2002; Olzscha et al., 2011). In our analysis, aggregation-positive proteins tended to have lower IE50 values, but the expression level at which aggregation first appeared did not correlate significantly with IE50 (**Figure 4**). This distinction is consistent with evidence that sequestration can protect cells (Escusa-Toret et al., 2013) and that overloading degradation pathways, rather than aggregate formation itself, can cause toxicity (Namba & Moriya, 2024). Importantly, the image-based Aggregate class did not necessarily represent physical protein aggregates. Fun12 and Pex22 were assigned to this class, yet closer examination revealed distinct intracellular states and structures rather than simply discrete aggregates (**Figures 4 and 5**). Separating these states is therefore essential for understanding how accumulation relates to toxicity.

The striking structures generated by Fun12, Nup60, and Pex22 show that extreme overexpression can expose sequence-encoded activities that are inconspicuous at normal abundance (**Figure 5B-G**).

Membrane proliferation induced by overexpression is not new: karmellae and organized smooth ER provide established examples (Wright et al., 1988; Snapp et al., 2003). Our observations place such phenomena within a broader spectrum of abundance-dependent intracellular organization. Diverse proteins produced distinctive localization patterns (**Figure 5A**), and analysis of selected examples mapped their structure-forming activities to specific sequence regions. Protein abundance therefore determines not only the burden imposed on the cell, but also which intracellular structures and states a protein sequence can generate.

Extensive reorganization is not necessarily incompatible with growth. Pex22 combined a high IE50 with large-scale membrane remodeling despite its low fraction of residues with pLDDT > 90 and low Cytoplasm score (**Figures 3C and 5**). Its variants also lacked the correspondence between membrane morphology and IE50 observed for Nup60 variants (**Figures S36 and S37**). What matters may therefore be not simply whether an unusual structure forms, but which interactions and cellular processes it engages. Pex22 normally localizes to the peroxisomal membrane (Koller et al., 1999; Halbach et al., 2009), and peroxisomes are dispensable for growth under the conditions used here. Structures formed upon Pex22 overexpression may therefore spare essential organelle functions, explaining its low toxicity despite extensive membrane remodeling.

The framework established here opens a route to classifying overexpression toxicity by mechanism. Expanding the protein set should reveal additional predictors and distinguish their contributions more precisely. Closely related paralogs offer particularly informative comparisons because they share much of their sequence background. Differences in IE50 and abundance-dependent localization can guide sequence swaps and targeted mutations that separate the effects of folding, interaction regions, targeting signals, and amino acid composition. Combining these comparisons with perturbations of cellular processing pathways and reporters of cellular state should move the analysis from statistical associations toward the processes that determine each protein’s expression limit.

This approach also has potential applications in protein design and diseases associated with increased protein abundance. Designed proteins must not only fold and function but also be tolerated at the abundance required for their intended use. IE50 offers a way to assess this requirement and identify sequence changes that reduce cellular burden without compromising activity. In disease models, abundance-based measurements could distinguish increased accumulation from increased toxicity at a given abundance and reveal how variants alter localization, interactions, or protein processing. More broadly, these experiments provide a means to examine how coding-sequence properties and expression regulation jointly shape protein function and fitness (Venkataraman & Landry, 2026).

We analyzed about 80 proteins in this study. Finding clear biological relationships in a set of this size shows how much can be learned from this approach. Applying it to larger protein sets should reveal additional constraints and increase the resolution with which their effects on protein expression limits can be distinguished. Expression limits may also change with genetic background and growth conditions. Indeed, a genome-wide comparison across 15 *S. cerevisiae* isolates revealed strong strain-specific differences in the consequences of gene overexpression (Robinson et al., 2021). We used C-terminal GFP fusions to measure protein abundance at high throughput. This made the large-scale analysis possible, but tag position itself can affect protein localization and cellular fitness (Weill et al., 2019). Some of the localization or growth phenotypes observed here may therefore have been influenced by the tag. Future studies using broader protein sets, different cellular contexts, and less perturbative methods for measuring protein abundance should reveal how broadly the constraints identified here apply.

## Materials and methods

Strains, plasmids, assay kits, and reagents used in this study are listed in Data S1.

### Strains, plasmids, and growth media

BYW2, a BY4741-derived strain constructed in this study by integrating the WTC846 repressor cassette at the *MET15* locus, was used as the host for gTOW2.0 expression. The parental strain BY4741 has the genotype *MATa his3Δ1 leu2Δ0 met15Δ0 ura3Δ0* (Brachmann et al., 1998). The WTC846 cassette contains TetR–NLS–Tup1 expressed from the *RNR2* promoter and TetR–NLS expressed from the P7tet.1 promoter (Azizoglu et al., 2021) (FRP2374). Plasmids were constructed using the homologous recombination machinery of yeast (Oldenburg et al., 1997). The cloned region of each plasmid was confirmed by DNA sequencing. Unless otherwise indicated, cells were cultured at 30°C in synthetic complete (SC) medium lacking the nutrients required for plasmid selection.

### Construction of the gTOW2.0 expression library

***Relevant to Figures 1—5*.** Target genes were expressed using the gTOW2.0 system, which combines an aTc-inducible expression system with copy-number amplification driven by selection for the defective *LEU2* allele, *leu2-89*. Target proteins were designed as C-terminal fusions to moxGFP (Costantini et al., 2015) through a nine-amino-acid linker (GGSGGGSGG). The gTOW2.0 expression plasmid contains the P7tet.1 promoter, a *PYK1* terminator, a 2μ replication origin, and *URA3* and *leu2-89* selectable markers. The initial chromosome I target set comprised 94 genes annotated in SGD, including dubious ORFs. Subsequent experiments were conducted using genes for which correctly assembled insert-containing plasmids were successfully obtained. As the objective was to establish a sufficiently large chromosome I protein set rather than to generate a complete library covering all 94 targets, constructs that failed during the initial cloning procedure were not systematically reconstructed.

### Analysis of previous genome-wide overexpression studies

***Relevant to Figure S1.*** Previously published genome-wide gene-overexpression studies in *Saccharomyces cerevisiae* were compiled to compare the experimental systems used and the genes reported to cause growth defects upon overexpression. For each study, the copy-number system, promoter, number of genes analyzed, and genes classified as toxic upon overexpression were recorded. Toxic genes were identified according to the criteria defined in the original studies, and genes were matched across studies using systematic ORF identifiers. To compare toxicity classifications across studies, each gene was assigned a binary toxic/non-toxic classification for each publication. Genes classified as toxic in at least one publication were defined as toxic in the combined dataset, whereas genes not classified as toxic in any publication were defined as non-toxic. The degree of agreement among studies was further summarized by the number of publications in which each gene was classified as toxic and grouped into three categories: ≥2 publications, 1 publication, or 0 publications. When an explicit toxicity classification was not provided by the original study, the reported fitness data were reanalyzed to derive a binary toxicity classification. The comparison included seven publications: Gelperin et al. (2005), Sopko et al. (2006), Yoshikawa et al. (2011), Douglas et al. (2012), Tomala and Korona (2013), Makanae et al. (2013), and Arita et al. (2021). Toxic genes were defined as those included in the available positive-gene list for Gelperin et al.; those included in the toxic-gene list in Table S6 for Sopko et al.; genes assigned to growth-behavior category 0 (“Slow”) for Yoshikawa et al.; those included in the positive-gene subset of Table S10 for Douglas et al.; genes with mean normalized fitness <0.91 for Tomala and Korona; genes with mean plasmid copy number under the LU condition <100 and the corresponding reported *p*-value <0.05 for Makanae et al.; and genes classified as “Toxic = Yes” at 100 nM β-estradiol in medium 1 for Arita et al. For each study, genes meeting the specified criterion were assigned as toxic and all other genes as non-toxic. Thus, the non-toxic category included genes absent from the source data or lacking an assessable measurement and did not necessarily indicate experimentally confirmed non-toxicity. The criteria for Tomala and Korona (2013) and Makanae et al. (2013) were operational definitions used for this comparison.

### Flow-cytometric measurement of protein expression and cell viability

***Relevant to Figure 1 and Figures S2--S8***. Cells were precultured overnight in SC-Ura or SC-Leu/Ura medium in 96-well plates at 30°C under static conditions. The overnight cultures were used as the 0-h samples before induction. For induction, 50 μL of each overnight culture was transferred into 150 μL of fresh SC-Ura or SC-Leu/Ura medium containing either no aTc or aTc at a final concentration of 500 nM. Samples were collected at 6 and 24 h after induction. Cells were washed twice with 200 μL PBS and stained with Zombie Red to assess cell viability. For each sample, fluorescence signals from 10,000 cells were measured using a CytoFLEX S flow cytometer (Beckman Coulter). Throughout this study, cultivation in SC-Ura is referred to as the standard multi-copy condition, whereas cultivation in SC-Leu/Ura is referred to as the gTOW high-copy condition.

#### Flow-cytometry data processing

Flow-cytometry data were processed in Python using FlowKit (White et al., 2021). Raw fluorescence values were obtained from the FCS files. Zombie Red fluorescence was analyzed using the FL11-A channel and GFP fluorescence using the FL5-A channel. Events for which either fluorescence value was unavailable were excluded from the corresponding analysis. Cells were classified as live or dead using a fixed Zombie Red fluorescence threshold of 40,000 arbitrary units (AU). Events with Zombie Red fluorescence <40,000 AU were classified as live, whereas events with fluorescence ≥40,000 AU were classified as dead. GFP fluorescence for each sample was quantified as the arithmetic mean fluorescence of live cells. Cell viability was calculated as

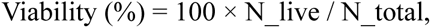

where *N_live* and *N_total* denote the numbers of live and total analyzed events, respectively. Well No. 96 contained *TDH3*pro-moxGFP, which was initially included as a maximum-expression control. Because the Vector and MOX controls were sufficient for the subsequent analyses, *TDH3*pro-moxGFP was omitted from later measurements and excluded from all analyses. For visualization of fluorescence distributions, GFP intensities were binned on a logarithmic scale between 1 and 10^7 AU. Histograms were normalized to the maximum bin frequency within each distribution. Live and dead populations were displayed separately where indicated.

#### Analysis of protein expression and cell viability across conditions

For each target protein and experimental condition, mean GFP fluorescence among live cells and the percentage of viable cells were used as the sample-level measures of protein expression and cell viability, respectively. These values were used to construct the expression and viability heatmaps, rank plots, and distributions shown in **Figures 1 and S3--S6**. **Figures S3--S6** summarize measurements obtained at 0, 6, and 24 h under standard multi-copy and gTOW high-copy conditions, with or without 500 nM aTc. Pairwise relationships among GFP-expression measurements obtained under different conditions were quantified using Pearson correlation coefficients. The same analysis was independently performed for cell viability to compare the consistency and divergence of the measured phenotypes across culture conditions and time points, as shown in **Figure S7**.

#### Analysis of expression and viability changes over time

Changes in protein expression and cell viability between 6 and 24 h after induction with 500 nM aTc were calculated separately under standard multi-copy and gTOW high-copy conditions. The change in protein expression was defined as

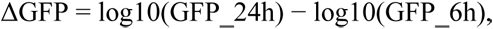

and the change in viability as

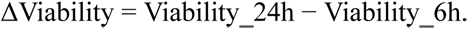

Pearson correlation coefficients and corresponding *p*-values were calculated between ΔGFP and ΔViability, as shown in **Figure S8**.

#### Analysis of the relationship between protein expression and fitness

To examine the relationship between protein expression and cellular fitness, flow-cytometric GFP measurements were integrated with maximum growth-rate measurements obtained using the microplate-reader assay described below. For each medium condition, maximum growth rates measured at 500 nM aTc were expressed relative to the corresponding empty-vector control. For each target protein, mean live-cell GFP fluorescence at 0 h, 6 h, and 24 h was compared with relative maximum growth rate. GFP fluorescence was log10-transformed for statistical analysis, and Pearson correlation coefficients were calculated between log10 GFP fluorescence and relative maximum growth rate.

#### Expression--viability trajectories

To visualize the temporal relationship between protein expression and loss of cell viability, each target protein was represented by its mean live-cell GFP fluorescence and viability at 0 h, 6 h, and 24 h.

Measurements at 0 h were obtained before induction, whereas the 6- and 24-h measurements were obtained after induction with 500 nM aTc. Arrows connected the 6- and 24-h measurements for each target protein to show changes in mean live-cell GFP fluorescence and viability.

### Measurement of growth and protein expression using a microplate reader

***Relevant to Figure 2 and Figures S9--S16.***

#### Culture conditions

For each strain, an overnight preculture was performed in SC-Ura medium. Cells were inoculated from a single colony into 200 μL medium in a 96-well plate and incubated at 30°C. Subsequently, 1 μL of the preculture was transferred into 100 μL of fresh medium in 384-well plates containing aTc at final concentrations of 0, 50, 150, or 500 nM. Cultures were grown under either standard multi-copy conditions (SC-Ura) or gTOW high-copy conditions (SC-Leu/Ura). Each condition was analyzed using four biological replicates.

#### Plate-reader measurements

Cell growth and GFP fluorescence were monitored using a Tecan Infinite F Nano+ microplate reader during continuous incubation at 30°C. Optical density was measured at 595 nm (OD595), and GFP fluorescence was recorded using excitation and emission wavelengths of 480 and 535 nm, respectively. Measurements were collected every 10 min for at least 48 h.

#### Preprocessing and quality control of plate-reader data

OD595 and GFP time-course measurements were processed in Python. To reduce short-timescale measurement noise, both OD and GFP time series were smoothed using a five-point moving average. Wells showing an abrupt decrease in raw OD595 of ≥0.2 between adjacent time points were classified as measurement errors and excluded from subsequent growth and fluorescence analyses. Only conditions for which both OD and GFP measurements were available were included in downstream analysis. Replicates were assigned within each combination of target protein, aTc concentration, medium condition, and experimental batch.

#### Calculation of maximum growth rate

Maximum growth rate (MGR) was estimated from the OD595 time series using a sliding-window approach. A window of 24 consecutive measurements, corresponding to 4 h, was moved across the smoothed OD time series in one-time-point increments. Within each window, linear regression of OD595 against time was performed, and the slope was calculated. The window showing the largest positive slope was selected, and its slope was used as the MGR for that well. For cultures whose final raw OD595 was ≤0.5, MGR was set to 0 and the culture was classified as non-growing. For these cultures, fluorescence-derived values calculated using the same procedure were retained for plotting.

#### Quantification of maximum fluorescence level

Protein expression was quantified from GFP fluorescence within the time window in which MGR was observed. For each well, raw GFP fluorescence values corresponding to the 4-h maximum-growth window were extracted. The maximum GFP fluorescence observed within this window was identified, and the raw OD595 value measured at the same time point was used to normalize fluorescence. The resulting expression measure was calculated as

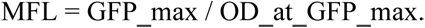

This GFP/OD value was used as the maximum fluorescence level (MFL) for subsequent analyses. Wells excluded because of abnormal OD measurements were also excluded from MFL analysis. This procedure preserves the temporal correspondence between protein-expression measurements and the phase of maximal cellular growth rather than using the maximum fluorescence observed over the entire culture period.

#### Normalization of growth measurements

Within each experimental batch and medium condition, MGR values were normalized to the mean MGR of the empty-vector controls, which was set to 100%.

#### Normalization of fluorescence measurements

MFL values were normalized separately for each experimental batch. For each batch, the minimum MFL measured for the empty-vector control was used as the lower reference. The upper reference was defined as the highest mean MFL observed among all target-protein/aTc combinations under gTOW high-copy conditions. Normalized MFL values were scaled from 1 to 100 according to

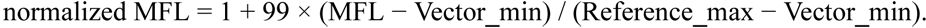

Values below 1 after scaling were set to 1. The resulting normalized GFP/OD metric is reported as MFL(/moxGFP) in the analysis tables. In all five experimental sets, the condition with the highest mean fluorescence was MOX under –LeuUra conditions with 500 nM aTc.

#### Analysis of expression and growth across induction conditions

Normalized MFL and MGR values were compared across the four aTc concentrations under standard multi-copy and gTOW high-copy conditions. For heatmaps and summary plots, replicate measurements for each target protein and induction condition were averaged. Pairwise relationships among MFL measurements obtained under different culture and induction conditions were evaluated using Pearson correlation coefficients. Equivalent pairwise comparisons were performed for MGR. Expression--growth trajectories were generated for each target protein by plotting normalized MFL against normalized MGR across increasing aTc concentrations. **Figures S9--S13** show the corresponding time-course profiles, condition-to-condition comparisons, and individual expression--growth relationships.

#### Calculation of 50% inhibitory expression levels (IE50)

The 50% inhibitory expression level (IE50) was defined as the normalized fluorescence-based expression level at which relative growth rate decreased to 50% of the vector control. IE50 was calculated independently for each biological replicate and separately under standard multi-copy and gTOW high-copy conditions. For each replicate, measurements were ordered by aTc concentration. MFL values were log10-transformed, and consecutive induction conditions were examined to identify the first interval in which MGR crossed 50%. When a crossing was detected, IE50 was estimated by linear interpolation of log10(MFL) between the two flanking measurements:

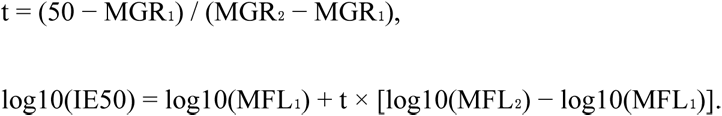

IE50 was then obtained by back-transformation to the linear scale. If MGR remained ≥50% at all measured induction levels, the highest observed MFL was used as a lower-bound estimate of the expression limit. If an IE50 value could not be obtained under gTOW high-copy conditions but a corresponding value was available under standard multi-copy conditions, the standard multi-copy value was used for the gTOW high-copy estimate. These substituted values were explicitly flagged in the processed data. Replicate-level IE50 values were subsequently averaged for each target protein to obtain the values used in gene-level analyses and figures. Gene-level IE50 estimates, including lower-bound estimates and values substituted from –Ura conditions, were included in subsequent correlation and regression analyses when the required measurements were available.

#### Comparison of expression limits between standard multi-copy and gTOW high-copy conditions

IE50 values obtained under standard multi-copy and gTOW high-copy conditions were compared across target proteins using Pearson correlation analysis. To evaluate the degree to which the measurable expression range was extended by gTOW high-copy selection, the ratio of IE50 under gTOW high-copy versus standard multi-copy conditions was calculated for each protein and compared with IE50 under gTOW high-copy conditions using Pearson correlation and linear regression.

#### Correlation structure among expression, growth, viability, and expression-limit measurements

To compare the relationships among experimental measurements, Pearson and Spearman correlation coefficients were calculated among GFP fluorescence measured by flow cytometry, MFL measured by plate reader, MGR, cell viability, and IE50 under standard multi-copy and gTOW high-copy conditions. The Pearson and Spearman correlation matrices were hierarchically clustered separately using average linkage and a distance of 1 − *r*, where *r* denotes the corresponding correlation coefficient. Pearson and Spearman coefficients obtained for all measurement pairs were also directly compared.

#### Comparison of IE50 with flow-cytometric GFP measurements

IE50 values were compared with GFP fluorescence measured by flow cytometry at 6 h after induction with 500 nM aTc under standard multi-copy and gTOW high-copy conditions. Pearson and Spearman correlation coefficients and corresponding *p*-values were calculated, and linear regression was used to determine *R*².

### Protein features used for expression-limit analysis

A proteome-wide feature dataset was compiled from sequence-, structure-, expression-, evolutionary-, and localization-related information. For the analyses presented in **Figure 3**, a selected parameter set was merged with the experimentally determined IE50 values using gene identifiers. The input parameter table contained 58 numeric protein features and 12 localization annotations used in the analysis. The analyzed features included amino-acid composition, molecular weight, codon adaptation index (CAI) (Engel et al., 2025), AlphaFold-derived structural parameters (Jumper et al., 2021) including the fraction of residues in different pLDDT ranges, predicted subcellular localization, and other sequence and physicochemical properties (Zhao et al., 2021). Subcellular localization scores were obtained using DeepLoc 2.0 (Thumuluri et al., 2022). The protein-feature names and values used for these analyses are provided in Parameters_merged.csv in the **Figure 3** data directory of the Namba_gTOW2 GitHub repository. Missing values were retained as missing and excluded pairwise from individual statistical analyses rather than being imputed.

#### Correlation analysis of protein expression limits and protein features

Associations between individual protein features and IE50 were analyzed separately under the standard multi-copy (SC-Ura) and gTOW high-copy (SC-Leu/Ura) conditions. For each protein feature, Pearson and Spearman correlation coefficients were calculated using proteins for which both the feature and IE50 measurements were available. Statistical significance was evaluated using the corresponding two-sided correlation tests. *P*-values obtained across the tested protein features were corrected for multiple testing using the Benjamini--Hochberg false-discovery-rate (FDR) procedure. An FDR-adjusted *p*-value <0.05 was considered significant. To examine whether associations between protein features and expression limits were conserved between the two expression systems, correlation coefficients obtained under standard multi-copy and gTOW high-copy conditions were compared between conditions.

#### Multiple regression analysis of major expression-limit determinants

To determine the extent to which the major independently identified protein properties collectively explained variation in expression limits, multiple linear regression models were constructed using the fraction of residues with pLDDT >90, the DeepLoc2 Cytoplasm score, and Sulphur content as explanatory variables and IE50 as the response variable. Models were fitted separately for standard multi-copy and gTOW high-copy conditions using proteins with complete measurements for all variables. Model performance was evaluated using the coefficient of determination (*R*²). *R*² was calculated using the same data used to fit the model. Models were fitted by ordinary least squares with an intercept, using explanatory variables without standardization.

#### Residual correlation analysis

Because several protein parameters covary with general properties related to protein expression and size, residual analyses were performed to determine whether correlations with IE50 persisted after accounting for these effects. For each experimental measurement, linear regression was performed using CAI and molecular weight as explanatory variables. The residual for each protein was calculated as the difference between the observed value and the value predicted by the regression model:

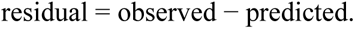

Pearson and Spearman correlations between these residual values and individual protein features were then calculated using the same pairwise-complete procedure as for the raw measurements. *P*-values were corrected across protein features using the Benjamini--Hochberg FDR procedure. Raw and residual correlation coefficients were compared to determine which associations could be explained by covariation with CAI and molecular weight and which remained detectable after these effects were removed.For the residual analysis in **Figure S18C, D**, residuals were calculated as the observed IE50 under –LeuUra conditions minus the value predicted by the three-parameter model incorporating Cytoplasm score, the fraction of residues with pLDDT > 90, and sulphur content. Pearson and Spearman correlations between these residuals and individual protein parameters were then calculated. For the color classification in **Figure S18C, D**, correlations with Benjamini–Hochberg-adjusted *p*-values < 0.10 were classified as significant.

#### Quantile regression analysis

Quantile regression was used to test whether protein properties were associated with the upper boundary of experimentally accessible expression levels. Analyses were performed separately for IE50 under standard multi-copy and gTOW high-copy conditions. For each protein feature, IE50 was treated as the response variable and the feature as the explanatory variable. A linear quantile-regression model was fitted to estimate the conditional 90th percentile of IE50 (τ = 0.9) using statsmodels.api.QuantReg (Seabold & Perktold, 2010). Models included an intercept and were fitted with a maximum of 10,000 iterations. Features with insufficient valid observations or zero variance were excluded. For comparison of regression coefficients among features, both IE50 and the explanatory protein feature were independently min--max normalized to the range 0--1 before model fitting. The regression slope and corresponding *p*-value were obtained for each feature, and *p*-values across the tested features were corrected using the Benjamini--Hochberg procedure. For visualization of individual relationships, quantile-regression lines were also fitted to the untransformed values so that the regression could be displayed in the original units.

#### LASSO regression analysis

LASSO regression (Tibshirani, 1996) was used to identify protein features associated with variation in IE50 while simultaneously considering multiple explanatory variables. Analyses were performed separately for IE50 under standard multi-copy and gTOW high-copy conditions. Experimental measurements and numeric protein parameters were matched by gene identifier, and only proteins with complete data for all variables included in the analysis were retained. Explanatory variables were standardized to zero mean and unit variance using StandardScaler. LASSO models were fitted using LassoCV in scikit-learn (Pedregosa et al., 2011) with five-fold cross-validation to select the regularization parameter independently for the standard multi-copy and gTOW high-copy datasets. Models were fitted with a maximum of 100,000 iterations. Protein features with non-zero coefficients in the fitted models were considered features retained by the LASSO analysis, and coefficients obtained under the two expression conditions were compared.

#### Correlation analysis across individual expression and fitness measurements

To determine whether associations with protein properties were specific to IE50 or were also detectable in the underlying experimental measurements, the same correlation framework was applied to individual GFP fluorescence, viability, MGR, and MFL datasets obtained under different induction and copy-number conditions. For each experimental variable, Pearson and Spearman correlations with each protein feature were calculated using pairwise-complete observations. Benjamini--Hochberg correction was applied to the resulting *p*-values. The analyses were performed both on the raw experimental measurements and on residual measurements obtained after regression against CAI and molecular weight. This analysis corresponds to the supplementary panels comparing protein-feature associations across flow-cytometric GFP, viability, plate-reader MGR, and MFL measurements.

### High-content imaging of protein localization

***Relevant to Figure 4 and Figures S26--S30***. Yeast cells expressing target proteins fused to moxGFP were precultured overnight under eight conditions comprising standard multi-copy (SC-Ura) or gTOW high-copy (SC-Leu/Ura) medium and four aTc concentrations (0, 50, 150, and 500 nM). Each preculture was transferred to fresh medium with the same medium composition and aTc concentration and cultured for 6 h before imaging. Images were acquired using an Opera Phenix high-content screening system equipped with a 63× water-immersion objective. Bright-field images were acquired for cell identification, and GFP fluorescence images were collected at three exposure settings to cover the broad dynamic range of target-protein expression. The relative intensity scaling among the three GFP channels was approximately 1:21:147 from shortest to longest exposure. The complete **Figure 4** image-analysis workflow was organized as a multistep pipeline comprising image indexing, classifier training, single-cell inference, localization-transition analysis, and downstream visualization.

#### Single-cell image preprocessing

Individual cells were segmented from bright-field images, and the resulting masks were used to extract corresponding GFP images centered on each cell. GFP regions of interest were resized or cropped to 224 × 224 pixels for localization classification. Bright-field images were used only for cell segmentation and determination of ROI coordinates and were not provided to the localization-classification model. Cell segmentation was performed using Cellpose v3.1.1 (Pachitariu & Stringer, 2022; Stringer et al., 2021) with a custom model trained on more than 100 manually annotated cells using a human-in-the-loop training procedure. The model was trained on bright-field images and used to generate cell masks from bright-field images. For GFP fluorescence measurements, three images were acquired at different exposure times for each field. The Cellpose-derived masks were applied to the corresponding GFP images, and fluorescence was evaluated independently for each segmented cell. Images containing any saturated pixel within the corresponding cell mask were excluded. Among the remaining non-saturated images, the image providing the highest fluorescence intensity was selected for quantitative analysis.

#### Quantification of single-cell GFP fluorescence

Single-cell GFP fluorescence intensities were converted to a common scale by correcting for relative exposure/channel sensitivity and were log10-transformed for downstream localization-transition analyses. The fluorescence values obtained from the three acquisition channels were converted to a common scale using empirically determined correction factors of ×21 for ch1, ×1 for ch3, and ×1/7 for ch4. Using the MOX control, these factors were derived from the slopes of fluorescence-intensity relationships among the channels and were used for calculation of expression levels in the manuscript dataset.

#### Machine-learning classification of subcellular localization

Single-cell GFP images were classified into eight biologically interpretable localization classes: Cytoplasm, Aggregate, Nucleus, ER, Mitochondria, Nuclear membrane, Puncta/Vesicle, and Plasma membrane. Additional quality-control categories included broken or incorrectly segmented cells, low-intensity cells, out-of-focus cells, and cells with no detectable fluorescence; these classes were excluded from subsequent biological analyses. Representative images of the localization and quality-control classes are shown in **Figure S27**. At least 100 single-cell images per class were initially annotated manually. A classification model based on the DINOv2 ViT-S/14 backbone (Oquab et al., 2023) was first trained using this manually annotated dataset. The model was then applied to unlabeled images, and high-confidence predictions (>0.95) were iteratively incorporated into the training dataset. This self-training procedure was repeated until the labeled dataset contained approximately 100,000 images. The final model was then applied to the complete single-cell image dataset. Model confidence was calibrated by temperature scaling using validation-set logits before inference. Model training used AdamW (weight decay, 1 × 10−4) with a batch size of 32 for both training and validation. A new optimizer was initialized for each of the two training phases. In phase 1, the backbone was frozen and the classification head was trained for one epoch at a learning rate of 3 × 10−4. In phase 2, the final three Transformer blocks, the final backbone normalization layer, and the classification head were trained for three epochs at a learning rate of 3 × 10−5. No learning-rate scheduler was used. Training used class-weighted cross-entropy loss, with class weights based on 1/√nc, where nc is the number of training cells in class c, normalized to a mean of 1, clipped to 0.75–3.0, and renormalized to a mean of 1; validation used unweighted cross-entropy loss. Cells were randomly split into training and validation sets at an 85:15 ratio with stratification by class rather than by gene or experiment, using a maximum pool of 200,000 cells. Training images were resized to 256 pixels, randomly cropped to 224 × 224 pixels, randomly flipped horizontally, and randomly rotated by up to ±5°. Validation images were center-cropped. Image intensities were normalized using the 1st and 99th percentiles and clipped, followed by ImageNet mean and standard-deviation normalization. Random seed 42 was used for Python, NumPy, PyTorch, and CUDA, with deterministic cuDNN behavior enabled and cuDNN benchmarking disabled. The model with the highest validation macro-F1 score was retained. The training environment used Python 3.10, PyTorch 2.7.*, torchvision 0.22.*, timm 1.0.26, huggingface_hub 1.9.0, and safetensors 0.7.0. Inference was performed in evaluation mode using a single center-cropped image per cell, without test-time augmentation. Images were normalized using the 1st and 99th intensity percentiles, clipped, converted from grayscale to RGB, center-cropped to a square, resized to 256 × 256 pixels, and center-cropped to 224 × 224 pixels, followed by ImageNet mean and standard-deviation normalization. Class probabilities were obtained by applying softmax to logits scaled by a temperature of 1.5089. Each cell was assigned to the class with the highest probability, and that probability was recorded as the confidence score.

#### Validation of localization classification

Classifier performance was evaluated using manually annotated images and summarized as a confusion matrix (**Figure S28A**). Rows represent manually assigned classes and columns represent predicted classes. Predicted localizations under low-expression conditions were additionally compared with sequence-based localization probabilities obtained using DeepLoc 2.0 (**Figure S28B**). Aggregate and Puncta/Vesicle classes were excluded from direct DeepLoc category matching because equivalent categories were not available in the DeepLoc output used for this comparison. The maximum GFP-expression value obtained by high-content imaging was compared with independent protein-expression measurements obtained by flow cytometry and plate-reader analysis and with IE50 under gTOW high-copy conditions using Spearman rank correlation (**Figure S28C**).

#### Assignment of low-expression localization

The dominant low-expression class was defined as the most frequently predicted localization among up to 200 cells with the lowest GFP fluorescence for each target protein. Ties were resolved using a predefined localization-class priority. Because low fluorescence occasionally caused cells to be assigned to Cytoplasm or Puncta, the display-order localization was manually specified as ER for *YAR023C*, *DRS2*, and *SYN8* and as Nuclear for *POP5*, *FUN19*, and *ECM1*. These manual assignments were used only to determine gene order in the visualization and did not alter individual-cell predictions, localization fractions, expression measurements, or statistical group assignments.

#### Analysis of expression-dependent localization changes

*Relevant to* ***Figure 4C,D*** *and **Figures S29 and S30**.* Quality-control classes were excluded before analysis. Predictions with confidence below 0.95 were excluded from the transition-profile analysis. Single-cell localization patterns were summarized as a function of GFP-expression level. Expression bins were defined using common fixed-width boundaries spanning the minimum to maximum log10 expression value within the analyzed panel. Expression-dependent localization analyses shown in **Figure 4C** and **Figures S29 and S30** used 20 expression bins. For each target protein, at least 200 cells were required for inclusion in the expression-dependent localization analysis. Within each expression bin, at least 51 cells were required for the bin to be considered valid. Proteins with fewer than three valid expression bins were excluded from the transition-profile analysis. Within each valid expression bin, the fraction of cells assigned to each of the eight localization classes was calculated as

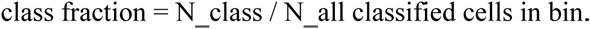

The class with the largest fraction was defined as the dominant localization class for that expression bin. For each valid bin, the expression range (expr_min and expr_max), median single-cell expression value, number of cells, dominant localization class, and fraction of each localization class were recorded. Expression-dependent localization distributions for individual target proteins are shown in **Figure S29**, whereas **Figure S30** summarizes the dominant localization classes across expression bins for all target proteins together with montages of representative cell images corresponding to these localization states.

#### Ordering and grouping of localization-transition profiles

Target proteins were grouped according to their localization at low expression and subsequent localization behavior at higher expression. Within major localization groups, proteins were ordered using IE50 under gTOW high-copy conditions for visualization. For cytoplasmic proteins, proteins that remained predominantly cytoplasmic across expression bins were distinguished from proteins that transitioned toward Aggregate localization. Other proteins were grouped according to their major low-expression localization, including Nucleus, Mitochondria, ER, Nuclear membrane, Plasma membrane, and Puncta/Vesicle. Among proteins whose dominant localization class at the lowest valid expression bin was Cytoplasm, proteins that remained cytoplasmic across all valid expression bins were classified as Cyto→Cyto, whereas those for which Aggregate became the dominant class in at least one expression bin were classified as Cyto→Agg. Proteins showing other localization transitions were excluded from this comparison.

#### Analysis of aggregation during overexpression

A target protein was classified as aggregation-positive if Aggregate became the dominant localization class in at least one valid expression bin during the expression series. The association between initial localization, defined as the dominant class in the lowest valid expression bin, and aggregation status was evaluated using a chi-square test of independence on the contingency table of initial localization class versus aggregation status (scipy.stats.chi2_contingency). IE50 values were compared between aggregation-positive and aggregation-negative proteins using a two-sided Mann--Whitney *U* test. The association between aggregation status and IE50 was additionally examined using logistic regression, with aggregation status as the binary response variable and IE50 as the explanatory variable. For aggregation-positive proteins, aggregation onset was defined using the maximum expression value of the first expression bin in which Aggregate became the dominant class. The relationship between aggregation-onset expression and IE50 was evaluated using Spearman rank correlation.

#### Statistical comparison of localization groups

IE50 under gTOW high-copy conditions and protein properties identified in **Figure 3** were compared among localization groups. These included the fraction of residues with pLDDT >90, the DeepLoc 2.0 Cytoplasm score, and sulphur content. Overall differences among localization groups were assessed using the Kruskal--Wallis test, and pairwise comparisons were performed using two-sided Mann--Whitney *U* tests where indicated.

### Quantification of mitochondrial and nuclear morphology

*Relevant to* ***Figure 4F,G*** *and **Figures S31--S33**.* Mitochondria and nuclei were labeled using Aco2–mScarlet-I3 (Aco2-RFP) and Hta2–mCherry (Hta2-RFP), respectively, in cells expressing GFP-tagged target proteins. Cells were cultured under standard multi-copy or gTOW high-copy conditions with 500 nM aTc, and organelle morphology was monitored by high-content microscopy. The single-cell quantification in **Figures S32 and S33** combined data from cells cultured for 6 h under standard multi-copy (–Ura) and gTOW high-copy (–LeuUra) conditions. Single-cell morphological features were quantified from the Aco2-RFP and Hta2-RFP images using CellProfiler. Pairwise correlations among morphological features were calculated to identify redundant measurements. Because many area- and shape-related parameters were strongly correlated, Mean_Mito_AreaShape_Area and Mean_Nuc_AreaShape_Area were selected as representative measures of mitochondrial and nuclear size, respectively. For nuclei, nuclear area and nuclear compactness were quantified at the single-cell level. These dimensionless coefficients were plotted in **Figure 4G**, with one point per target protein, and compared among proteins grouped by their low-expression localization. Organelle morphology was analyzed using CellProfiler (Stirling et al., 2021) pipelines for mitochondria (version 4.2.6) and nuclei (version 4.2.8). Previously generated cell masks were imported as cell objects. Aco2-RFP images were intensity-rescaled, masked to cell regions, and enhanced using the tubeness filter. Mitochondrial objects were identified using adaptive two-class Otsu thresholding with a 50-pixel window and an object diameter range of 3–200 pixels. Nuclear objects were identified from intensity-rescaled Hta2-RFP images using global two-class Otsu thresholding with an object diameter range of 5–80 pixels. Area and shape features were measured using MeasureObjectSizeShape. Mean_Mito_AreaShape_Area and Mean_Nuc_AreaShape_Area represent the mean area, in pixel units, of the corresponding segmented objects assigned to each cell. Mitochondrial count represents the number of segmented mitochondrial objects assigned to each cell. Nuclear compactness was calculated as

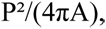

where P and A are the perimeter and area of each nuclear object, and averaged within each cell.

### Fluorescence microscopy of strongly overexpressed proteins

***Relevant to Figure 5 and Figures S34--S37***. Cells expressing GFP-fused target proteins were cultured under gTOW high-copy conditions in the presence of 500 nM aTc and observed by fluorescence microscopy. Representative images were used to visualize intracellular localization patterns under strong overexpression. Cells were precultured in SC-Leu/Ura medium without induction. For induction, 50 μL of the preculture was transferred into 150 μL of fresh SC-Leu/Ura medium containing aTc at a final concentration of 500 nM. Cells were imaged 6--12 h after induction using an Olympus FV3000 confocal microscope equipped with a 100× oil-immersion objective. GFP fluorescence was excited at 488 nm and detected at 540 nm, and bright-field and GFP images were acquired for each field. Cells were segmented from bright-field images using Cellpose v3.1.1 with the custom human-in-the-loop model described above, and the resulting binary masks were used to generate montage images for visualization. No quantitative image analysis was performed.

### Construction and analysis of Fun12, Nup60, and Pex22 variants

***Relevant to Figure 5 and Figures S35--S37***. Fragments and sequence variants of Fun12, Nup60, and Pex22 were constructed to identify sequence regions responsible for the intracellular structures observed during strong overexpression. Amino-acid positions are numbered relative to the corresponding full-length proteins and are inclusive. For Fun12 (aa 1--1002), the full-length protein and two truncation constructs were used: an N-terminal intrinsically disordered region (IDR; aa 1--399) and the remaining folded region (Fold; aa 400--1002). For Nup60 (aa 1--539), the constructs comprised the full-length protein, the N-terminal region (aa 1--162), the C-terminal region (aa 163--539), the N-terminal KR-rich region (aa 1--20), a fragment containing the amphipathic helix (AH; construct spanning aa 21--47), a fragment containing the KR-rich region and AH (aa 1--47), and a fragment containing the AH and the remaining N-terminal region (aa 21--162). KR-to-A mutants were generated in the aa 1--47 and aa 1--162 constructs by replacing residues R3, K4, R7, R8, R19, and K20 with alanine. For Pex22 (aa 1--180), the constructs comprised the full-length protein, the N-terminal KR-rich region (aa 1--14), the putative ALPS region (aa 15--32), the combined KR-rich and ALPS region (aa 1--32), aa 33--58, aa 18--58, aa 1--58, and the C-terminal folded region (aa 59--180). KR-to-A mutants were generated in both the aa 1--32 and full-length constructs by replacing the K/R residues at positions 6, 8, 11, and 13 with alanine. Detailed amino-acid boundaries and substitutions for all constructs are provided in Data S1. The boundary between the N-terminal IDR and C-terminal folded region of Fun12 was selected based on the AlphaFold2-predicted structure. The Nup60 AH was annotated based on previous studies (Cibulka et al., 2022; Mészáros et al., 2015). Pex22 residues 15–32 were selected as a putative amphipathic lipid packing sensor (ALPS) motif based on sequence characteristics described for ALPS motifs, including an amphipathic arrangement of hydrophobic and polar residues (Bigay et al., 2005; Drin et al., 2007). This designation represents a sequence-based prediction rather than a previously established lipid-packing-sensing function of Pex22. Expression limits of all variants were determined using the same microplate-reader and IE50 procedures described above.

### FRAP analysis

***Relevant to Figure 5 and Figure S35.*** Fluorescence recovery after photobleaching (FRAP) was performed using an Olympus FV-3000 confocal microscope equipped with a 100× oil-immersion objective. GFP was excited at 488 nm. After acquisition of three pre-bleach frames, photobleaching was performed at a single focal point using a high-intensity laser, resulting in a circular bleached area. Two FRAP acquisition modes were used. For visualization of fluorescence recovery over the entire field of view, conventional time-lapse imaging was performed over approximately 5 s, with approximately 100 frames acquired per measurement. These image sequences were used for qualitative visualization and were not used for quantitative FRAP analysis. Quantitative FRAP measurements were instead acquired using line-scan imaging to achieve substantially higher temporal resolution. Each line-scan time series comprised 1,000 frames, including three pre-bleach frames and 997 post-bleach frames. Post-bleach images were acquired at intervals of approximately 11–14 ms, depending on the measurement, corresponding to a total acquisition period of approximately 11–13 s. Ten cells were analyzed for each condition. Fluorescence recovery was monitored after bleaching, and images were analyzed using Fiji (Schindelin et al., 2012). FRAP recovery curves were fitted with a single-exponential recovery function,

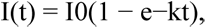

where I(t) is the normalized fluorescence intensity at time t, I0 is the asymptotic recovery amplitude, and k is the recovery rate constant. Fluorescence intensities were background-corrected by subtracting the black-background signal and normalized to the maximum corrected intensity of each measurement. The time axis was shifted such that the midpoint between the third and fourth frames was defined as t = 0, and fitting was performed from the fourth frame onward. Parameters were estimated independently for each replicate by nonlinear least-squares fitting using the Levenberg--Marquardt algorithm (scipy.optimize.curve_fit), without parameter bounds or manually specified initial values. For the plotted recovery curves, the mean fitted I0 and k values across replicates were used.

### Transmission electron microscopy

***Relevant to Figure 5 and Figures S35--S37.*** Cells were precultured in SC-Leu/Ura medium at 30°C. Subsequently, 200 μL of the preculture was transferred to SC-Leu/Ura medium containing 500 nM aTc and cultured at 30°C for 12 h. Cells were collected and transported to Tokai Electron Microscope Co., where all subsequent sample preparation and transmission electron microscopy procedures were performed. Samples were sandwiched between copper plates, rapidly frozen in liquid propane, and freeze-substituted in anhydrous ethanol. Cells were infiltrated with a 5:5 mixture of propylene oxide and Quetol-651 resin (Nissin EM Co.). Ultrathin sections of 80 nm thickness were prepared using an Ultracut UCT ultramicrotome (Leica), stained with 2% uranyl acetate and lead staining solution (Sigma-Aldrich), and observed using a JEM-1400 Plus transmission electron microscope (JEOL Ltd.).

#### Serial-section analysis

For serial-section analysis of Nup60-overexpressing cells, consecutive 80-nm sections through the same intracellular membrane structures were imaged by TEM. Consecutive sections were visually inspected to determine whether the same structure was present across adjacent sections. Structures were identified based on their spatial continuity and morphology, and the images were interpreted directly without computational alignment.

### Statistical analysis and reproducibility

Data analyses were performed primarily using Python 3.10. In the boxplots in **Figure 2B–E**, boxes span the 25th–75th percentiles with the median indicated inside each box, whiskers extend to the most extreme values within 1.5 times the interquartile range of the box edges, and points beyond the whiskers indicate outliers. Pearson correlation was used to assess linear associations, and Spearman rank correlation was used for rank-based associations. Two-group comparisons were performed using two-sided Mann--Whitney *U* tests where indicated, and comparisons among three or more groups were performed using Kruskal--Wallis tests.

## Code availability

Analysis scripts, notebooks, CellProfiler pipelines, and software environment files are available in the Namba_gTOW2 GitHub repository.

## Data availability

High-content microscopy images and associated cell-segmentation images are available in SSBD:repository (Kyoda et al., 2025) under accession ssbd-repos-000505 (HMlab_HCS_WTC-gTOW; Namba & Moriya, 2026; doi:10.24631/ssbd.repos.2026.04.505). Raw flow cytometry files and processed data tables are available in the Namba_gTOW2 GitHub repository, organized by figure.

## Acknowledgments

We thank the members of the Moriya laboratory at Okayama University for valuable discussions related to this study. We also thank the Division of Instrumental Analysis at Okayama University for providing access to the confocal microscope. We thank the members of the Boone and Andrews laboratories at the University of Toronto for helpful discussions, and Helena Friesen for her generous assistance and guidance in the use of experimental equipment. S.N. acknowledges support from the Japan Society for the Promotion of Science (JSPS) Overseas Research Fellowships.

## Author contributions

S. N. and H. M. conceived the study, analyzed the data, and wrote the manuscript. S. N. and Y. N. performed the experiments. C. B. and B. A. provided materials and equipment.

