## Supplementary Figures for "Limit-pushing overexpression reveals constraints on protein abundance"

Namba et al.

Figure S1

A

| Publication | Copy number | Promoter | Analyzed gene number | Toxic gene |
|---|---|---|---|---|
| Arita (2021) | Genomic | Z3EV | 5690 | 987 |
| Makanae (2013) | gTOW | Native | 5806 | 759 |
| Tomala (2013) | 2 µ | *GAL1/10* | 5182 | 306 |
| Douglas (2012) | Cen | *GAL1/10* | 5192 | 360 |
| Yoshikawa (2011) | 2 µ | *GAL1/10* | 4300 | 1302 |
| Sopko (2006) | 2 µ | *GAL1/10* | 5280 | 768 |
| Gelperin (2005) | 2 µ | *GAL1/10* | 5854 | 322 |

B

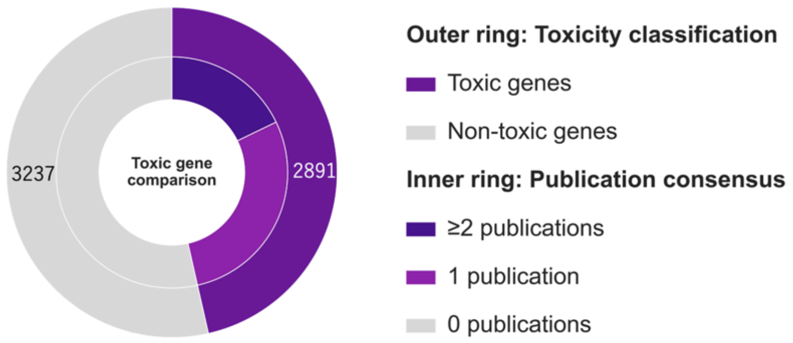


C

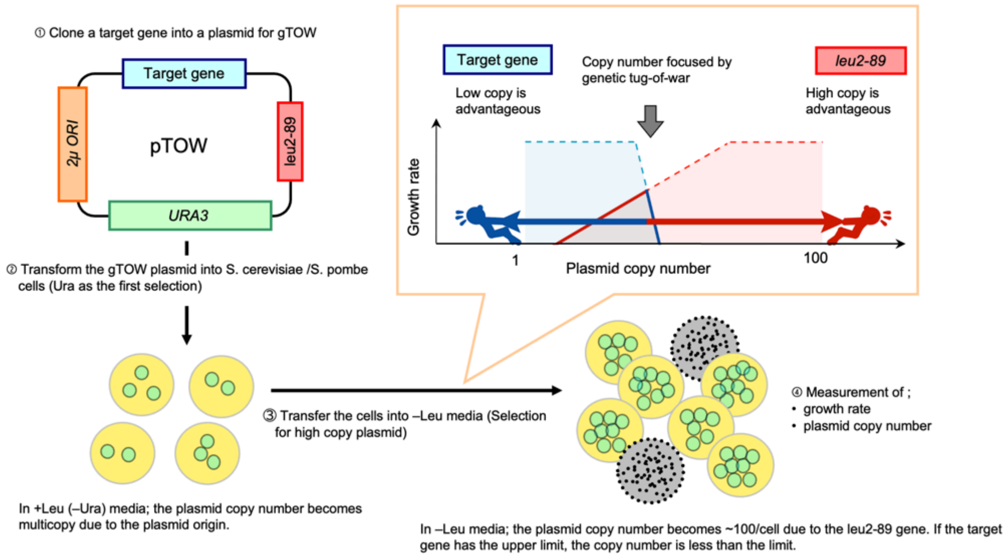


#### Figure S1. Previous genome-wide overexpression studies and the principle of the genetic tug-of-war (gTOW) method.

**(A)** Summary of previous genome-wide gene overexpression studies in *Saccharomyces cerevisiae*. The copy-number system, promoter, number of genes analyzed, and number of genes reported to cause growth defects upon overexpression are shown for each study.

**(B)** Comparison of genes classified as toxic by the studies listed in **(A)**. The outer ring indicates genes reported as toxic in at least one study (2,891 genes) and genes not reported as toxic in any study (3,237 genes). The inner ring indicates the consistency of toxicity classification across studies: genes classified as toxic in two or more studies, in only one study, or in none of the studies.

**(C)** Schematic representation of the genetic tug-of-war (gTOW) method. A target gene is cloned into a 2μ-based multicopy plasmid carrying *URA3* and the defective *LEU2* allele, *leu2-89*. Under –Ura selection, the plasmid is maintained at the copy number determined primarily by the 2μ origin. Upon transfer to –Leu medium, selection for *leu2-89* favors an increase in plasmid copy number, whereas the fitness cost imposed by increased dosage of the target gene favors a decrease in copy number. The balance between these opposing selective pressures determines the maximum tolerable copy number of the target gene, which can be quantified together with cellular growth (Moriya et al., 2006, 2012).

Figure S2

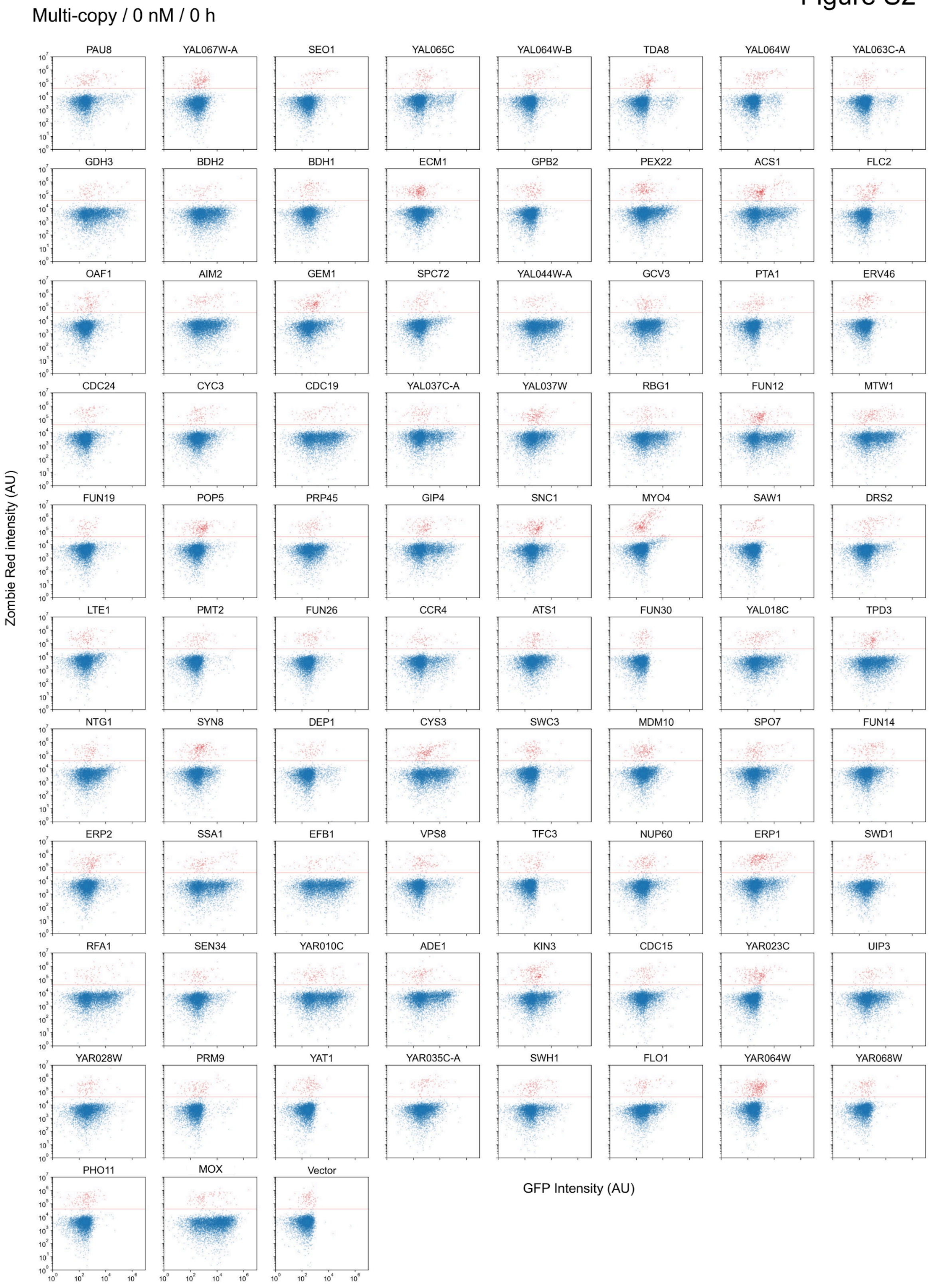
Multi-copy / 0 nM / 0 h
PAU8
YAL067W-A
SEO1
YAL065C
YAL064W-B
TDA8
YAL064W
YAL063C-A
GDH3
BDH2
BDH1
ECM1
GPB2
PEX22
ACS1
FLC2
OAF1
AIM2
GEM1
SPC72
YAL044W-A
GCV3
PTA1
ERV46
CDC24
CYC3
CDC19
YAL037C-A
YAL037W
RBG1
FUN12
MTW1
FUN19
POP5
PRP45
GIP4
SNC1
MYO4
SAW1
DRS2
LTE1
PMT2
FUN26
CCR4
ATS1
FUN30
YAL018C
TPD3
NTG1
SYN8
DEP1
CYS3
SWC3
MDM10
SPO7
FUN14
ERP2
SSA1
EFB1
VPS8
TFC3
NUP60
ERP1
SWD1
RFA1
SEN34
YAR010C
ADE1
KIN3
CDC15
YAR023C
UIP3
YAR028W
PRM9
YAT1
YAR035C-A
SWH1
FLO1
YAR064W
YAR068W
PHO11
MOX
Vector
Zombie Red intensity (AU)
GFP Intensity (AU)

### Figure S2 (continued)

Multi-copy / 0 nM / 6 h

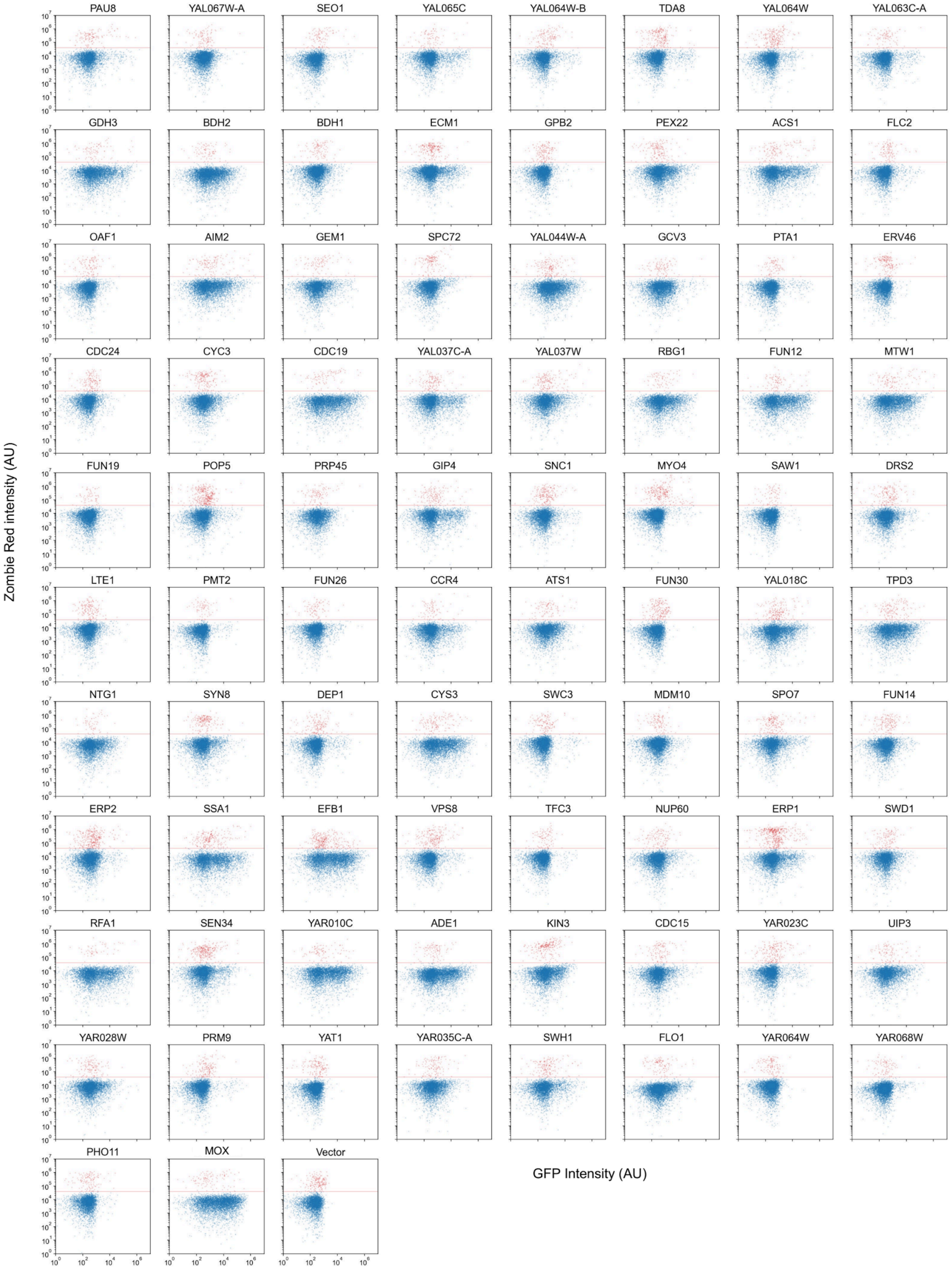

Figure S2 (continued)

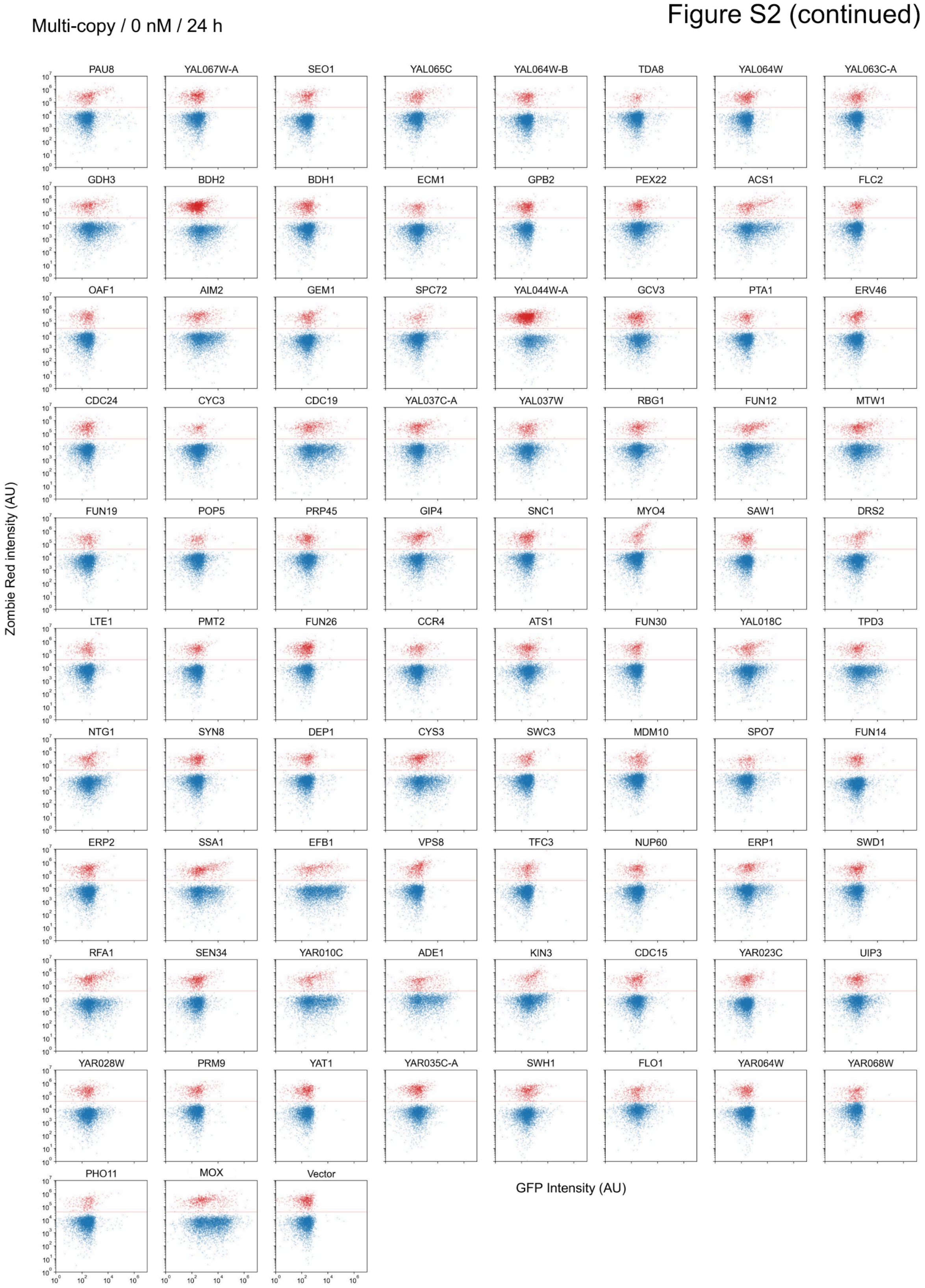

Figure S2 (continued)

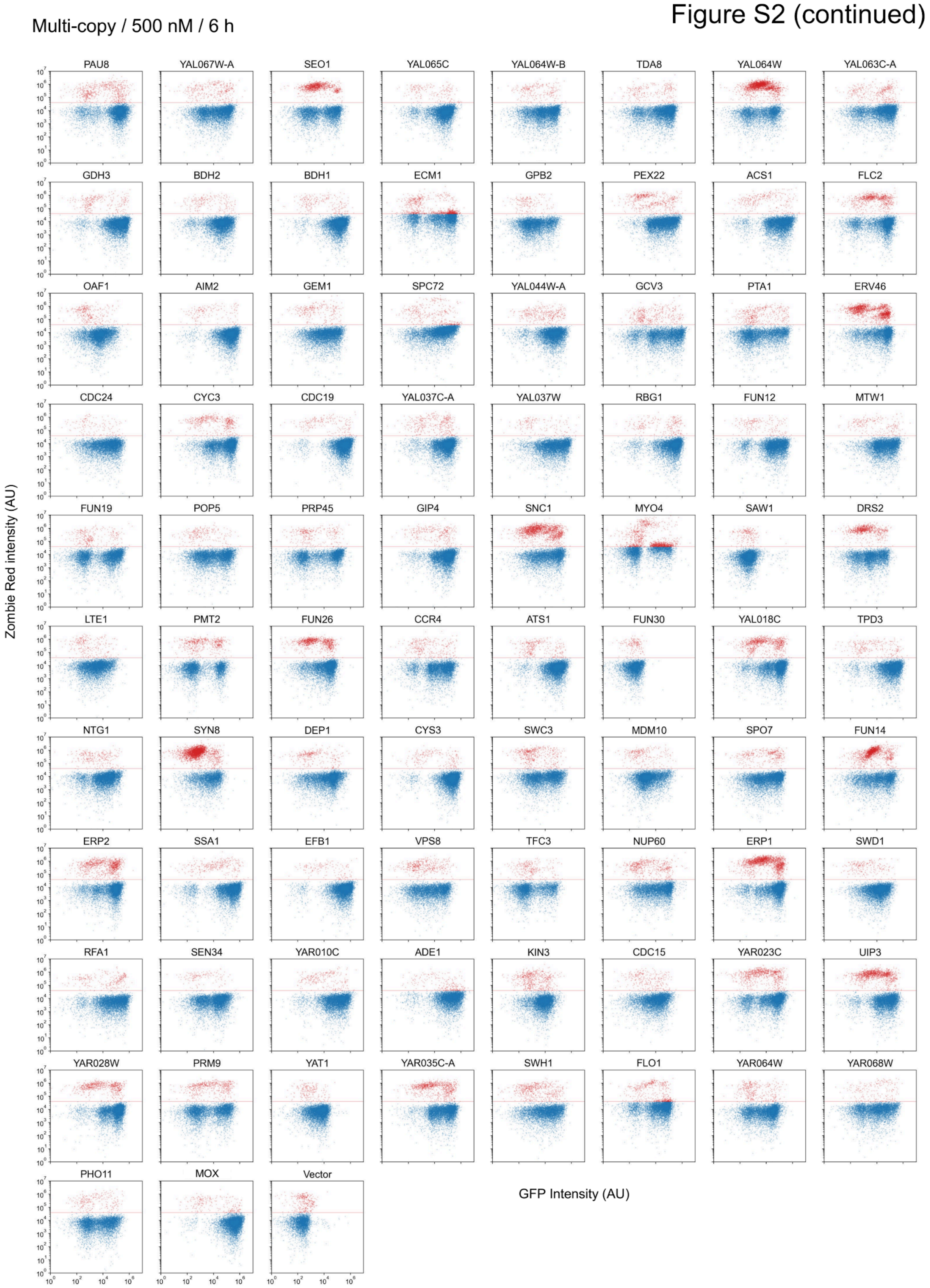

### Figure S2 (continued)

Multi-copy / 500 nM / 24 h

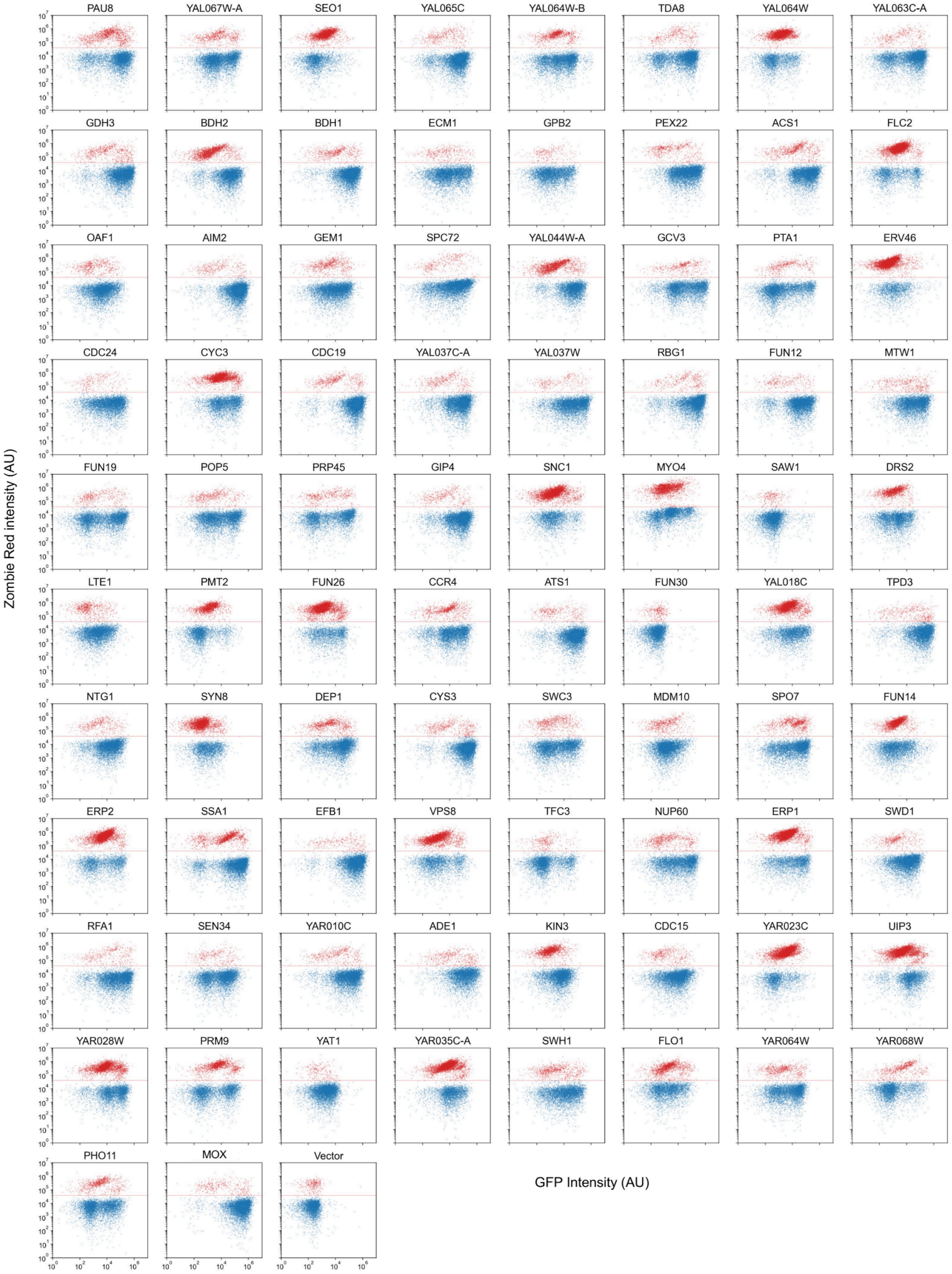

### Figure S2 (continued)

gTOW high-copy / 0 nM / 0 h

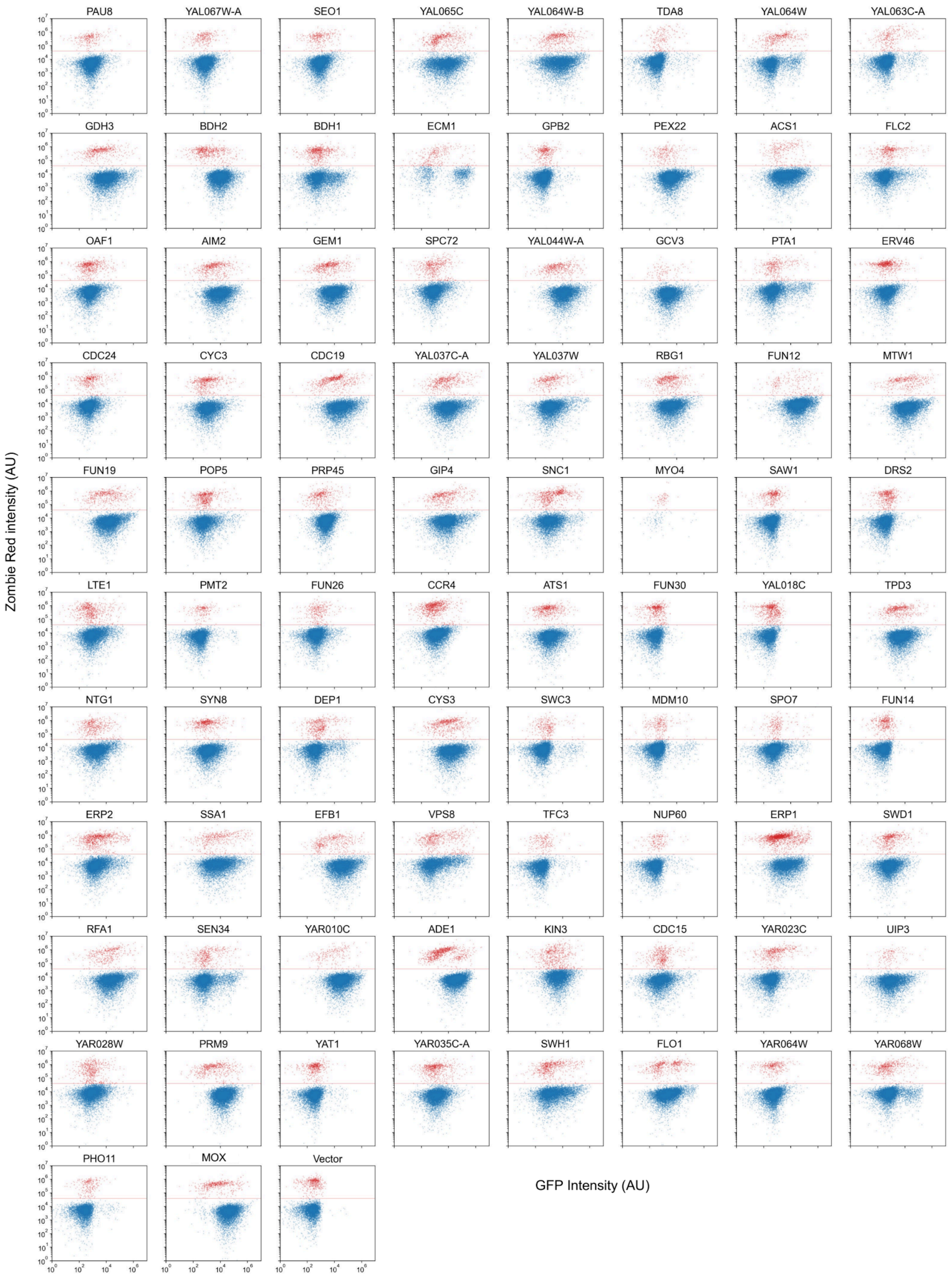

Figure S2 (continued)

gTOW high-copy / 0 nM / 6 h

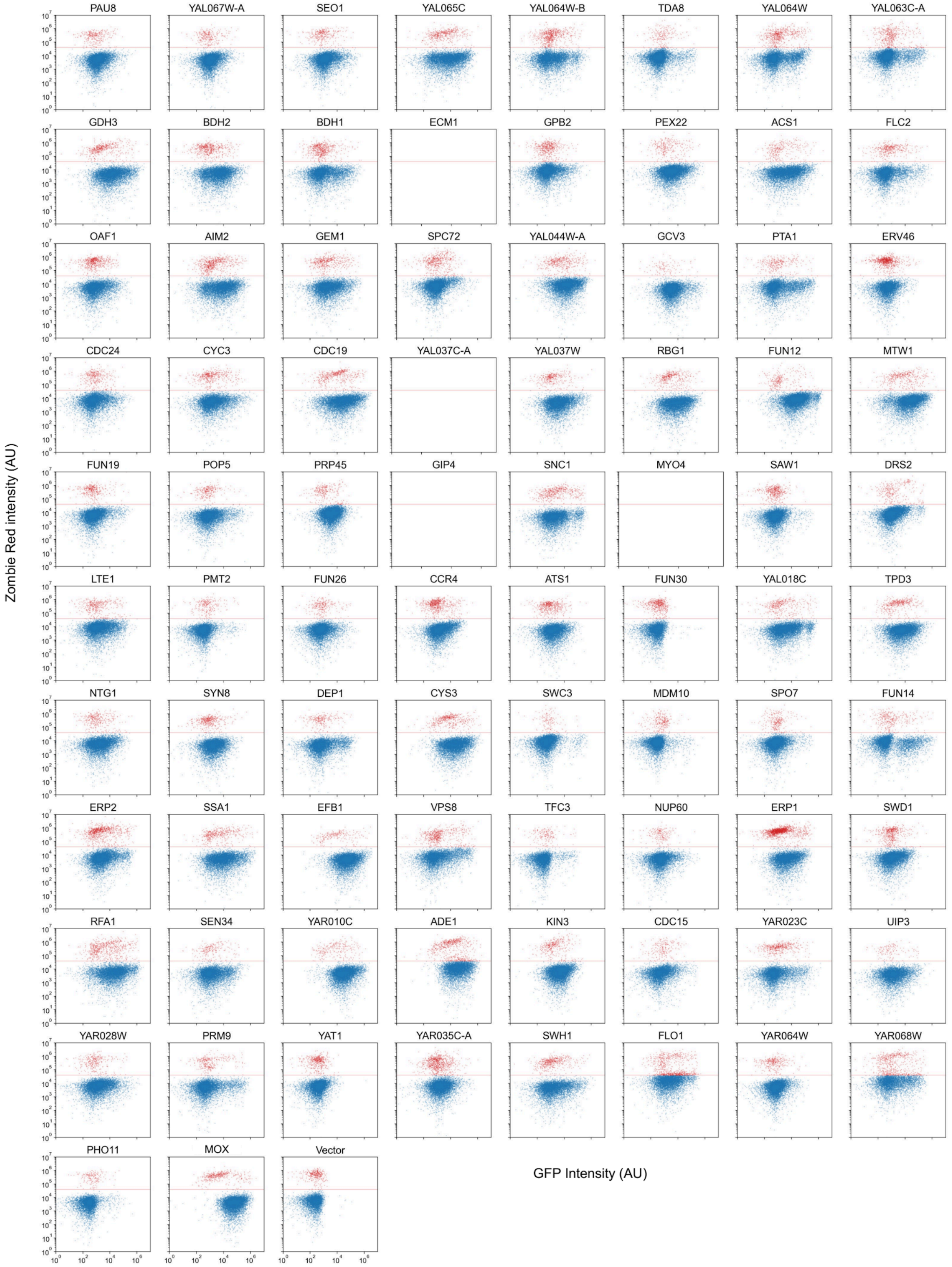

### Figure S2 (continued)

gTOW high-copy / 0 nM / 24 h

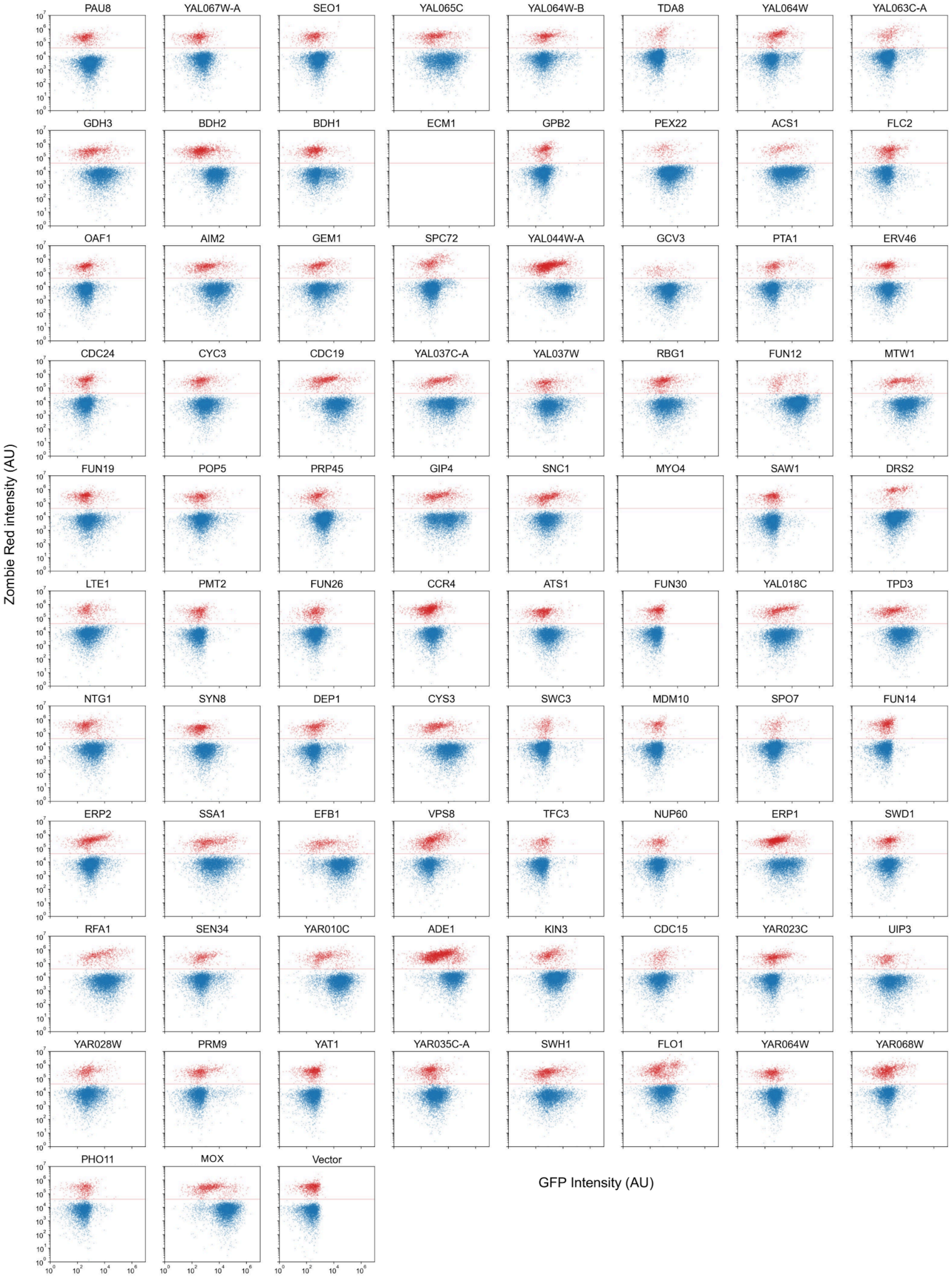

### Figure S2 (continued)

gTOW high-copy / 500 nM / 6 h

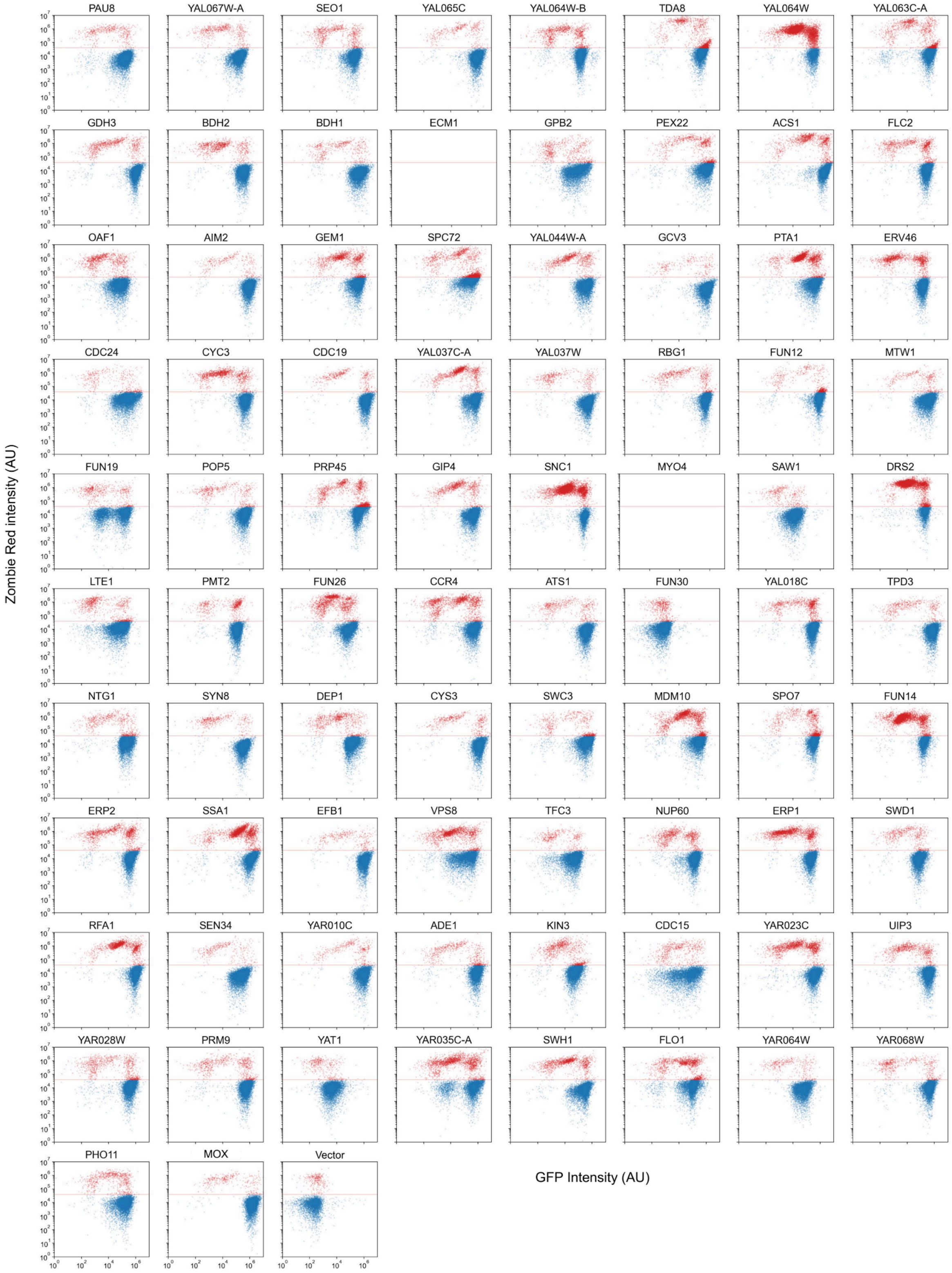

gTOW high-copy / 500 nM / 24 h

### Figure S2 (continued)

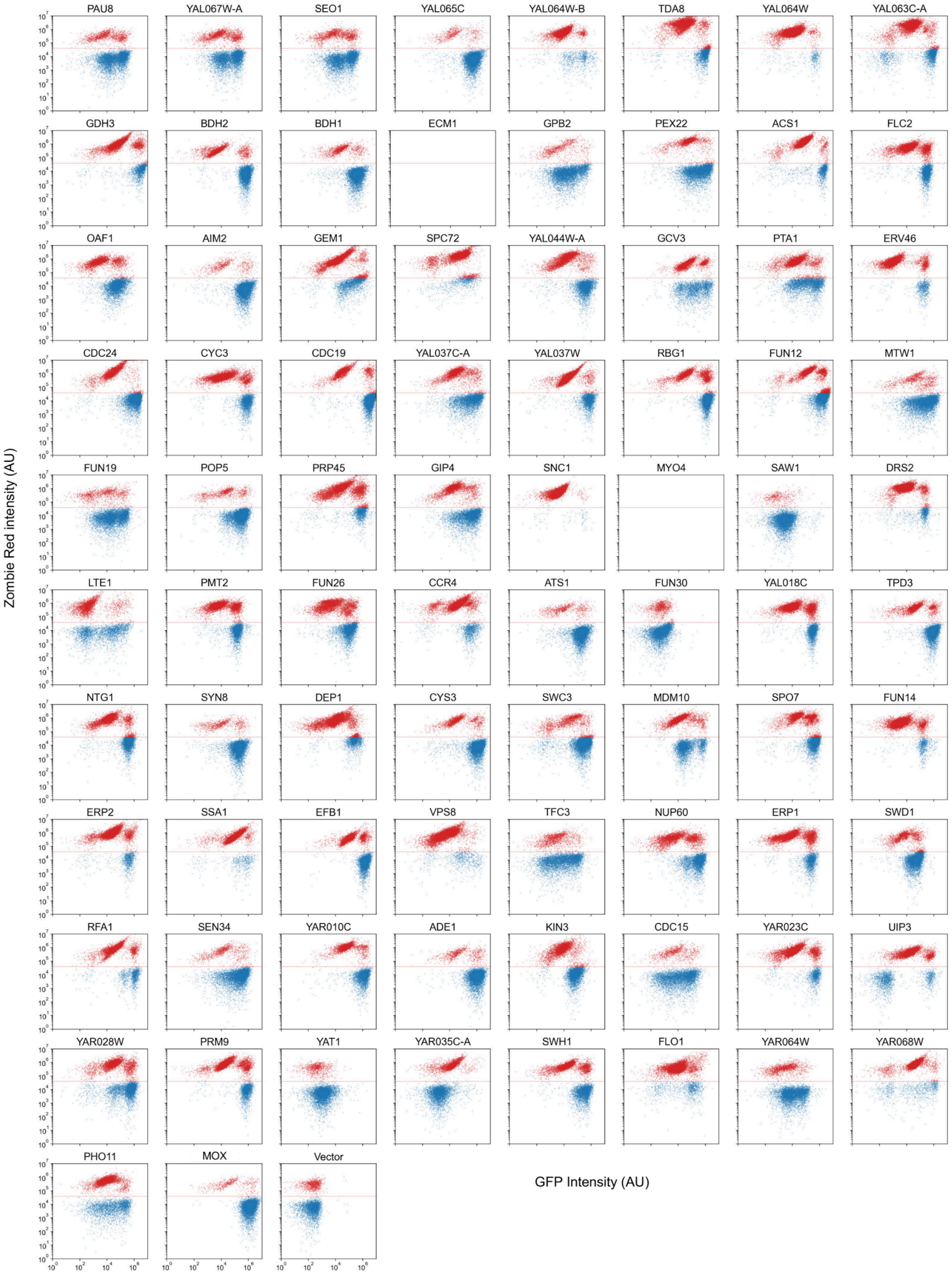

#### Figure S2. Flow cytometric analysis of protein expression and cell viability under standard multi-copy and gTOW high-copy conditions.

Strains expressing each target protein fused to moxGFP were cultured under standard multi-copy (–Ura) or gTOW high-copy (–LeuUra) conditions. Protein expression was induced with 500 nM aTc, and GFP and Zombie Red fluorescence were measured by flow cytometry at 0, 6, and 24 h. Uninduced cultures (0 nM aTc) were analyzed in parallel. Each panel represents an individual target protein. The *x*-axis indicates GFP fluorescence intensity (AU), and the *y*-axis indicates Zombie Red fluorescence intensity (AU). Live and dead cells were identified as Zombie Red-negative and Zombie Red-positive, respectively. MOX (moxGFP alone) and Vector (empty vector) are shown as controls.

Figure S3

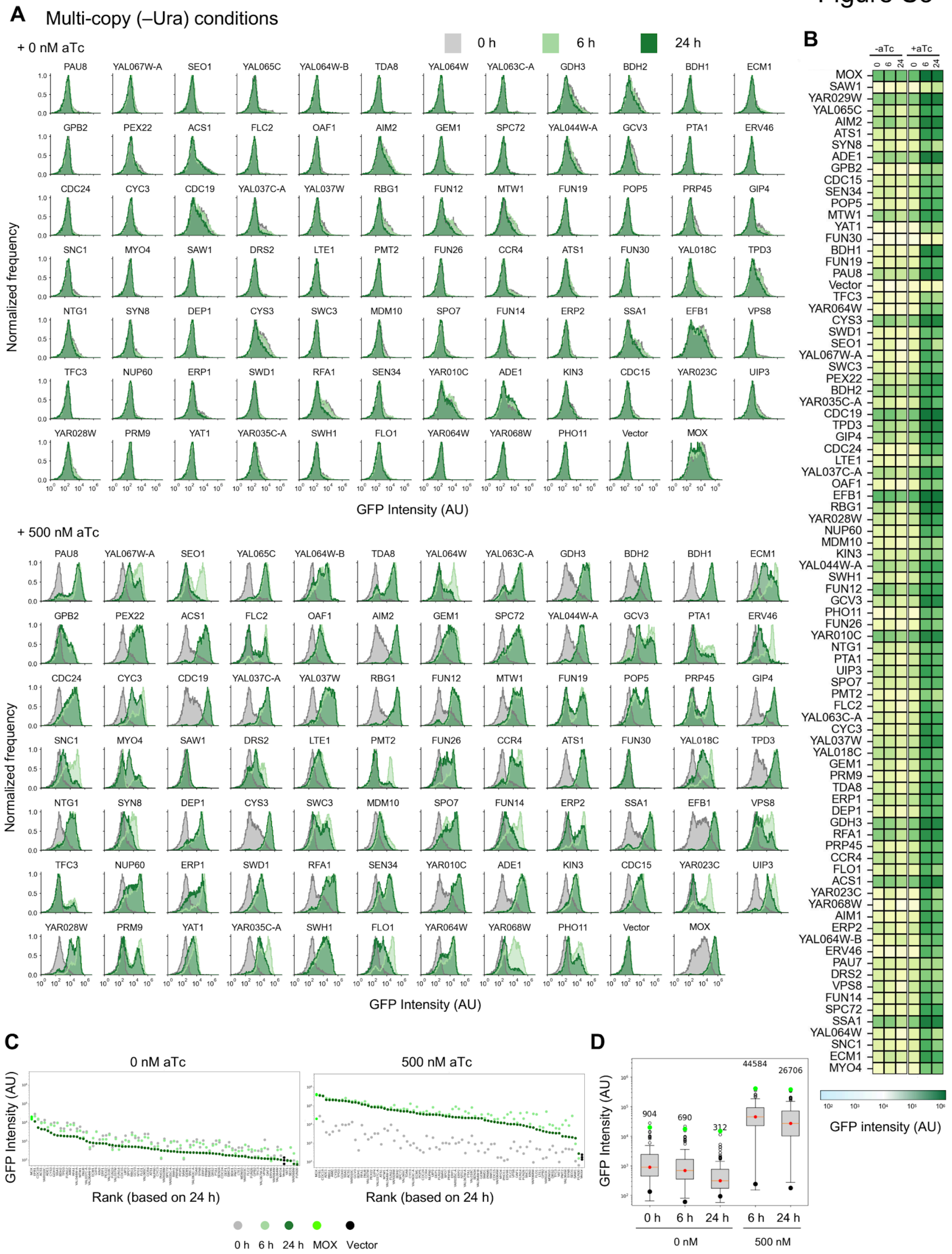


**Figure S3. Protein expression under standard multi-copy conditions.**

**(A)** GFP fluorescence distributions of strains expressing each target protein fused to moxGFP under standard multi-copy (–Ura) conditions. Cells were cultured without aTc (0 nM; top) or with 500 nM aTc (bottom), and GFP fluorescence was measured at 0, 6, and 24 h. Histograms show normalized frequencies of GFP fluorescence intensity (AU).

**(B)** Heatmap of GFP fluorescence intensity for each target protein under the conditions shown in **(A)**. Columns indicate 0, 6, and 24 h without (–aTc) or with (+aTc) 500 nM aTc.

**(C)** GFP fluorescence intensity of each target protein at 0, 6, and 24 h without aTc (left) or with 500 nM aTc (right). Proteins are ranked according to GFP fluorescence intensity at 24 h. MOX and Vector are shown as controls.

**(D)** Distribution of GFP fluorescence intensities across all target proteins under each condition. Numbers above the boxplots indicate the median GFP fluorescence intensity.

Figure S4

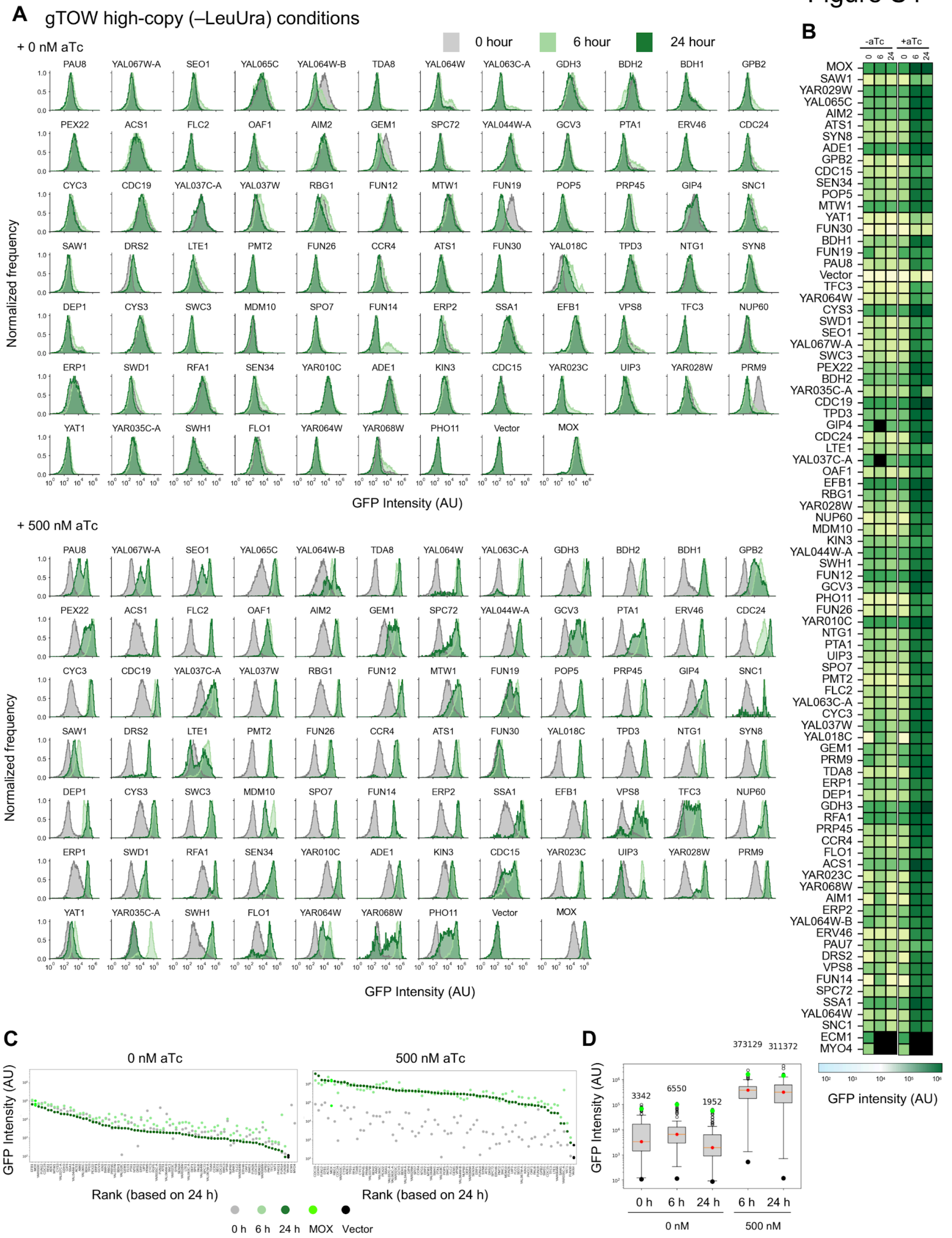


### Figure S4. Protein expression under gTOW high-copy conditions.

**(A)** GFP fluorescence distributions of strains expressing each target protein fused to moxGFP under gTOW high-copy (–LeuUra) conditions. Cells were cultured without aTc (0 nM; top) or with 500 nM aTc (bottom), and GFP fluorescence was measured at 0, 6, and 24 h. Histograms show normalized frequencies of GFP fluorescence intensity (AU).

**(B)** Heatmap of GFP fluorescence intensity for each target protein under the conditions shown in **(A)**. Columns indicate 0, 6, and 24 h without (–aTc) or with (+aTc) 500 nM aTc.

**(C)** GFP fluorescence intensity of each target protein at 0, 6, and 24 h without aTc (left) or with 500 nM aTc (right). Proteins are ranked according to GFP fluorescence intensity at 24 h. MOX and Vector are shown as controls.

**(D)** Distribution of GFP fluorescence intensities across all target proteins under each condition. Numbers above the boxplots indicate the median GFP fluorescence intensity.

Figure S5

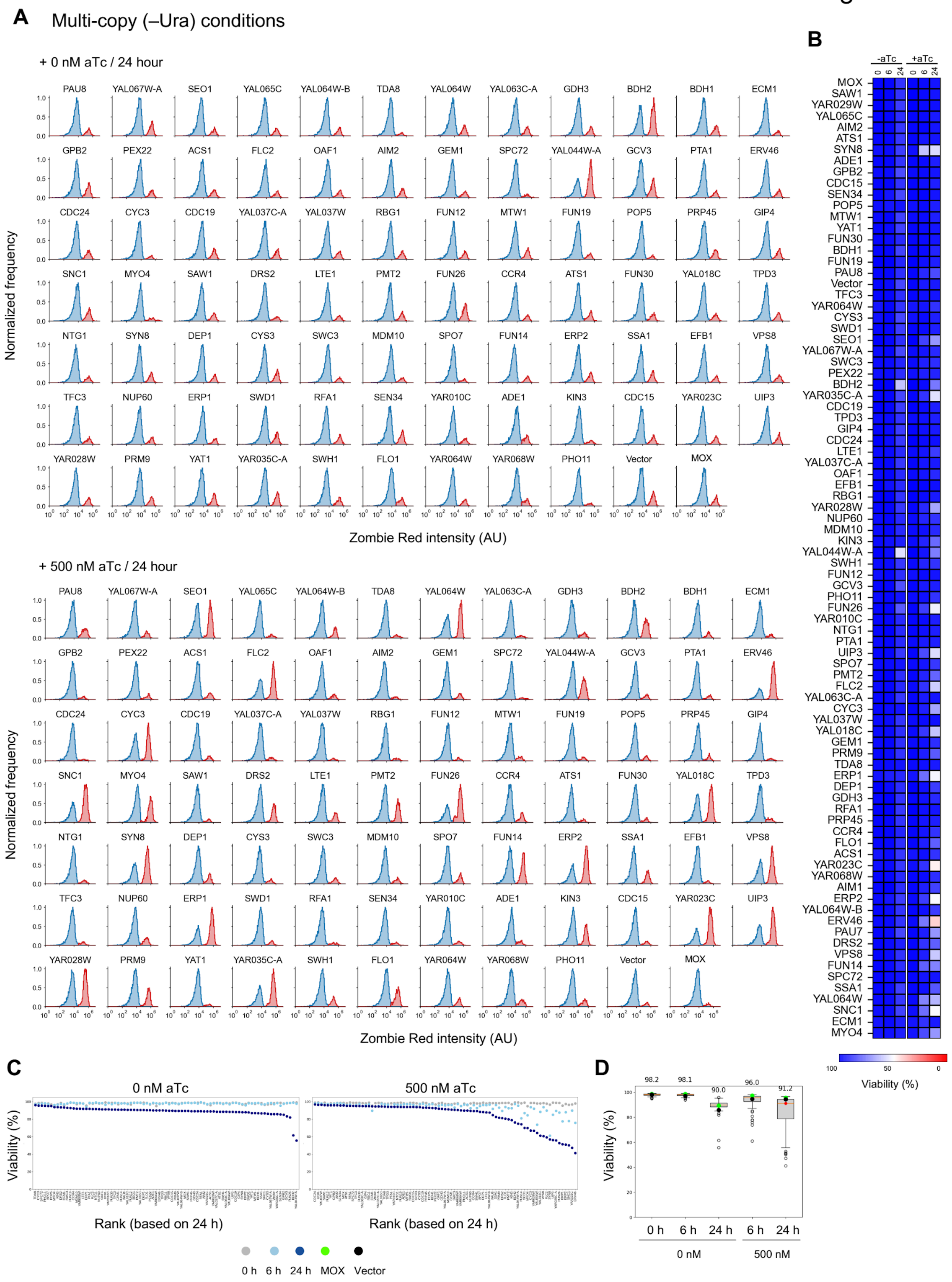


### Figure S5. Cell viability under standard multi-copy conditions.

**(A)** Zombie Red fluorescence distributions of strains expressing each target protein fused to moxGFP under standard multi-copy (–Ura) conditions at 24 h. Cells were cultured without aTc (0 nM; top) or with 500 nM aTc (bottom). Distributions of Zombie Red-negative and Zombie Red-positive cells are shown separately.

**(B)** Heatmap of cell viability for each target protein at 0, 6, and 24 h without (–aTc) or with (+aTc) 500 nM aTc.

**(C)** Cell viability of each strain at 0, 6, and 24 h without aTc (left) or with 500 nM aTc (right). Proteins are ranked according to viability at 24 h. MOX and Vector are shown as controls.

**(D)** Distribution of cell viability across all target proteins under each condition. Numbers above the boxplots indicate the median viability (%).

Figure S6

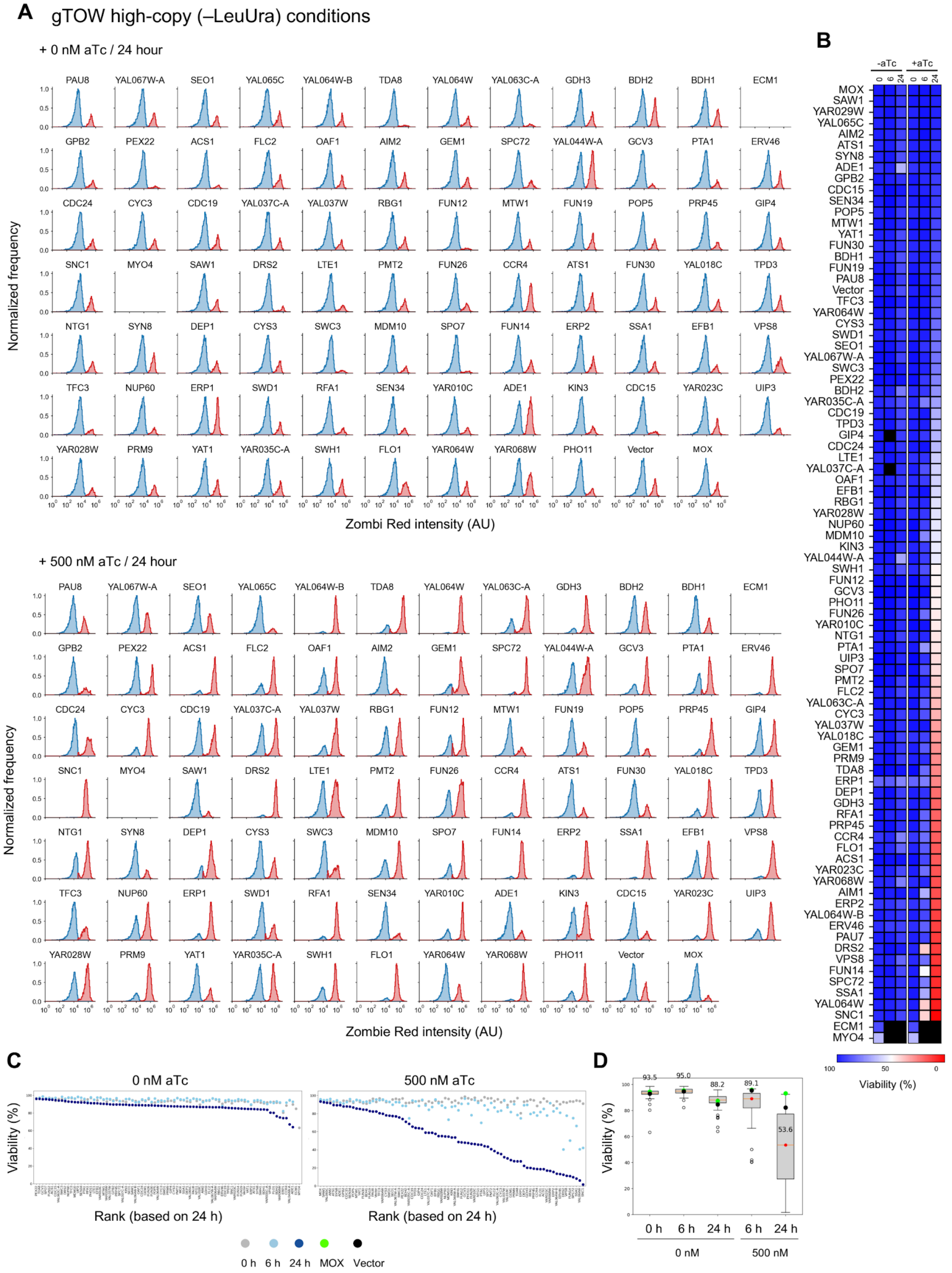


### Figure S6. Cell viability under gTOW high-copy conditions.

**(A)** Zombie Red fluorescence distributions of strains expressing each target protein fused to moxGFP under gTOW high-copy (–LeuUra) conditions at 24 h. Cells were cultured without aTc (0 nM; top) or with 500 nM aTc (bottom). Distributions of Zombie Red-negative and Zombie Red-positive cells are shown separately.

**(B)** Heatmap of cell viability for each target protein at 0, 6, and 24 h without (–aTc) or with (+aTc) 500 nM aTc.

**(C)** Cell viability of each strain at 0, 6, and 24 h without aTc (left) or with 500 nM aTc (right). Proteins are ranked according to viability at 24 h. MOX and Vector are shown as controls.

**(D)** Distribution of cell viability across all target proteins under each condition. Numbers above the boxplots indicate the median viability (%).

Figure S7

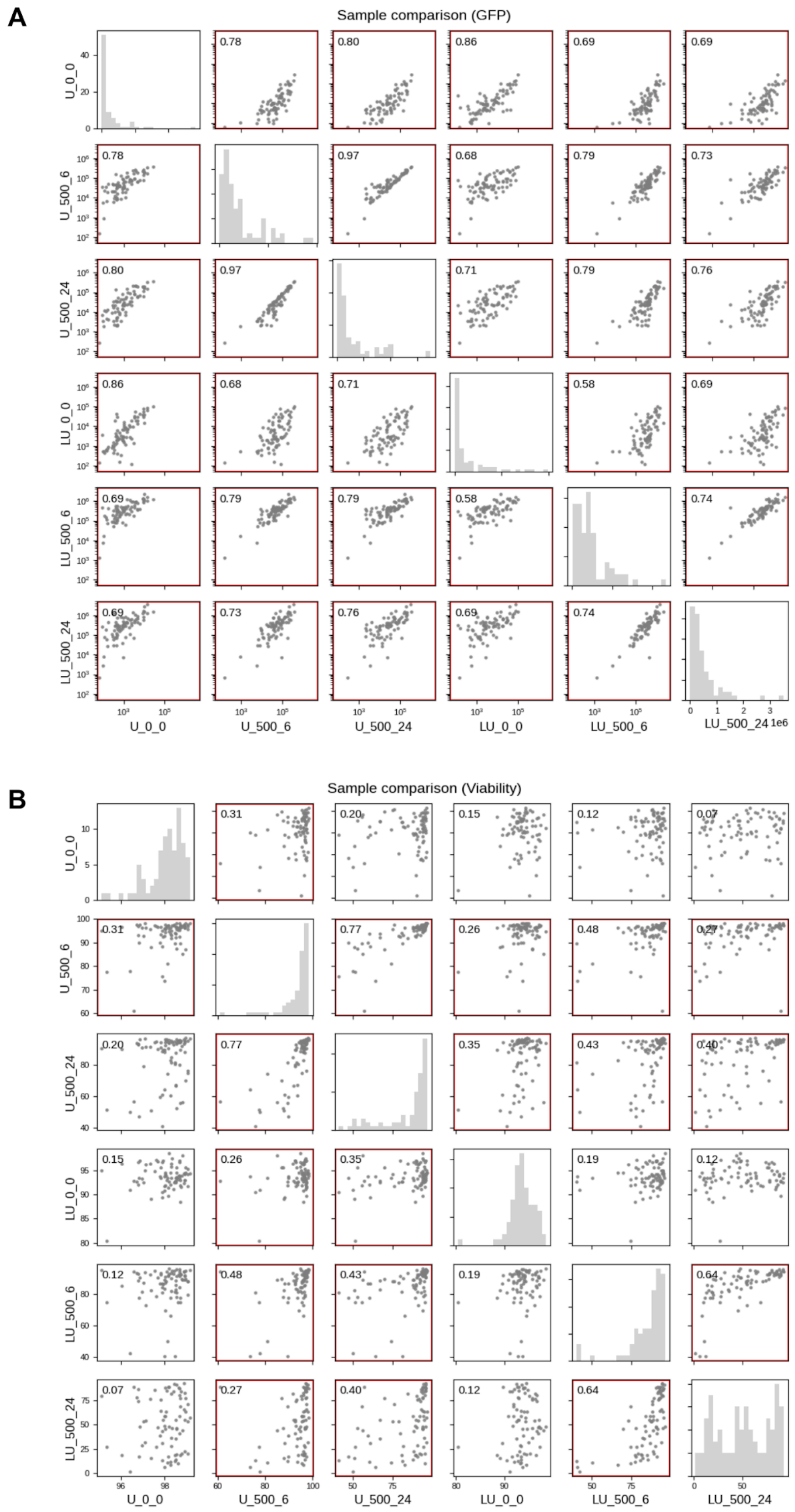


**Figure S7. Pairwise comparison of GFP fluorescence and cell viability across culture conditions.**

**(A)** Pairwise comparison of GFP fluorescence intensity across standard multi-copy (U) and gTOW high-copy (LU) conditions at the indicated aTc concentrations and time points. Each point represents a target protein. Histograms on the diagonal show the distribution of GFP fluorescence intensities for each condition. Numbers indicate Pearson correlation coefficients between conditions. Framed panels indicate significant correlations (Benjamini–Hochberg-adjusted $p < 0.05$).

**(B)** Pairwise comparison of cell viability across the same conditions. Each point represents a target protein, and diagonal histograms show the distribution of viability values. Numbers indicate Pearson correlation coefficients between conditions. Framed panels indicate significant correlations (Benjamini–Hochberg-adjusted $p < 0.05$). U and LU denote standard multi-copy (–Ura) and gTOW high-copy (–LeuUra) conditions, respectively; numbers following U or LU indicate the aTc concentration (0 or 500 nM) and sampling time (0, 6, or 24 h).

Figure S8

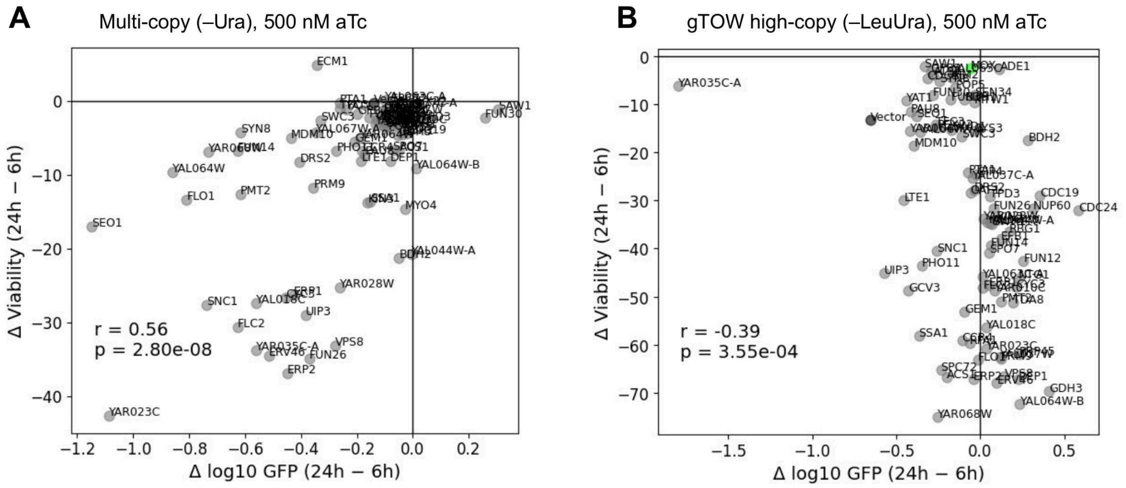


#### Figure S8. Relationship between changes in protein expression and cell viability from 6 to 24 h.

Changes in GFP fluorescence intensity and cell viability between 6 and 24 h after induction with 500 nM aTc are shown for each target protein under **(A)** standard multi-copy (–Ura) and **(B)** gTOW high-copy (–LeuUra) conditions. The *x*-axis indicates the change in log10 GFP fluorescence intensity (24 h − 6 h), and the *y*-axis indicates the change in cell viability (24 h − 6 h). Each point represents a target protein. Pearson correlation coefficients (*r*) and *p*-values are shown in each panel.

Figure S9

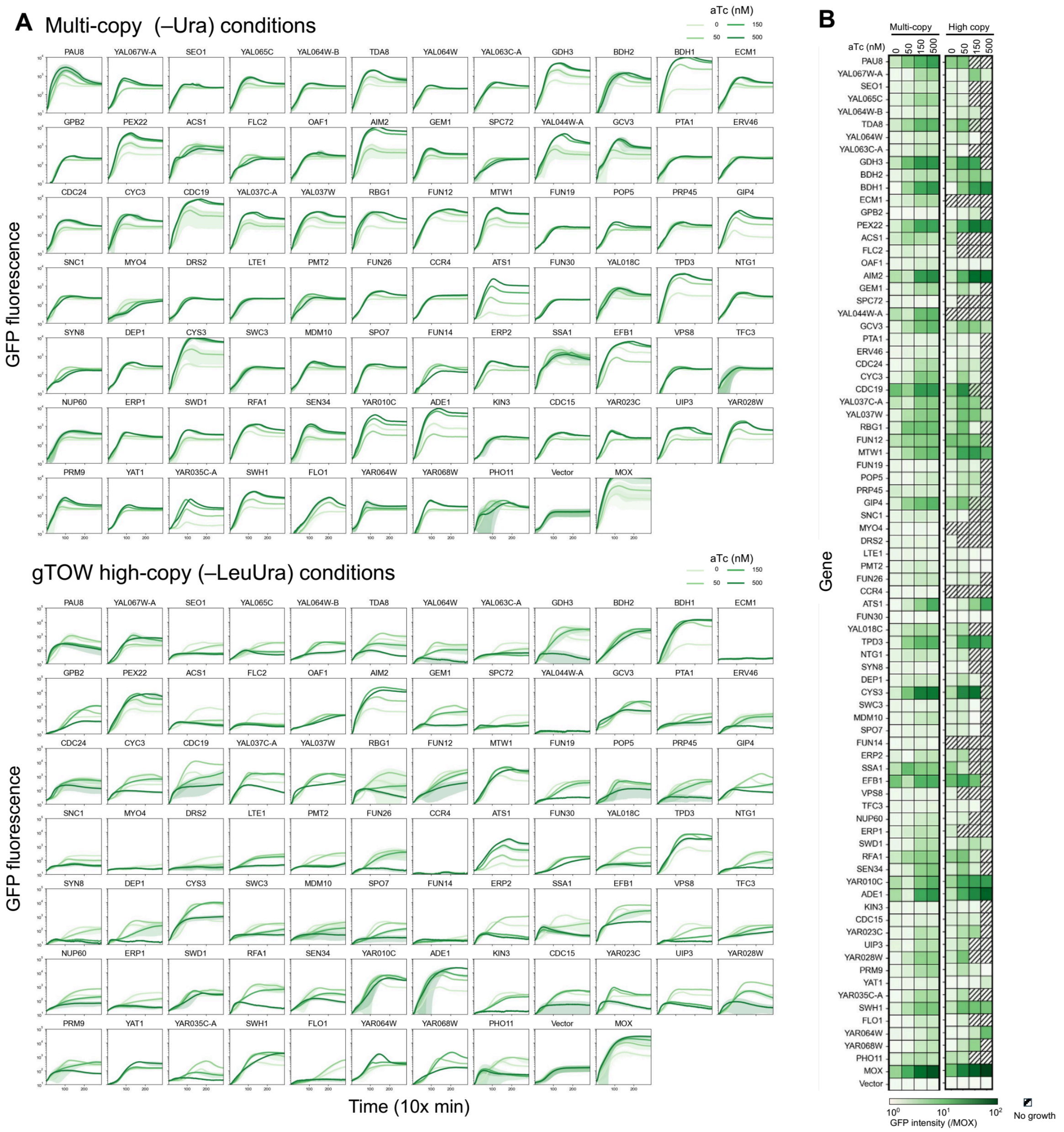

#### Figure S9. Time-course analysis of protein expression under different induction conditions.

**(A)** Time-course profiles of GFP fluorescence for each target protein under standard multi-copy (–Ura; top) and gTOW high-copy (–LeuUra; bottom) conditions. Cells were cultured with 0, 50, 150, or 500 nM aTc, and GFP fluorescence was measured at 10-min intervals using a plate reader. MOX and Vector are shown as controls. Lines indicate mean values, and shaded bands represent ± SD.

**(B)** Heatmap of maximum GFP fluorescence levels obtained from the time-course measurements shown in **(A)**. Columns indicate aTc concentrations (0, 50, 150, and 500 nM) under standard multi-copy and gTOW high-copy conditions. GFP fluorescence is expressed relative to MOX. Hatched cells indicate conditions in which no growth was detected.

Figure S10

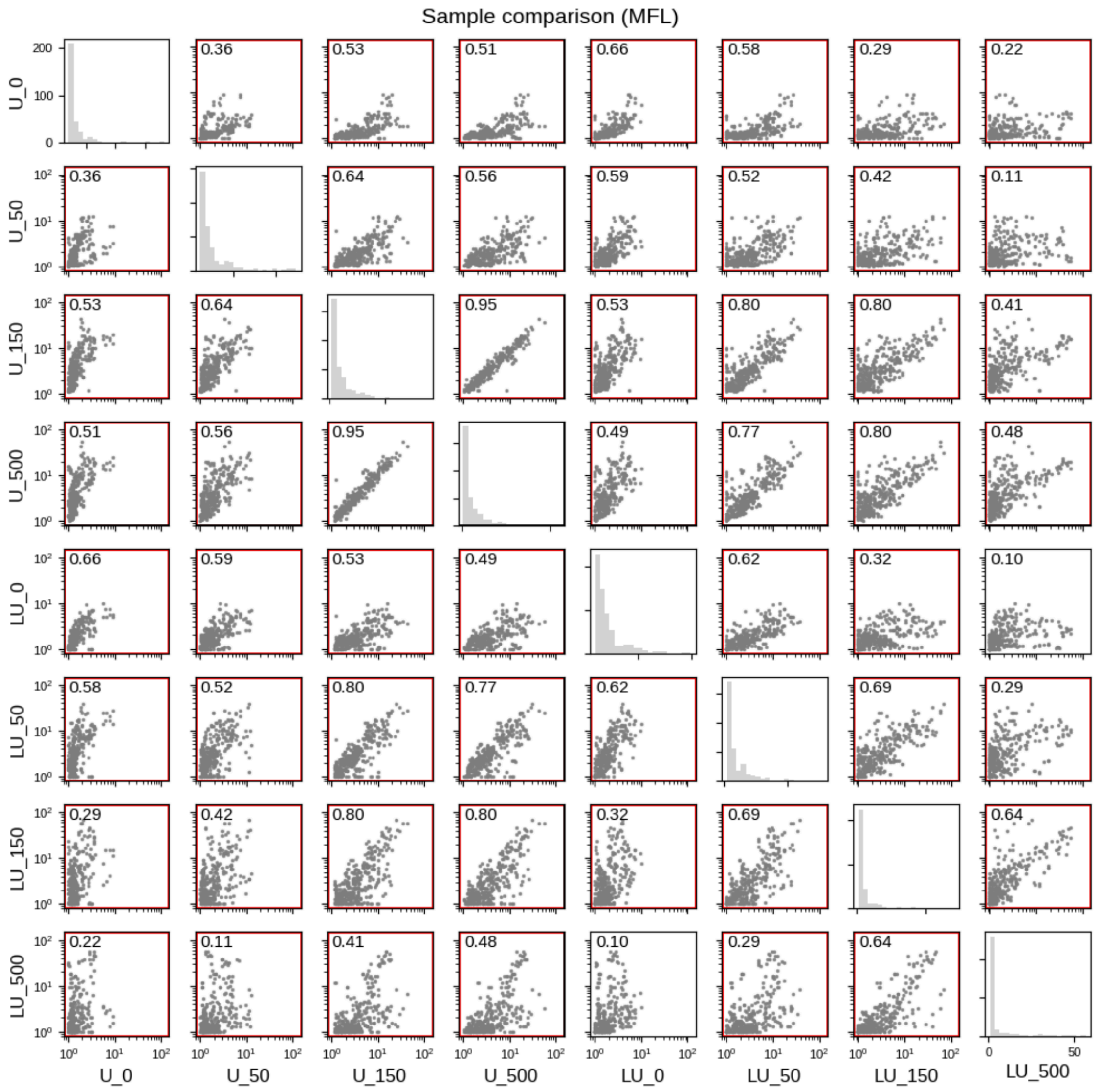


**Figure S10. Pairwise comparison of maximum fluorescence levels across induction conditions.**

Pairwise comparison of maximum fluorescence levels (MFL) for each target protein under standard multi-copy (U) and gTOW high-copy (LU) conditions at 0, 50, 150, and 500 nM aTc. Each point represents a target protein. Histograms on the diagonal show the distribution of MFL values for each condition. Numbers indicate Pearson correlation coefficients between conditions. U and LU denote standard multi-copy (–Ura) and gTOW high-copy (–LeuUra) conditions, respectively. Framed panels indicate significant correlations (Benjamini–Hochberg-adjusted $p < 0.05$).

Figure S11

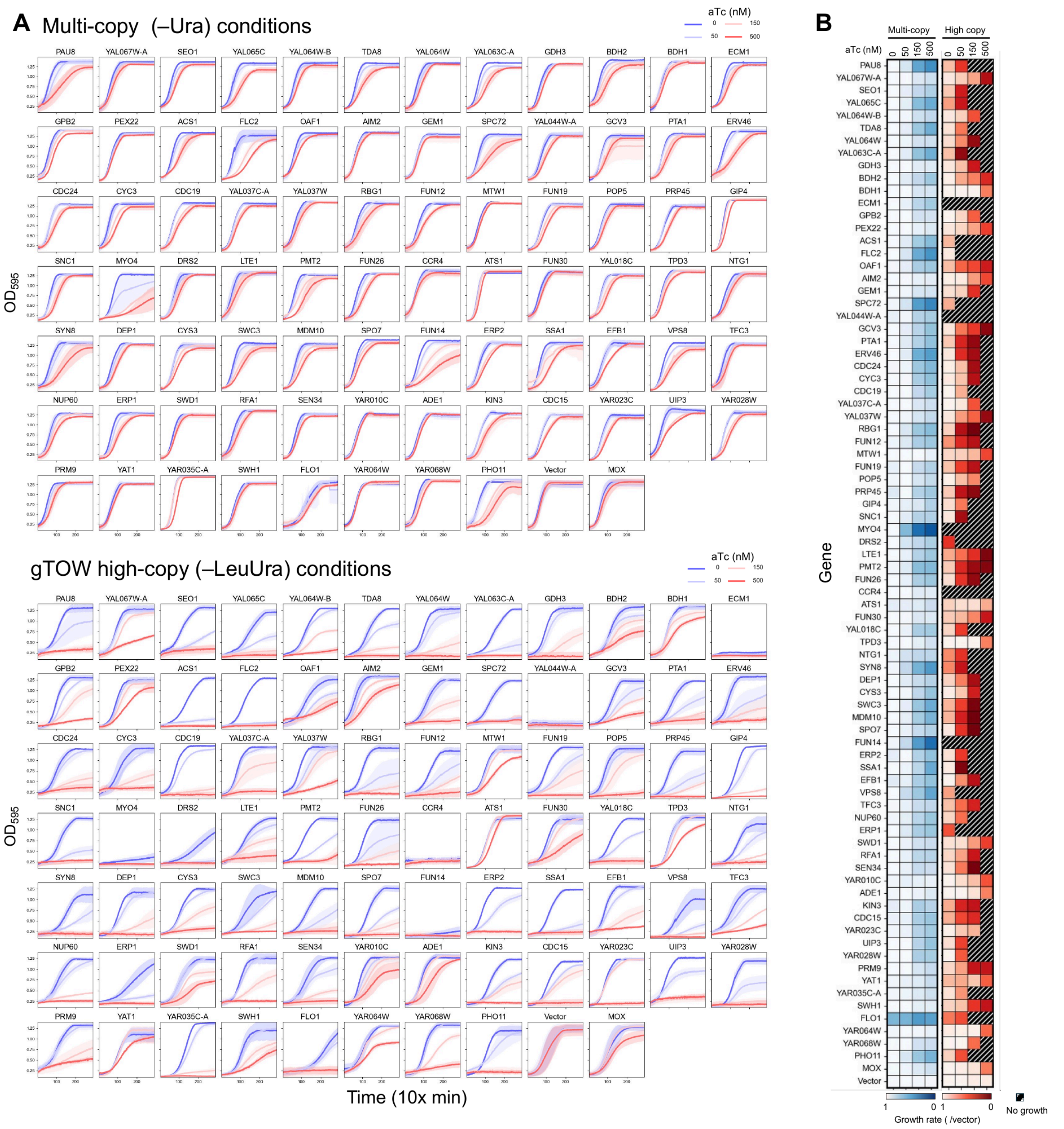


### Figure S11. Time-course analysis of growth under different induction conditions.

**(A)** Time-course profiles of growth (OD595) for each target protein under standard multi-copy (–Ura; top) and gTOW high-copy (–LeuUra; bottom) conditions. Cells were cultured with 0, 50, 150, or 500 nM aTc, and OD595 was measured at 10-min intervals using a plate reader. MOX and Vector are shown as controls. Lines indicate mean values, and shaded bands represent ± SD.

**(B)** Heatmap of maximum growth rates obtained from the time-course measurements shown in **(A)**. Columns indicate aTc concentrations (0, 50, 150, and 500 nM) under standard multi-copy and gTOW high-copy conditions. Growth rates are expressed relative to Vector. Hatched cells indicate conditions in which no growth was detected.

Figure S12

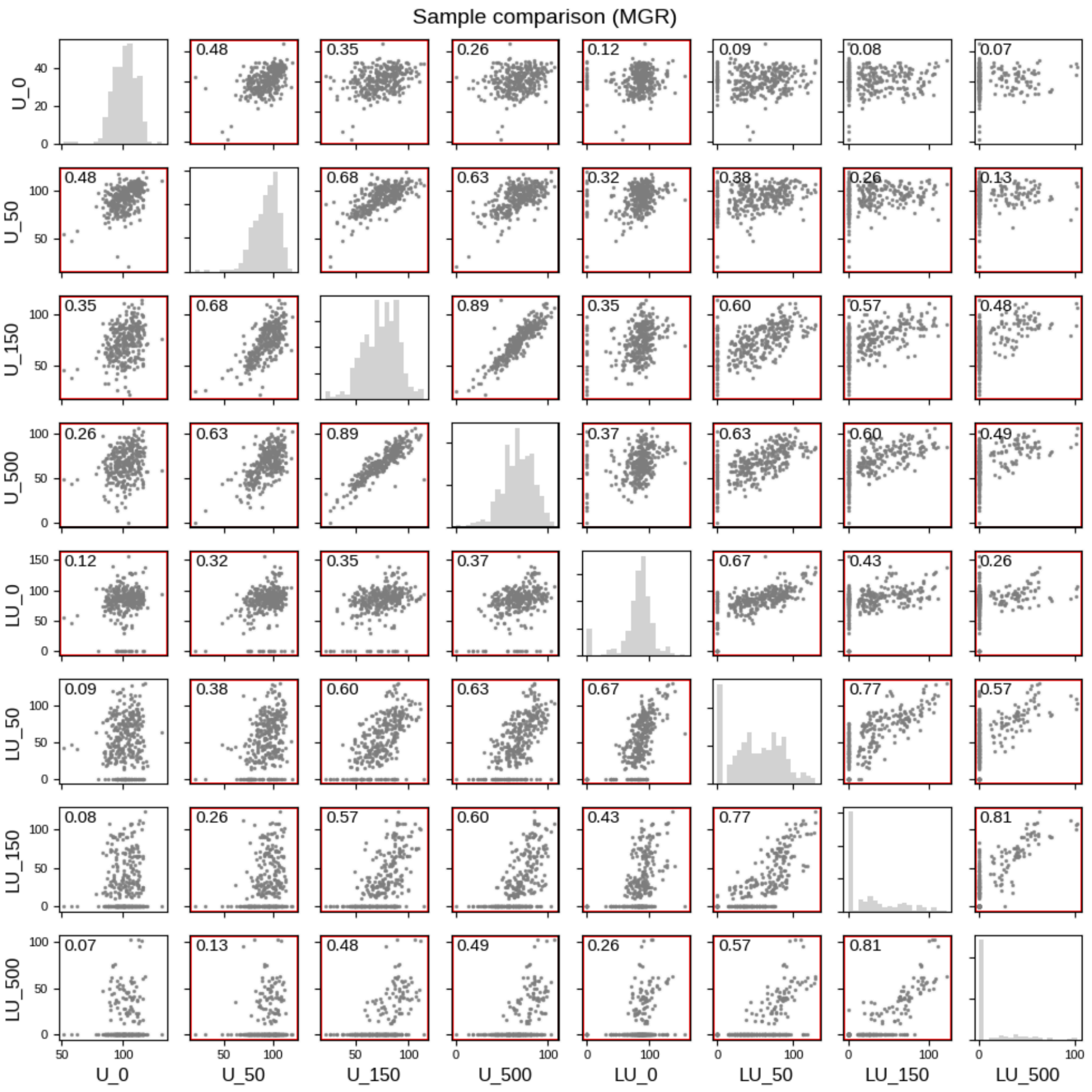


#### Figure S12. Pairwise comparison of maximum growth rates across induction conditions.

Pairwise comparison of maximum growth rates (MGR) for each target protein under standard multi-copy (U) and gTOW high-copy (LU) conditions at 0, 50, 150, and 500 nM aTc. Each point represents a target protein. Histograms on the diagonal show the distribution of MGR values for each condition. Numbers indicate Pearson correlation coefficients between conditions. U and LU denote standard multi-copy (–Ura) and gTOW high-copy (–LeuUra) conditions, respectively. Framed panels indicate significant correlations (Benjamini–Hochberg-adjusted $p < 0.05$).

Figure S13

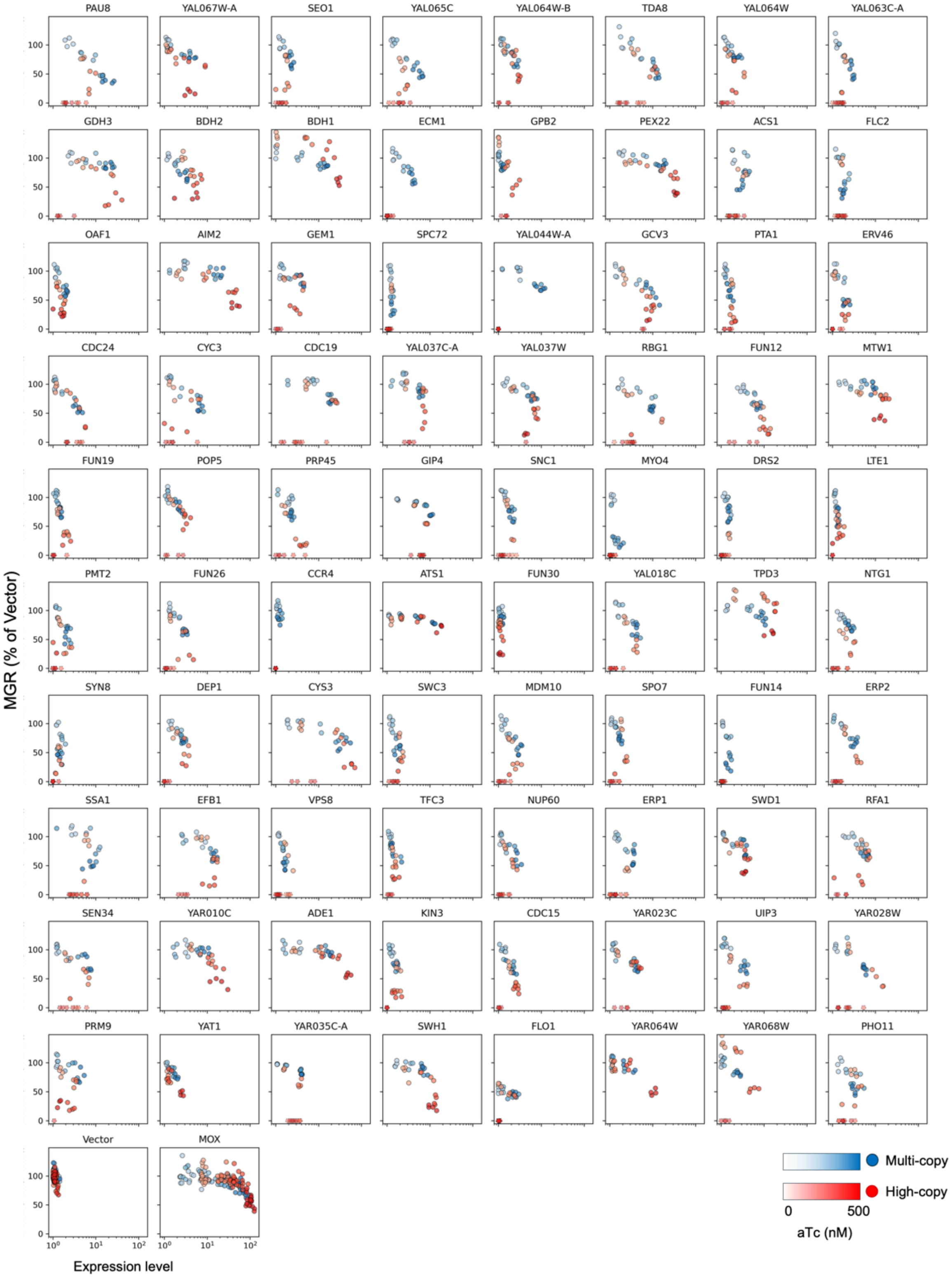
PAU8
YAL067W-A
SEO1
YAL065C
YAL064W-B
TDA8
YAL064W
YAL063C-A
GDH3
BDH2
BDH1
ECM1
GPB2
PEX22
ACS1
FLC2
OAF1
AIM2
GEM1
SPC72
YAL044W-A
GCV3
PTA1
ERV46
CDC24
CYC3
CDC19
YAL037C-A
YAL037W
RBG1
FUN12
MTW1
FUN19
POP5
PRP45
GIP4
SNC1
MYO4
DRS2
LTE1
PMT2
FUN26
CCR4
ATS1
FUN30
YAL018C
TPD3
NTG1
SYN8
DEP1
CYS3
SWC3
MDM10
SPO7
FUN14
ERP2
SSA1
EFB1
VPS8
TFC3
NUP60
ERP1
SWD1
RFA1
SEN34
YAR010C
ADE1
KIN3
CDC15
YAR023C
UIP3
YAR028W
PRM9
YAT1
YAR035C-A
SWH1
FLO1
YAR064W
YAR068W
PHO11
Vector
MOX
MGR (% of Vector)
Expression level
Multi-copy
High-copy
0
500
aTc (nM)

#### Figure S13. Relationship between protein expression level and growth rate for individual target proteins.

Relationship between protein expression level and growth rate for each target protein under standard multi-copy (–Ura) and gTOW high-copy (–LeuUra) conditions. Each point represents a measurement at one of four aTc concentrations (0, 50, 150, or 500 nM). Expression levels are shown on a log10 scale, and growth rates are expressed relative to Vector. MOX and Vector are shown as controls.

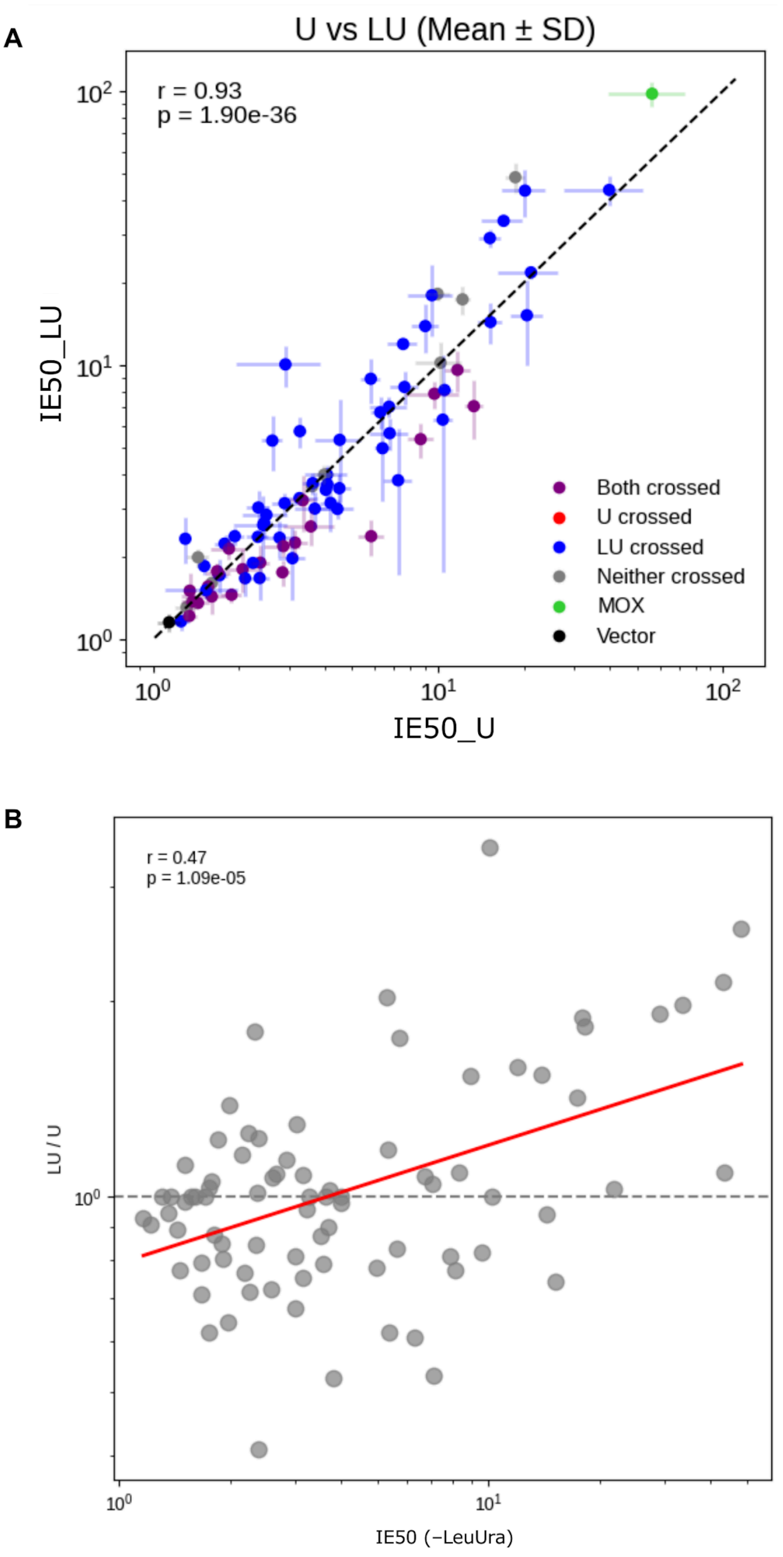


**Figure S14. Comparison of expression limits between standard multi-copy and gTOW high-copy conditions.**

**(A)** Comparison of IE50 values obtained under standard multi-copy (U) and gTOW high-copy (LU) conditions. Points are classified according to whether MGR crossed 50% under either condition. Error bars indicate mean ± SD. The dashed line indicates equality between the two conditions. Pearson correlation coefficient (*r*) and *p*-value are shown.

**(B)** Relationship between IE50 under gTOW high-copy conditions and the ratio of IE50 between gTOW high-copy and standard multi-copy conditions (LU/U). The linear regression is shown, and the dashed line indicates LU/U = 1. Pearson correlation coefficient (*r*) and *p*-value are shown.

Figure S15

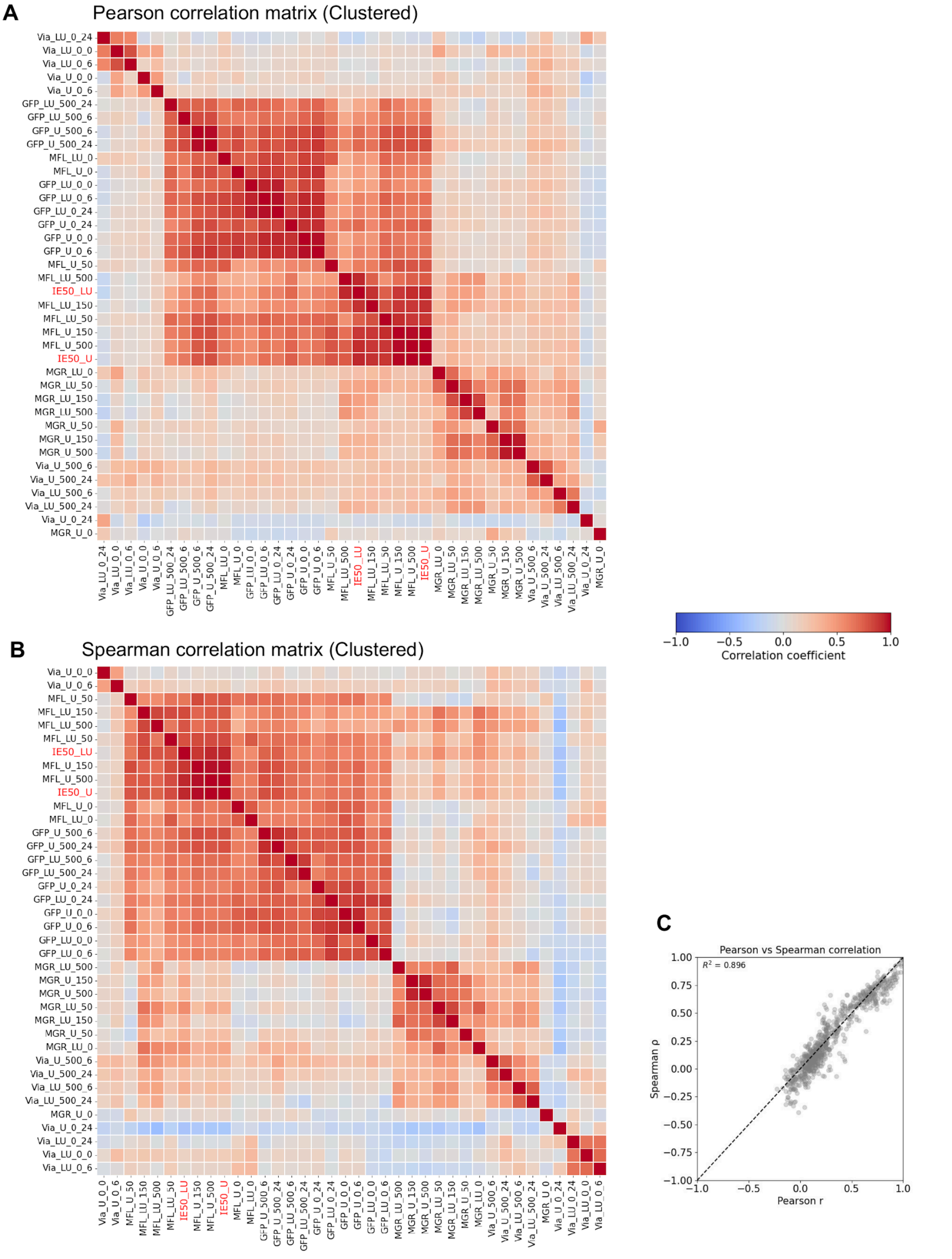
A
Pearson correlation matrix (Clustered)
Via_LU_0_24
Via_LU_0_0
Via_LU_0_6
Via_U_0_0
Via_U_0_6
GFP_LU_500_24
GFP_LU_500_6
GFP_U_500_6
GFP_U_500_24
MFL_LU_0
MFL_U_0
GFP_LU_0_0
GFP_LU_0_6
GFP_LU_0_24
GFP_U_0_24
GFP_U_0_0
GFP_U_0_6
MFL_U_50
MFL_LU_500
IE50_LU
MFL_LU_150
MFL_LU_50
MFL_U_150
MFL_U_500
IE50_U
MGR_LU_0
MGR_LU_50
MGR_LU_150
MGR_LU_500
MGR_U_50
MGR_U_150
MGR_U_500
Via_U_500_6
Via_U_500_24
Via_LU_500_6
Via_LU_500_24
Via_U_0_24
MGR_U_0
−1.0
−0.5
0.0
0.5
1.0
Correlation coefficient
B
Spearman correlation matrix (Clustered)
Via_U_0_0
Via_U_0_6
MFL_U_50
MFL_LU_150
MFL_LU_500
MFL_LU_50
IE50_LU
MFL_U_150
MFL_U_500
IE50_U
MFL_U_0
MFL_LU_0
GFP_U_500_6
GFP_U_500_24
GFP_LU_500_6
GFP_LU_500_24
GFP_U_0_24
GFP_LU_0_24
GFP_U_0_0
GFP_U_0_6
GFP_LU_0_0
GFP_LU_0_6
MGR_LU_500
MGR_U_150
MGR_U_500
MGR_LU_50
MGR_LU_150
MGR_U_50
MGR_LU_0
Via_U_500_6
Via_U_500_24
Via_LU_500_6
Via_LU_500_24
MGR_U_0
Via_U_0_24
Via_LU_0_24
Via_LU_0_0
Via_LU_0_6
C
Pearson vs Spearman correlation
$R^2$ = 0.896
Spearman ρ
Pearson r

#### Figure S15. Correlation structure among expression, growth, viability, and expression-limit measurements.

**(A, B)** Clustered correlation matrices of the measured parameters using Pearson **(A)** and Spearman **(B)** correlation coefficients. Parameters include GFP fluorescence, maximum fluorescence level (MFL), maximum growth rate (MGR), viability, and IE50 under standard multi-copy (U) and gTOW high-copy (LU) conditions. Matrix entries represent correlation coefficients ranging from –1 to 1.

**(C)** Comparison of Pearson and Spearman correlation coefficients for all parameter pairs shown in **(A)** and **(B)**. The dashed line indicates equality between the two coefficients. $R^2$ is shown in the panel.

Figure S16

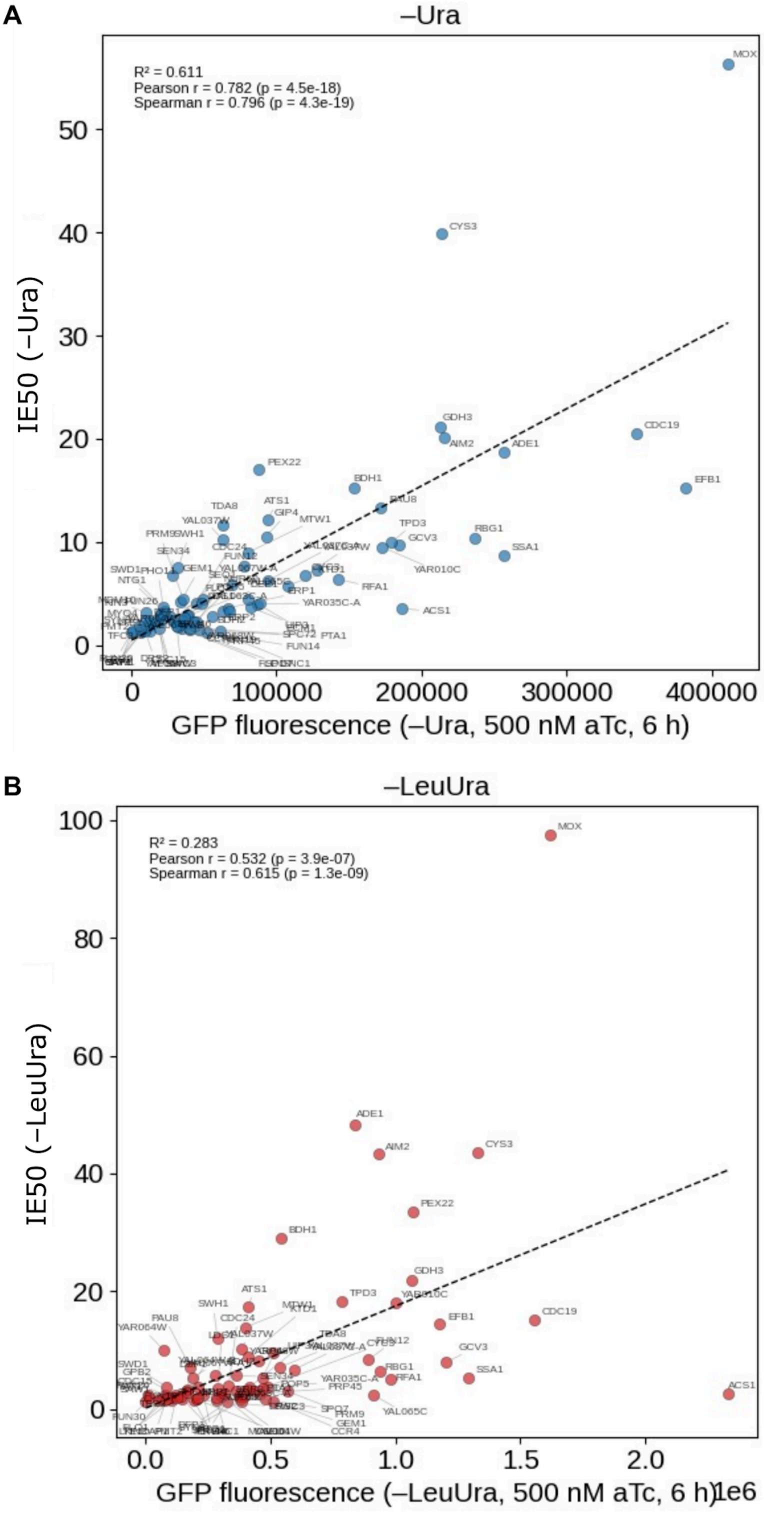
A
–Ura
R² = 0.611
Pearson r = 0.782 (p = 4.5e-18)
Spearman r = 0.796 (p = 4.3e-19)
IE50 (–Ura)
GFP fluorescence (–Ura, 500 nM aTc, 6 h)
MOX
CYS3
GDH3
AIM2
ADE1
CDC19
EFB1
PEX22
BDH1
PAU8
TPD3
GCV3
RBG1
SSA1
YAR010C
ACS1
RFA1
YAR035C-A
PTA1
FUN14
B
–LeuUra
R² = 0.283
Pearson r = 0.532 (p = 3.9e-07)
Spearman r = 0.615 (p = 1.3e-09)
IE50 (–LeuUra)
GFP fluorescence (–LeuUra, 500 nM aTc, 6 h)
1e6
MOX
ADE1
AIM2
CYS3
PEX22
BDH1
GDH3
TPD3
ATS1
EFB1
CDC19
GCV3
SSA1
ACS1
RBG1
RFA1
YAL065C

#### Figure S16. Relationship between IE50 and GFP fluorescence measured by flow cytometry.

Relationship between IE50 and GFP fluorescence measured at 6 h after induction with 500 nM aTc under **(A)** standard multi-copy (–Ura) and **(B)** gTOW high-copy (–LeuUra) conditions. Each point represents a target protein. Dashed lines indicate linear regression. $R^2$, Pearson and Spearman correlation coefficients, and *p*-values are shown in each panel.

Figure S17

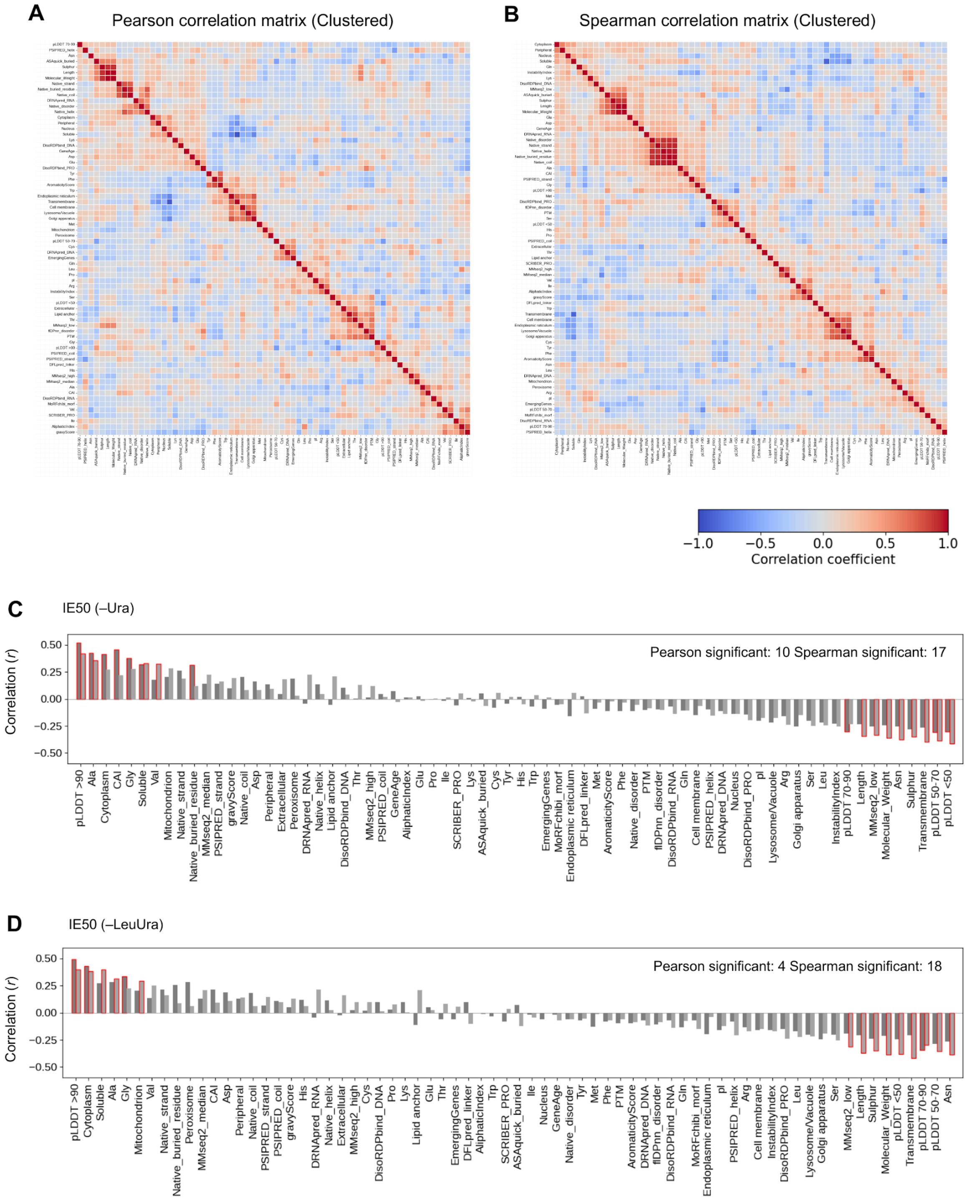


**Figure S17. Correlations among protein parameters and their relationships with IE50.**

**(A, B)** Clustered correlation matrices of approximately 70 protein parameters analyzed in this study using Pearson **(A)** and Spearman **(B)** correlation coefficients. Matrix entries represent correlation coefficients ranging from –1 to 1.

**(C, D)** Correlations between each protein parameter and IE50 under standard multi-copy (–Ura; **C**) and gTOW high-copy (–LeuUra; **D**) conditions. Pearson and Spearman correlation coefficients are shown for each parameter. Parameters are ordered according to their Pearson correlation coefficients. Outlined bars indicate significant correlations (FDR < 0.05, Benjamini–Hochberg correction). The numbers of parameters showing significant Pearson and Spearman correlations are indicated in each panel.

Figure S18

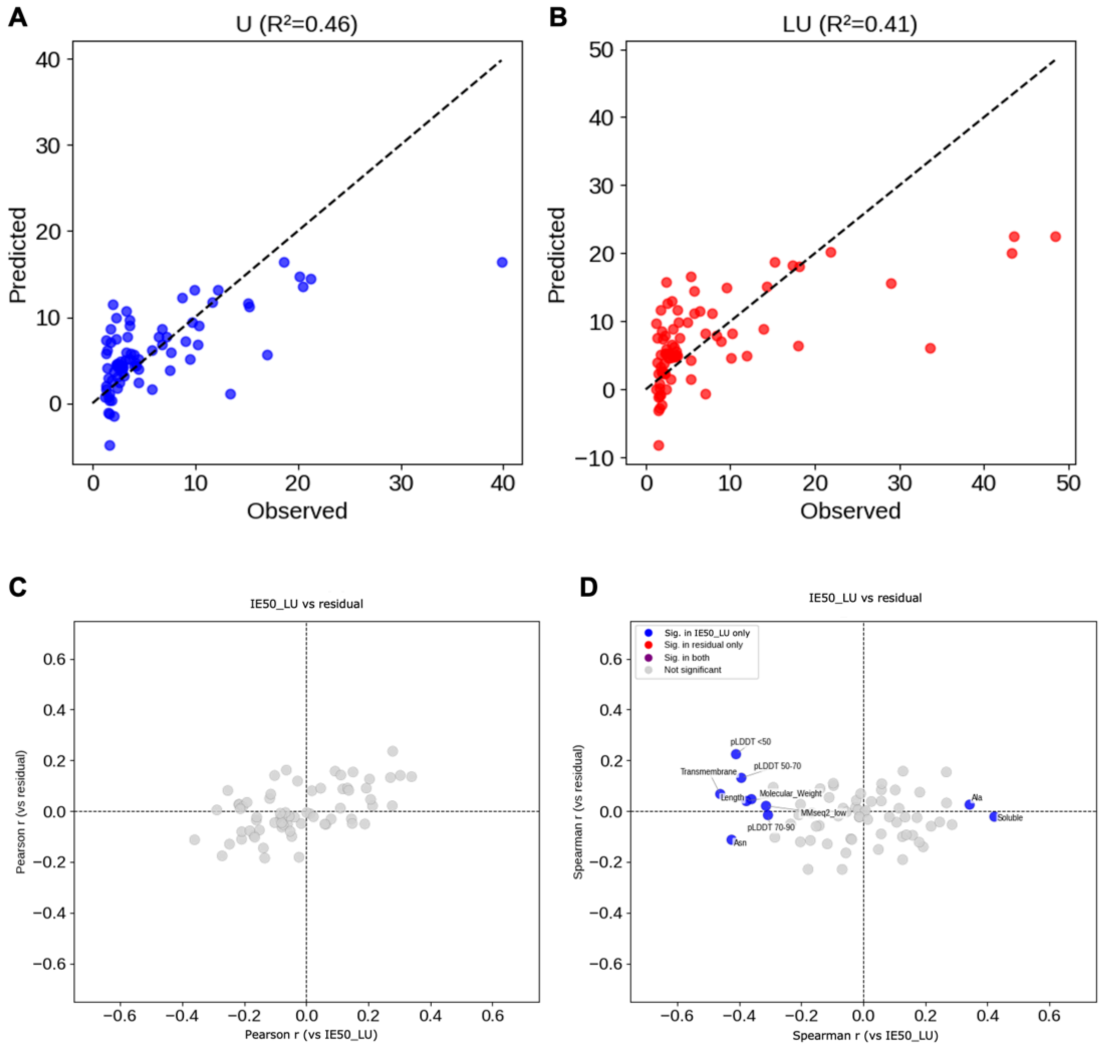


#### Figure S18. Prediction of IE50 using Cytoplasm score, pLDDT > 90, and Sulphur content.

**(A, B)** Relationship between observed IE50 values and values predicted by a multiple regression model using Cytoplasm score, pLDDT > 90, and Sulphur content under standard multi-copy (–Ura; **A**) and gTOW high-copy (–LeuUra; **B**) conditions. Dashed lines indicate equality between observed and predicted values. $R^2$ values are shown above each panel. $R^2$ was calculated using the same data used to fit the model.

**(C, D)** Relationship between the correlation of each protein parameter with IE50 under –LeuUra conditions and its correlation with the residuals from the three-parameter regression model. Pearson **(C)** and Spearman **(D)** correlation coefficients are shown. Each point represents a protein parameter. Dashed lines indicate zero correlation. Parameters are classified according to whether they show significant correlations with IE50 under –LeuUra conditions only, with the regression residual only, with both, or with neither. For this classification, correlations with Benjamini–Hochberg-adjusted *p*-values $< 0.10$ were classified as significant.

Figure S19

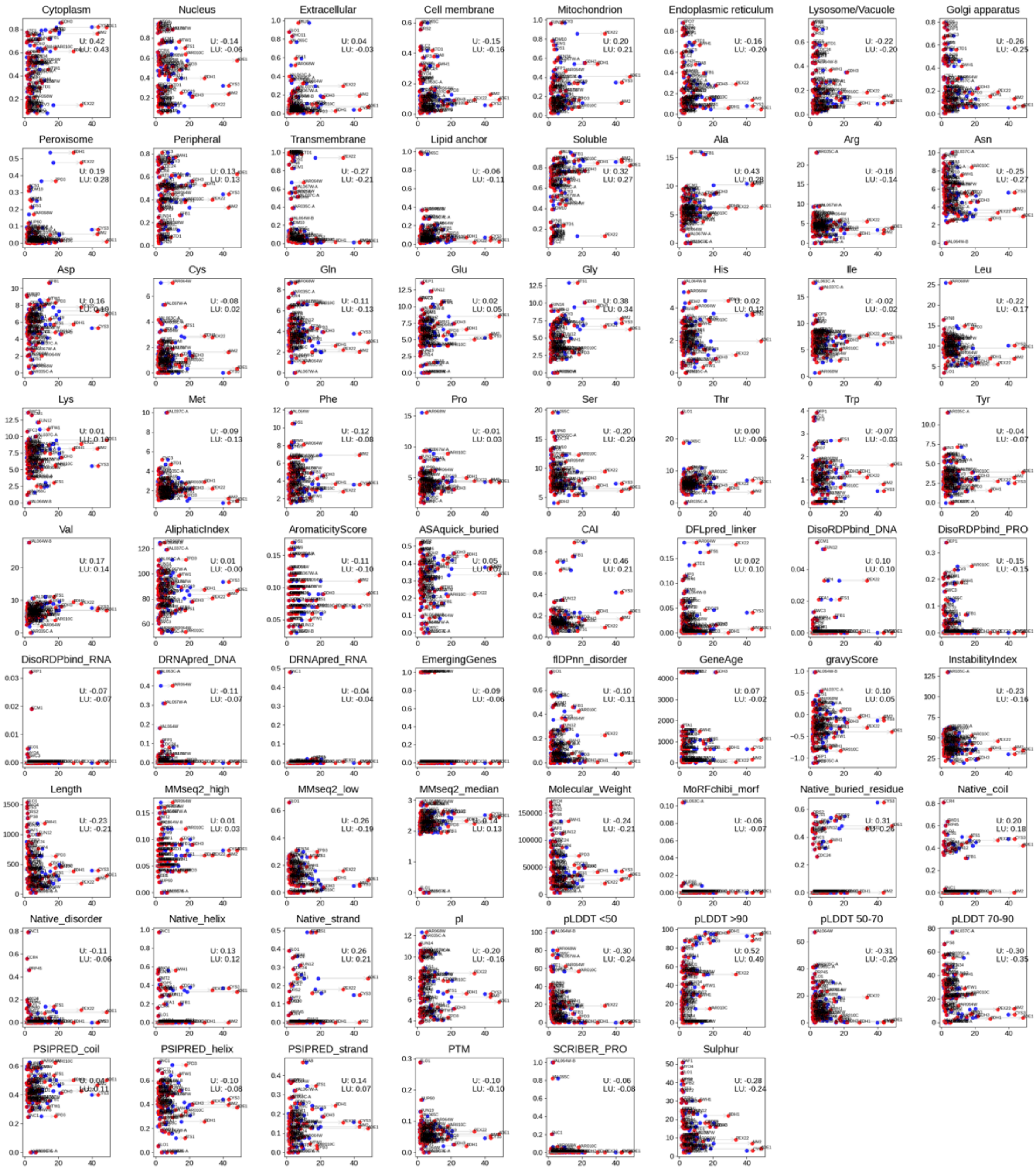


#### Figure S19. Relationships between individual protein parameters and IE50.

Scatter plots showing the relationships between IE50 (horizontal axis) and individual protein parameters (vertical axis). Data are shown for standard multi-copy (U; –Ura) and gTOW high-copy (LU; –LeuUra) conditions. Each point represents a target protein, with the corresponding gene name indicated, and values for the same gene are connected by lines. The parameter analyzed is shown above each panel. Pearson correlation coefficients for each condition are indicated as U and LU. Parameters include subcellular localization scores, amino acid composition, physicochemical properties, predicted structural features, and other gene- and protein-level annotations.

Figure S20

**A** Pearson correlation matrix (Clustered)

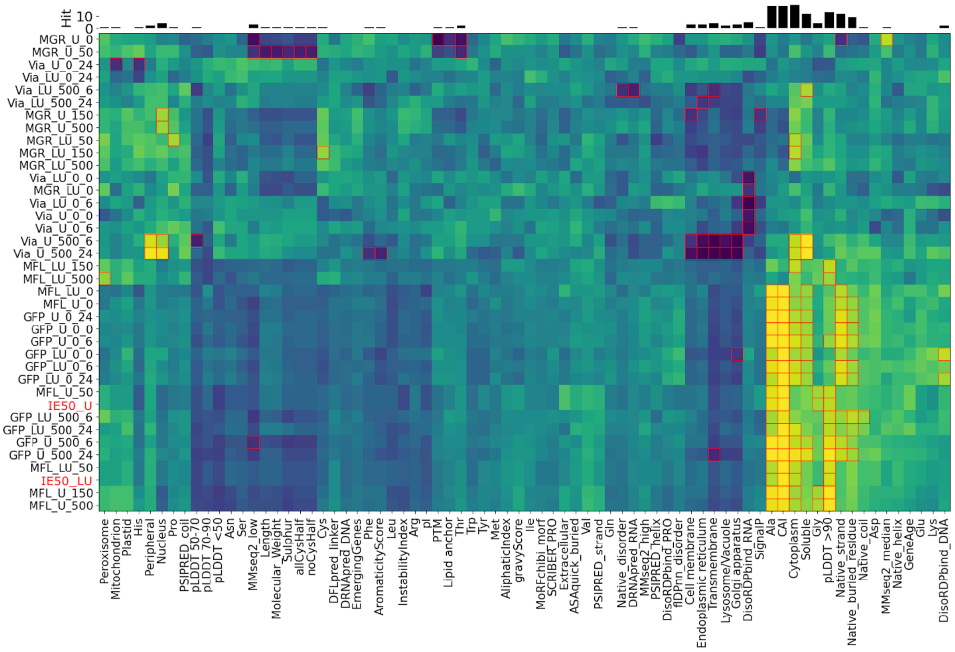


**B** Spearman correlation matrix (Clustered)

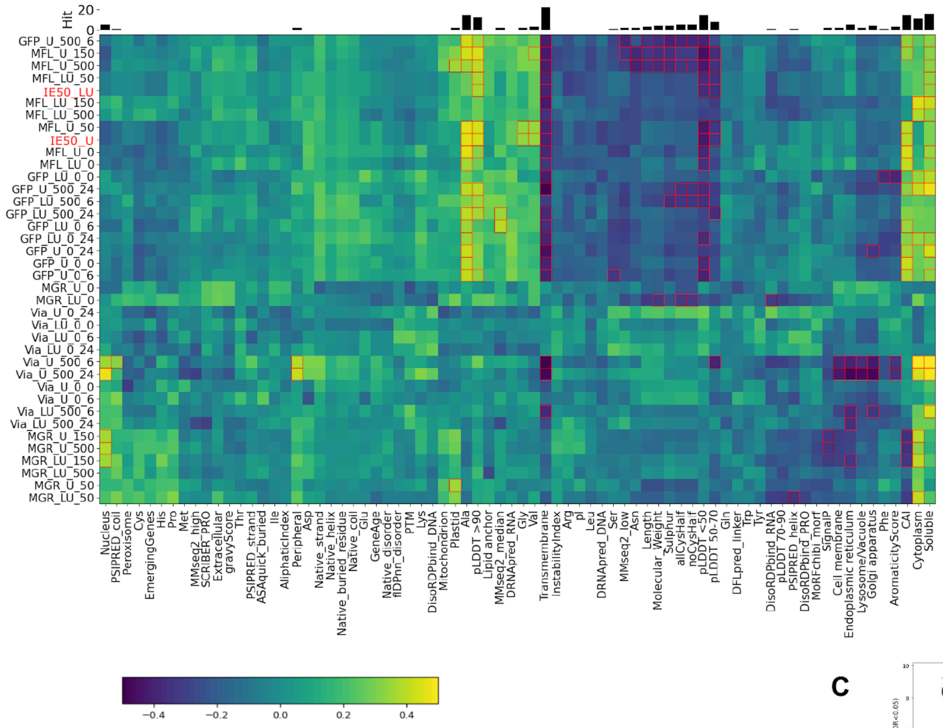


**C**

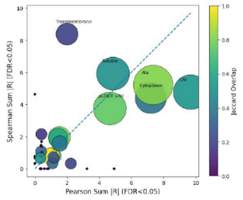

#### Figure S20. Correlations between protein properties and experimental measurements of expression, growth, viability, and expression limits.

**(A, B)** Correlation matrices between protein parameters and experimental measurements using Pearson **(A)** and Spearman **(B)** correlation coefficients. Rows indicate experimental measurements, including GFP fluorescence measured by flow cytometry, maximum fluorescence level (MFL), maximum growth rate (MGR), cell viability, and IE50 under standard multi-copy (U) and gTOW high-copy (LU) conditions. Columns indicate the protein parameters analyzed in **Figure 3**. Matrix entries represent correlation coefficients. Outlines indicate statistically significant correlations after multiple-testing correction. Bars above the matrices indicate the number of significant correlations for each protein parameter.

**(C)** Comparison of the sums of the absolute values of significant correlation coefficients (FDR < 0.05) obtained using Pearson and Spearman correlation analyses. Each point represents a protein parameter. Point size indicates the total number of significant correlations. The Jaccard similarity between the sets of measurements showing significant Pearson and Spearman correlations is also indicated. The dashed line indicates equality between the sums of the absolute values of significant Pearson and Spearman correlation coefficients.

Figure S21

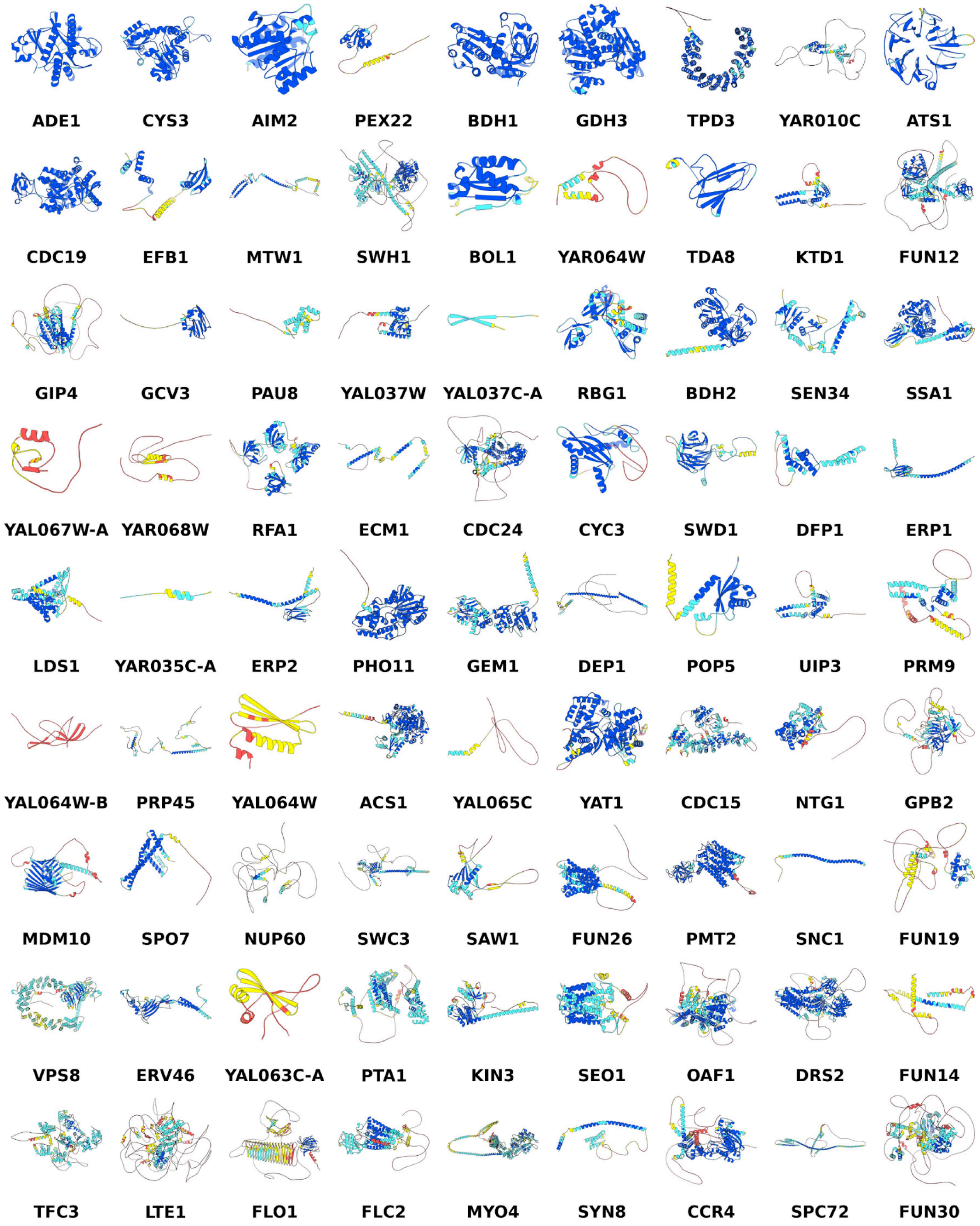


#### Figure S21. AlphaFold2-predicted structures of the target proteins.

AlphaFold2-predicted structures of all target proteins analyzed in this study. Per-residue pLDDT scores indicate prediction confidence. Protein names are indicated below each structure.

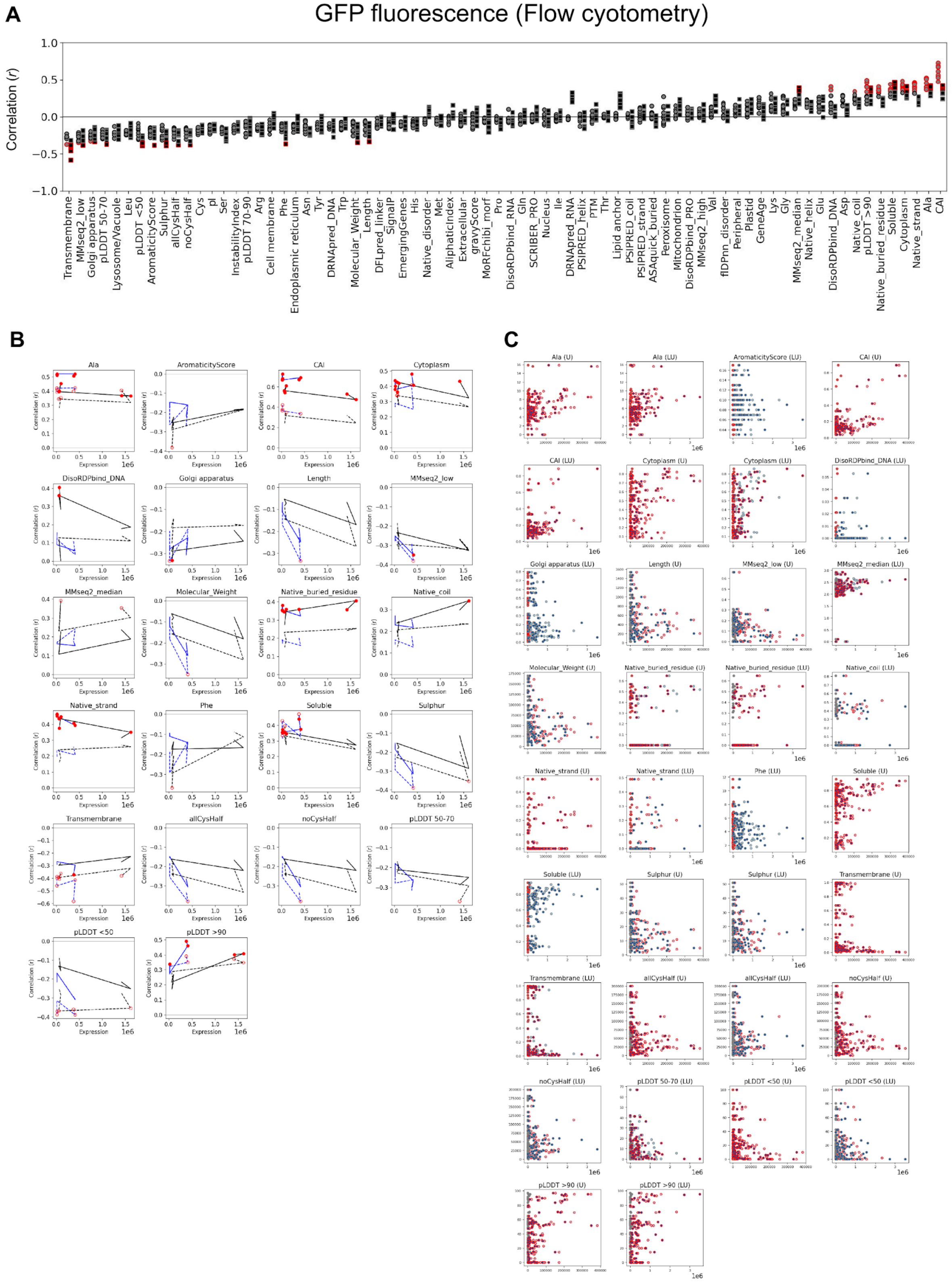


**Figure S22. Relationships between protein properties and GFP fluorescence measured by flow cytometry.**

**(A)** Correlations between each protein parameter and GFP fluorescence measured by flow cytometry under different culture and induction conditions. Parameters are ordered by correlation coefficient. Outlines indicate correlations for which either the Pearson or Spearman correlation was significant after Benjamini–Hochberg correction (FDR < 0.05).

**(B)** Changes in Pearson and Spearman correlations between selected protein parameters and GFP fluorescence across increasing expression levels. Data are shown for standard multi-copy (U) and gTOW high-copy (LU) conditions. Solid and dashed lines indicate Pearson and Spearman correlations, respectively. Filled circles indicate significant Pearson correlations, and open circles indicate significant Spearman correlations (FDR < 0.05, Benjamini–Hochberg correction). The *x*-axis indicates the corresponding MOX expression level for each experimental condition.

**(C)** Relationships between GFP fluorescence and selected protein parameters under the indicated U or LU conditions. Points represent measurements at 0 nM aTc/0 h, 500 nM aTc/6 h, and 500 nM aTc/24 h. Conditions are classified according to whether a significant Pearson or Spearman correlation was detected (FDR < 0.05). Only U or LU conditions showing at least one significant correlation across the analyzed time points are displayed.

Figure S23

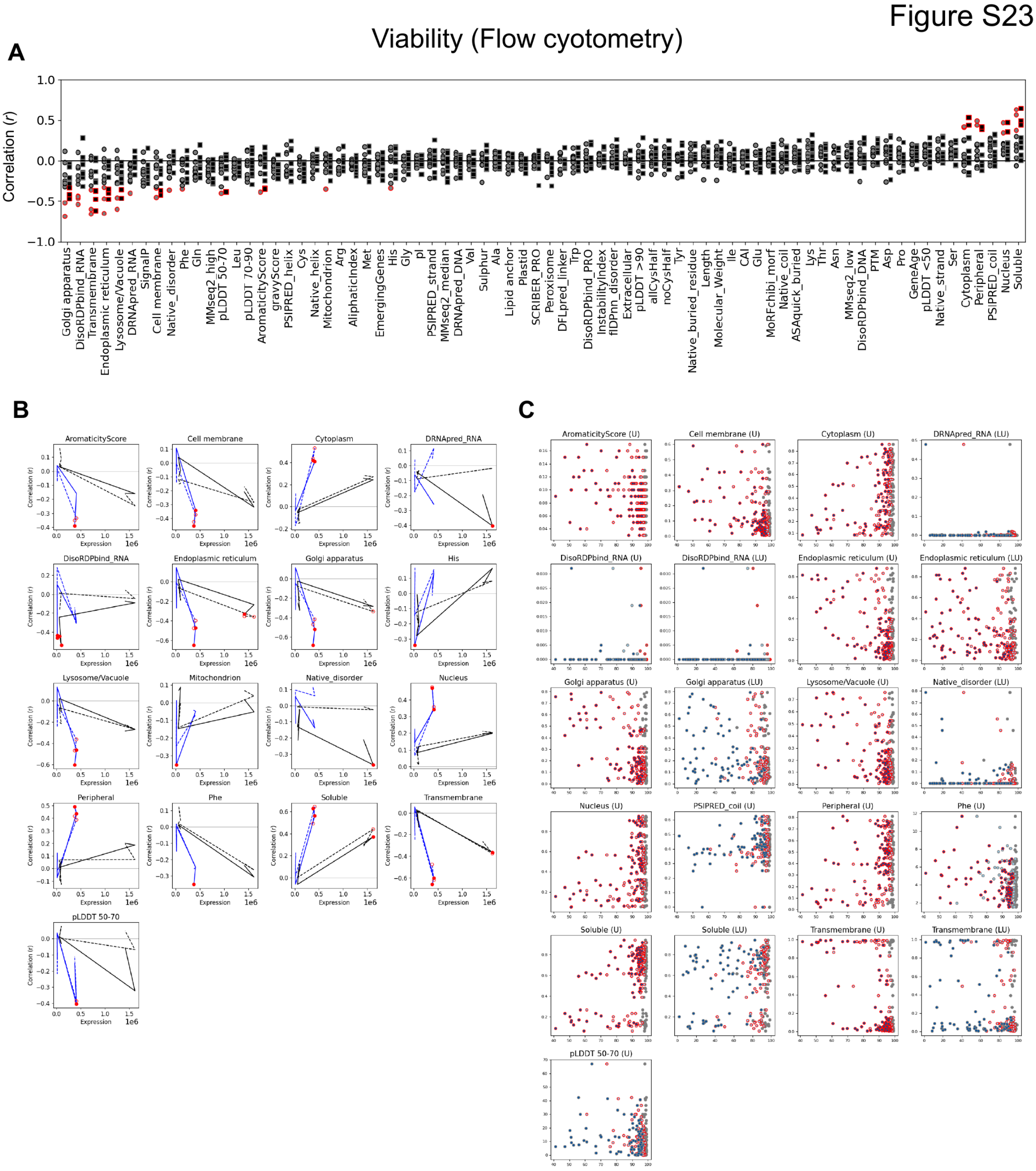


#### Figure S23. Relationships between protein properties and cell viability measured by flow cytometry.

**(A)** Correlations between each protein parameter and cell viability measured by flow cytometry under different culture and induction conditions. Each point represents the correlation coefficient obtained under an individual experimental condition. Outlines indicate correlations for which either the Pearson or Spearman correlation was significant after Benjamini–Hochberg correction (FDR < 0.05).

**(B)** Changes in Pearson and Spearman correlations between selected protein parameters and cell viability across increasing expression levels. Data are shown for standard multi-copy

(U) and gTOW high-copy (LU) conditions. Solid and dashed lines indicate Pearson and Spearman correlations, respectively. Filled circles indicate significant Pearson correlations, and open circles indicate significant Spearman correlations (FDR < 0.05, Benjamini–Hochberg correction). The *x*-axis indicates the corresponding MOX expression level for each experimental condition.

**(C)** Relationships between cell viability and selected protein parameters under the indicated U or LU conditions. Points represent measurements at 0 nM aTc/0 h, 500 nM aTc/6 h, and 500 nM aTc/24 h. Conditions are classified according to whether a significant Pearson or Spearman correlation was detected (FDR < 0.05). Only U or LU conditions showing at least one significant correlation across the analyzed time points are displayed.

Figure S24

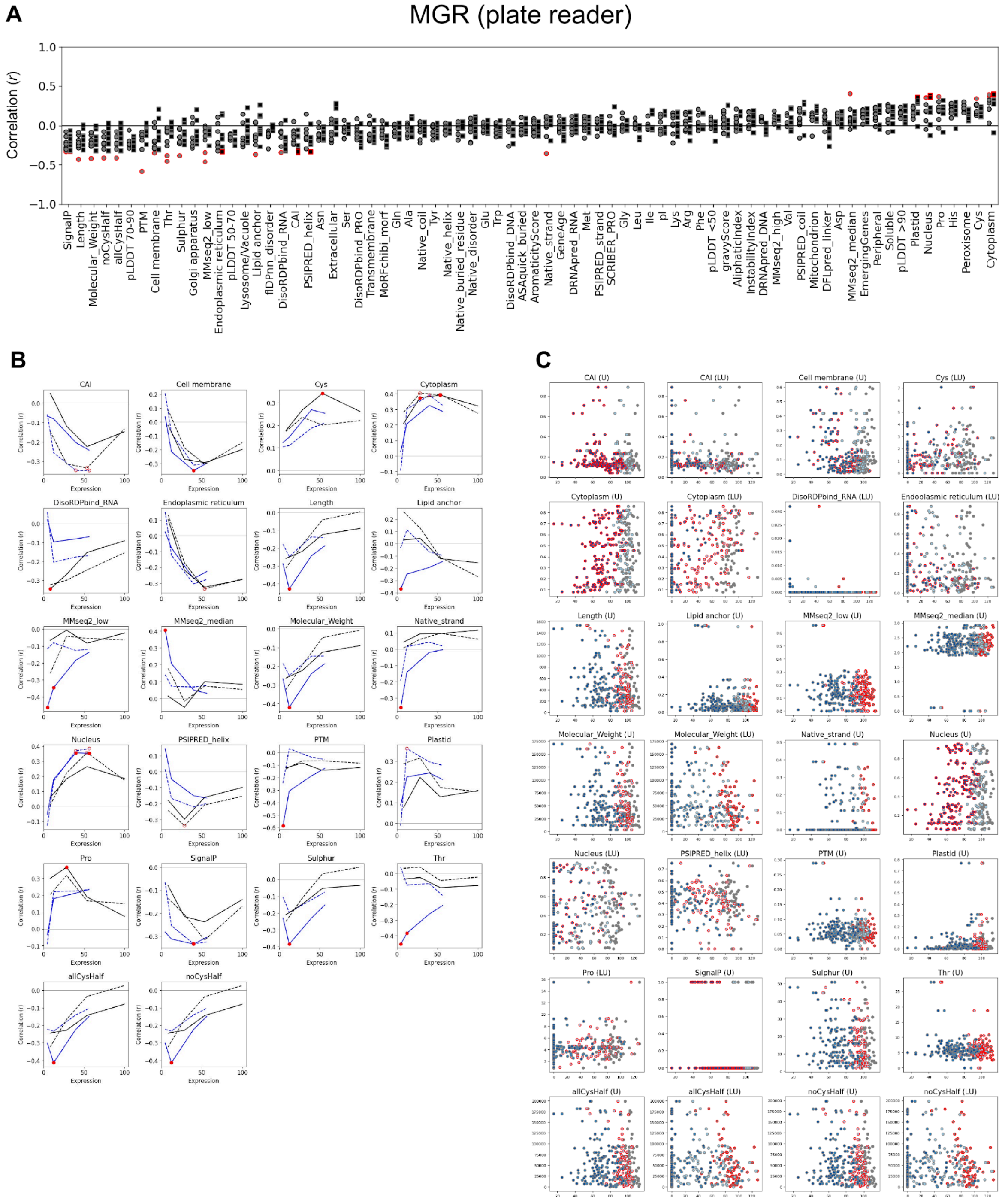


### Figure S24. Relationships between protein properties and maximum growth rate measured by plate reader.

**(A)** Correlations between each protein parameter and maximum growth rate (MGR) under different induction conditions. Each point represents the correlation coefficient obtained under an individual experimental condition. Outlines indicate correlations for which either the Pearson or Spearman correlation was significant after Benjamini–Hochberg correction (FDR < 0.05).

**(B)** Changes in Pearson and Spearman correlations between selected protein parameters and MGR across increasing expression levels. Data are shown for standard multi-copy (U) and gTOW high-copy (LU) conditions. Solid and dashed lines indicate Pearson and Spearman correlations, respectively. Filled circles indicate significant Pearson correlations, and open circles indicate significant Spearman correlations (FDR < 0.05, Benjamini–Hochberg correction). The *x*-axis indicates the corresponding MOX expression level for each induction condition.

**(C)** Relationships between MGR and selected protein parameters under the indicated U or LU conditions. Points represent measurements at 0, 50, 150, and 500 nM aTc. Conditions are classified according to whether a significant Pearson or Spearman correlation was detected (FDR < 0.05). Only U or LU conditions showing at least one significant correlation across the analyzed induction conditions are displayed.

Figure S25

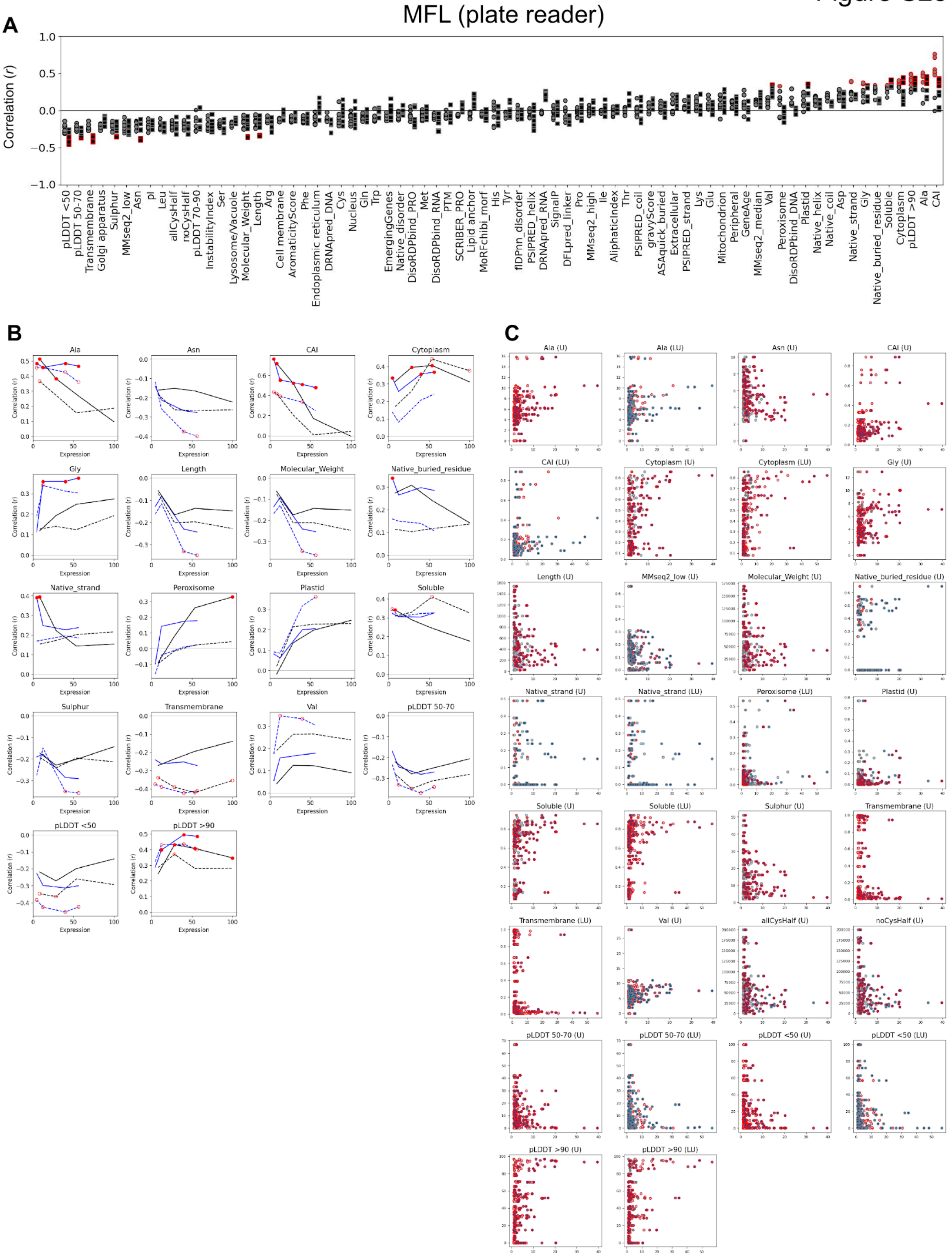
A
MFL (plate reader)
Correlation (r)
1.0
0.5
0.0
−0.5
−1.0
pLDDT <50
pLDDT 50-70
Transmembrane
Golgi apparatus
Sulphur
MMseq2_low
Asn
pI
Leu
allCysHalf
noCysHalf
pLDDT 70-90
InstabilityIndex
Ser
Lysosome/Vacuole
Molecular_Weight
Length
Arg
Cell membrane
AromaticityScore
Phe
Endoplasmic reticulum
DRNApred_DNA
Cys
Nucleus
Gln
Trp
EmergingGenes
Native_disorder
DisoRDPbind_PRO
Met
DisoRDPbind_RNA
PTM
SCRIBER_PRO
Lipid anchor
MoRFchibi_morf
His
Tyr
flDPnn_disorder
PSIPRED_helix
DRNApred_RNA
SignalP
DFLpred_linker
Pro
MMseq2_high
Ile
AliphaticIndex
Thr
PSIPRED_coil
gravyScore
ASAquick_buried
Extracellular
PSIPRED_strand
Lys
Glu
Mitochondrion
Peripheral
GeneAge
MMseq2_median
Val
Peroxisome
DisoRDPbind_DNA
Plastid
Native_helix
Native_coil
Asp
Native_strand
Gly
Native_buried_residue
Soluble
Cytoplasm
pLDDT >90
Ala
CAI
B
Ala
Asn
CAI
Cytoplasm
Gly
Length
Molecular_Weight
Native_buried_residue
Native_strand
Peroxisome
Plastid
Soluble
Sulphur
Transmembrane
Val
pLDDT 50-70
pLDDT <50
pLDDT >90
Correlation (r)
Expression
C
Ala (U)
Ala (LU)
Asn (U)
CAI (U)
CAI (LU)
Cytoplasm (U)
Cytoplasm (LU)
Gly (U)
Length (U)
MMseq2_low (U)
Molecular_Weight (U)
Native_buried_residue (U)
Native_strand (U)
Native_strand (LU)
Peroxisome (LU)
Plastid (U)
Soluble (U)
Soluble (LU)
Sulphur (U)
Transmembrane (U)
Transmembrane (LU)
Val (U)
allCysHalf (U)
noCysHalf (U)
pLDDT 50-70 (U)
pLDDT 50-70 (LU)
pLDDT <50 (U)
pLDDT <50 (LU)
pLDDT >90 (U)
pLDDT >90 (LU)

#### Figure S25. Relationships between protein properties and maximum fluorescence level measured by plate reader.

**(A)** Correlations between each protein parameter and maximum fluorescence level (MFL) under different induction conditions. Each point represents the correlation coefficient obtained under an individual experimental condition. Outlines indicate correlations for which either the Pearson or Spearman correlation was significant after Benjamini–Hochberg correction (FDR < 0.05).

**(B)** Changes in Pearson and Spearman correlations between selected protein parameters and MFL across increasing expression levels. Data are shown for standard multi-copy (U) and gTOW high-copy (LU) conditions. Solid and dashed lines indicate Pearson and Spearman correlations, respectively. Filled and open circles indicate significant Pearson and Spearman correlations, respectively (FDR < 0.05, Benjamini–Hochberg correction). The *x*-axis indicates the corresponding MOX expression level for each induction condition.

**(C)** Relationships between MFL and selected protein parameters under the indicated U or LU conditions. Points represent measurements at 0, 50, 150, and 500 nM aTc. Conditions are classified according to whether a significant Pearson or Spearman correlation was detected (FDR < 0.05). Only U or LU conditions showing at least one significant correlation across the analyzed induction conditions are displayed.

Figure S26

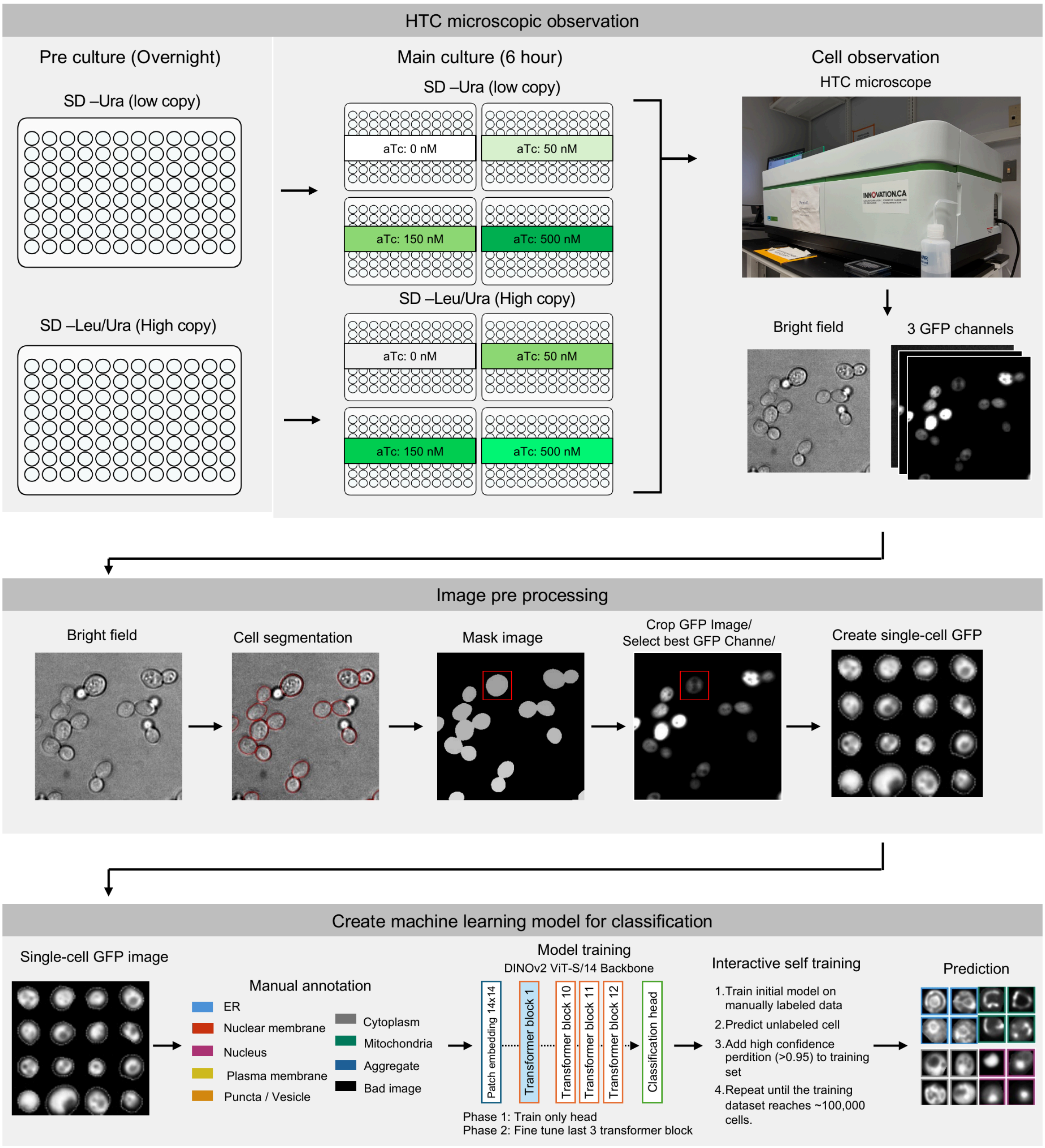


#### Figure S26. Workflow for high-content imaging and machine-learning-based classification of subcellular localization.

Yeast cells expressing target proteins fused to moxGFP were precultured overnight under standard multi-copy (SC –Ura) or gTOW high-copy (SC –Leu/Ura) conditions with 0, 50, 150, or 500 nM aTc, then transferred to fresh medium with the same medium composition and aTc concentration and cultured for 6 h. Bright-field images and GFP fluorescence images were acquired using a high-content microscope. Individual cells were segmented from bright-field images, and corresponding GFP images were cropped to generate single-cell GFP images. Subcellular localization was classified using a machine-learning model based on the DINOv2 ViT-S/14 backbone. Training images were initially manually annotated into Cytoplasm, Aggregate, Nucleus, ER, Mitochondria, Nuclear membrane, Plasma membrane, Puncta/Vesicle, or Bad image classes. The training dataset was subsequently expanded by iterative self-training, in which high-confidence predictions (>0.95) were added to the training set, until approximately 100,000 labeled images were obtained. The resulting model was used to classify GFP localization patterns in the complete single-cell image dataset.

Figure S27

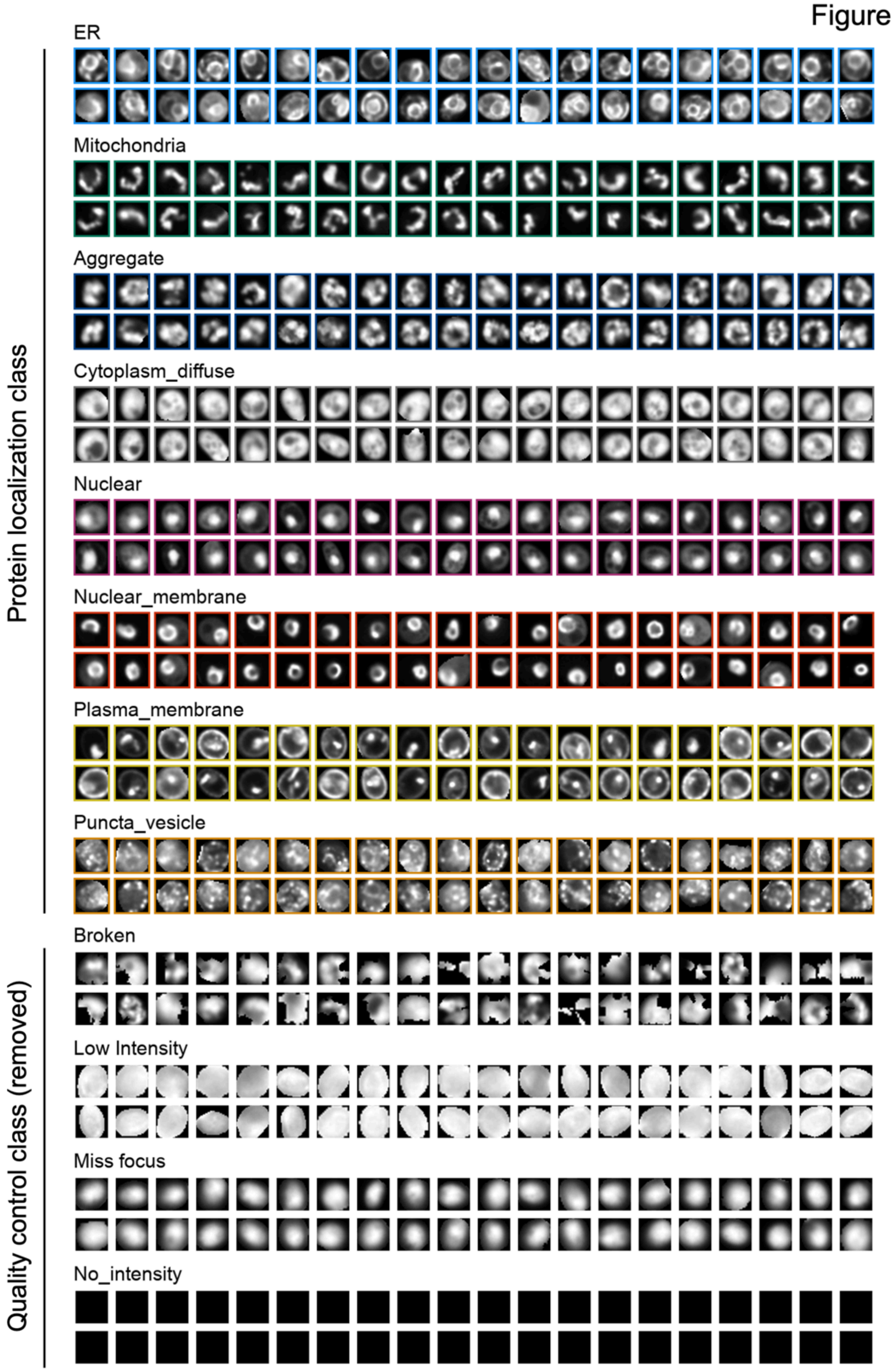


**Figure S27. Representative single-cell images for localization classes used in the machine-learning classification.**

Representative GFP images assigned to each class by the machine-learning classifier. Eight protein localization classes used for subsequent analyses are shown in the upper section: ER, Mitochondria, Aggregate, Cytoplasm_diffuse, Nuclear, Nuclear_membrane, Plasma_membrane, and Puncta_vesicle. Four quality-control classes excluded from subsequent localization analyses are shown in the lower section: Broken, Low Intensity, Miss focus, and No_intensity. Localization classes correspond to those used in **Figure 4**.

Figure S28

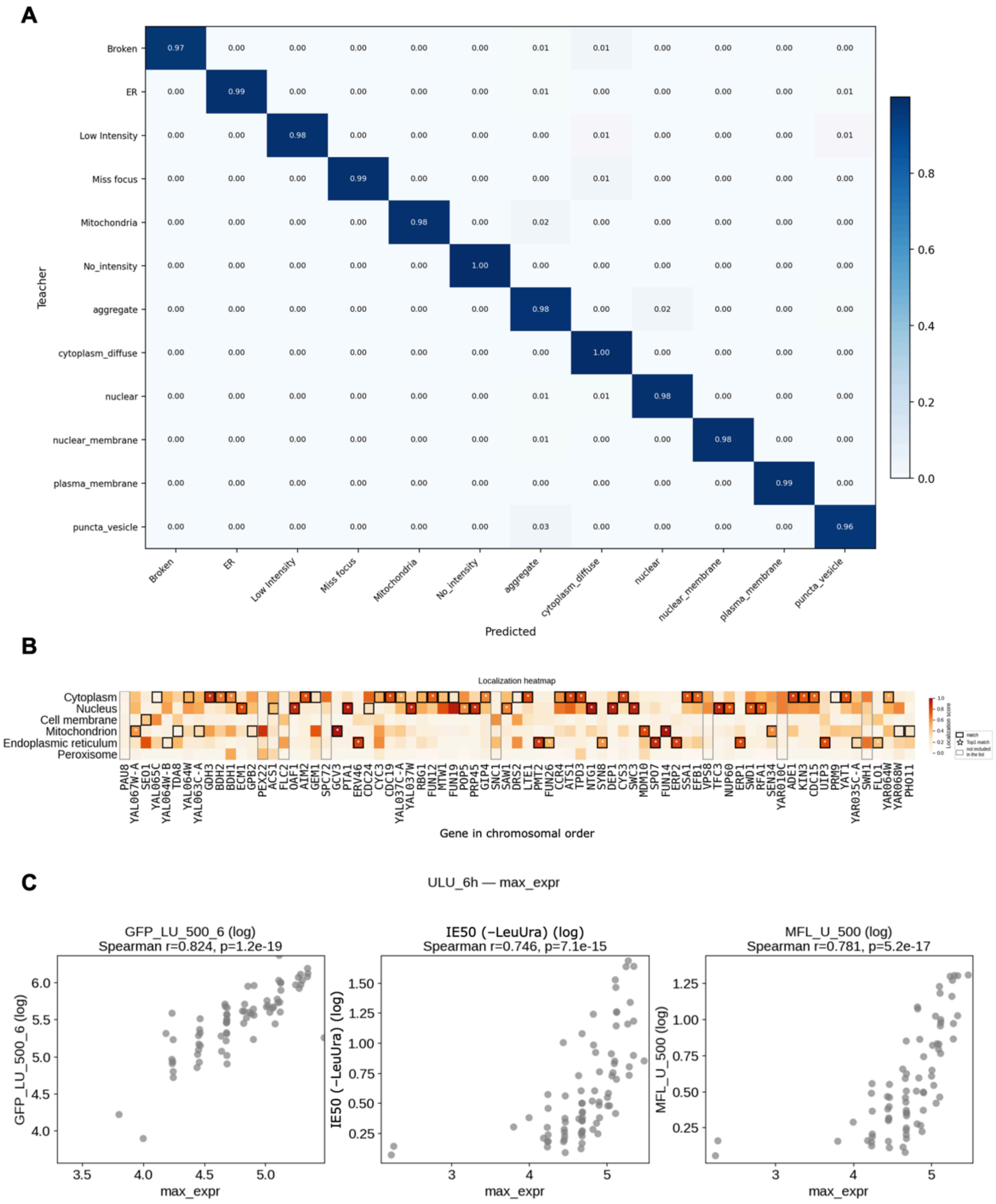


**Figure S28. Validation of localization classification and fluorescence measurements obtained by high-content imaging.**

**(A)** Confusion matrix of the machine-learning classifier used for single-cell localization classification. Rows indicate manually assigned classes and columns indicate predicted classes. Values indicate the fraction of cells assigned to each predicted class.

**(B)** Comparison of low-expression localization, defined as the most frequently predicted localization class among up to 200 cells with the lowest GFP fluorescence intensities for each target protein, with subcellular localization scores predicted by DeepLoc 2.0. Target genes are arranged from left to right in chromosomal order. The heatmap shows predicted localization scores, with localization observed by high-content imaging indicated for comparison. Asterisks indicate cases in which the observed localization corresponded to the highest-scoring predicted localization. Proteins classified as Aggregate or Puncta/Vesicle are indicated separately because these classes were not included among the DeepLoc 2.0 localization categories shown.

**(C)** Relationship between the maximum GFP fluorescence intensity measured by high-content imaging (`max_expr`) and GFP fluorescence measured by flow cytometry at 6 h after induction with 500 nM aTc (left), IE50 under –LeuUra conditions (middle), and MFL measured by plate reader at 500 nM aTc (right). Spearman correlation coefficients and *p*-values are shown above each panel.

Figure S29

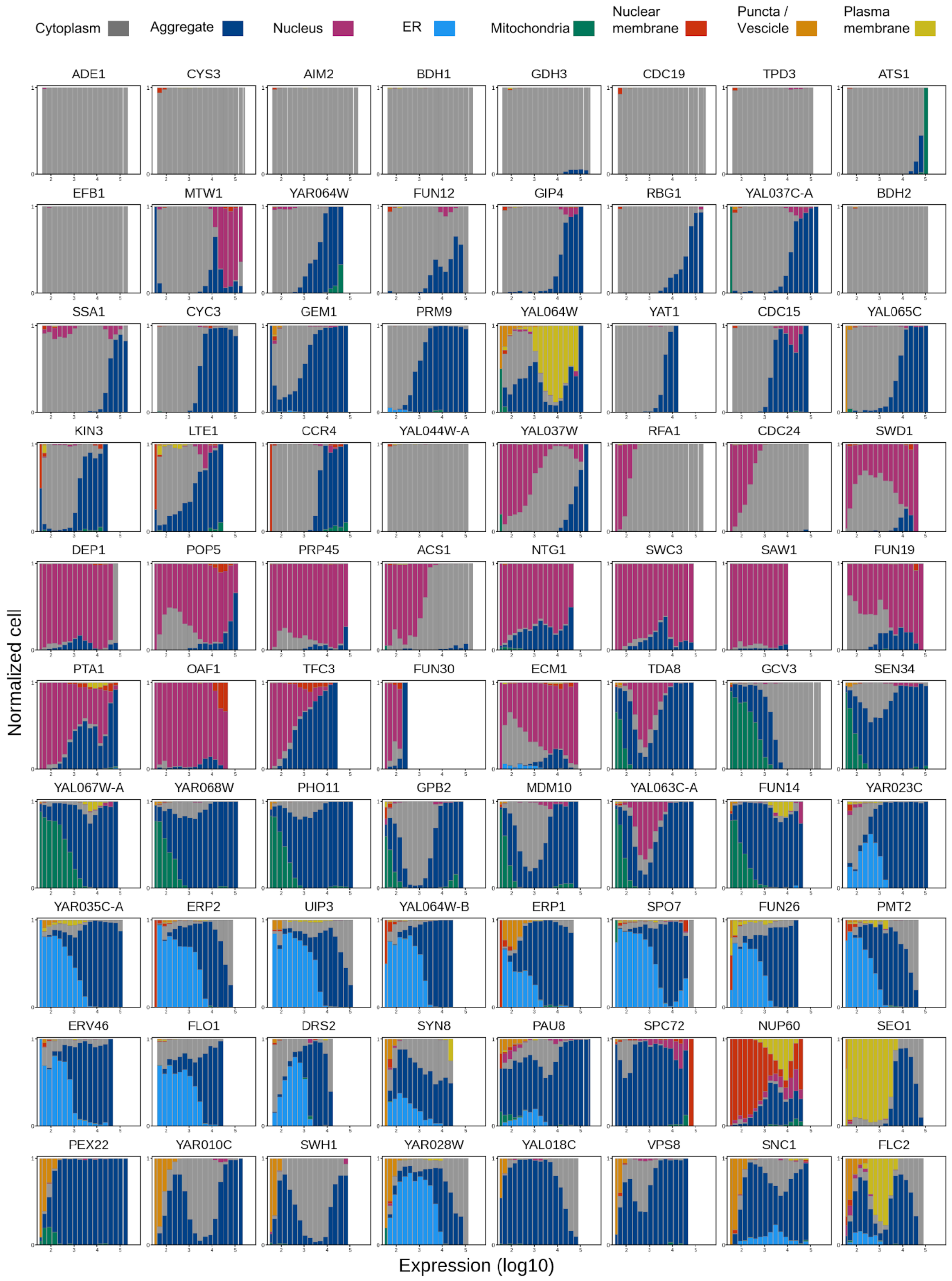

Cytoplasm
Aggregate
Nucleus
ER
Mitochondria
Nuclear membrane
Puncta / Vescicle
Plasma membrane
ADE1
CYS3
AIM2
BDH1
GDH3
CDC19
TPD3
ATS1
EFB1
MTW1
YAR064W
FUN12
GIP4
RBG1
YAL037C-A
BDH2
SSA1
CYC3
GEM1
PRM9
YAL064W
YAT1
CDC15
YAL065C
KIN3
LTE1
CCR4
YAL044W-A
YAL037W
RFA1
CDC24
SWD1
DEP1
POP5
PRP45
ACS1
NTG1
SWC3
SAW1
FUN19
PTA1
OAF1
TFC3
FUN30
ECM1
TDA8
GCV3
SEN34
YAL067W-A
YAR068W
PHO11
GPB2
MDM10
YAL063C-A
FUN14
YAR023C
YAR035C-A
ERP2
UIP3
YAL064W-B
ERP1
SPO7
FUN26
PMT2
ERV46
FLO1
DRS2
SYN8
PAU8
SPC72
NUP60
SEO1
PEX22
YAR010C
SWH1
YAR028W
YAL018C
VPS8
SNC1
FLC2
Normalized cell
Expression (log10)

#### Figure S29. Expression-dependent changes in subcellular localization for individual target proteins.

Distribution of subcellular localization classes as a function of GFP fluorescence intensity for each target protein. Single cells were binned according to GFP fluorescence intensity, and the fraction of cells assigned to each localization class was calculated within each bin. The localization classes are Cytoplasm, Aggregate, Nucleus, ER, Mitochondria, Nuclear membrane, Puncta/Vesicle, and Plasma membrane, as indicated at the top. The *x*-axis indicates GFP fluorescence intensity ($\log_{10}$), and the *y*-axis indicates the normalized fraction of cells in each localization class.

Figure S30

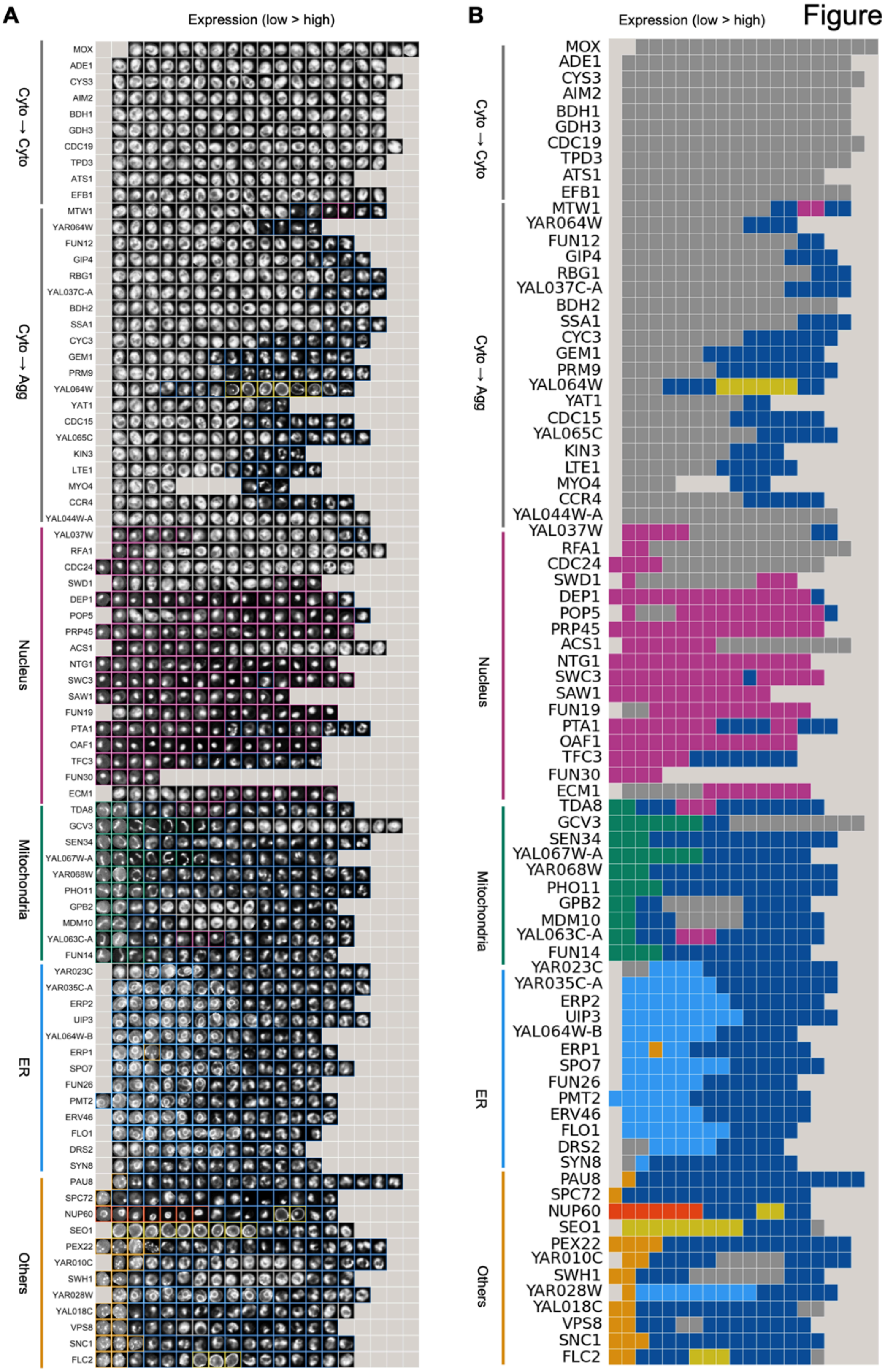
A
B
Expression (low > high)
Expression (low > high)
Cyto → Cyto
Cyto → Agg
Nucleus
Mitochondria
ER
Others
MOX
ADE1
CYS3
AIM2
BDH1
GDH3
CDC19
TPD3
ATS1
EFB1
MTW1
YAR064W
FUN12
GIP4
RBG1
YAL037C-A
BDH2
SSA1
CYC3
GEM1
PRM9
YAL064W
YAT1
CDC15
YAL065C
KIN3
LTE1
MYO4
CCR4
YAL044W-A
YAL037W
RFA1
CDC24
SWD1
DEP1
POP5
PRP45
ACS1
NTG1
SWC3
SAW1
FUN19
PTA1
OAF1
TFC3
FUN30
ECM1
TDA8
GCV3
SEN34
YAL067W-A
YAR068W
PHO11
GPB2
MDM10
YAL063C-A
FUN14
YAR023C
YAR035C-A
ERP2
UIP3
YAL064W-B
ERP1
SPO7
FUN26
PMT2
ERV46
FLO1
DRS2
SYN8
PAU8
SPC72
NUP60
SEO1
PEX22
YAR010C
SWH1
YAR028W
YAL018C
VPS8
SNC1
FLC2

#### Figure S30. Expression-dependent changes in subcellular localization across all target proteins.

Representative single-cell GFP images and corresponding dominant localization classes across increasing protein expression levels for all target proteins analyzed by high-content imaging. Target proteins are grouped according to their localization patterns at low expression and subsequent localization changes upon overexpression, as indicated on the left. Within each row, GFP fluorescence intensity increases from left to right. The localization class assigned by the machine-learning classifier is indicated for each single-cell image. The heatmap on the right shows the dominant localization class in each expression bin for the corresponding target proteins. Localization classes are defined in **Figures 4B and S27**. Expression bins without valid localization data are indicated separately.

Figure S31

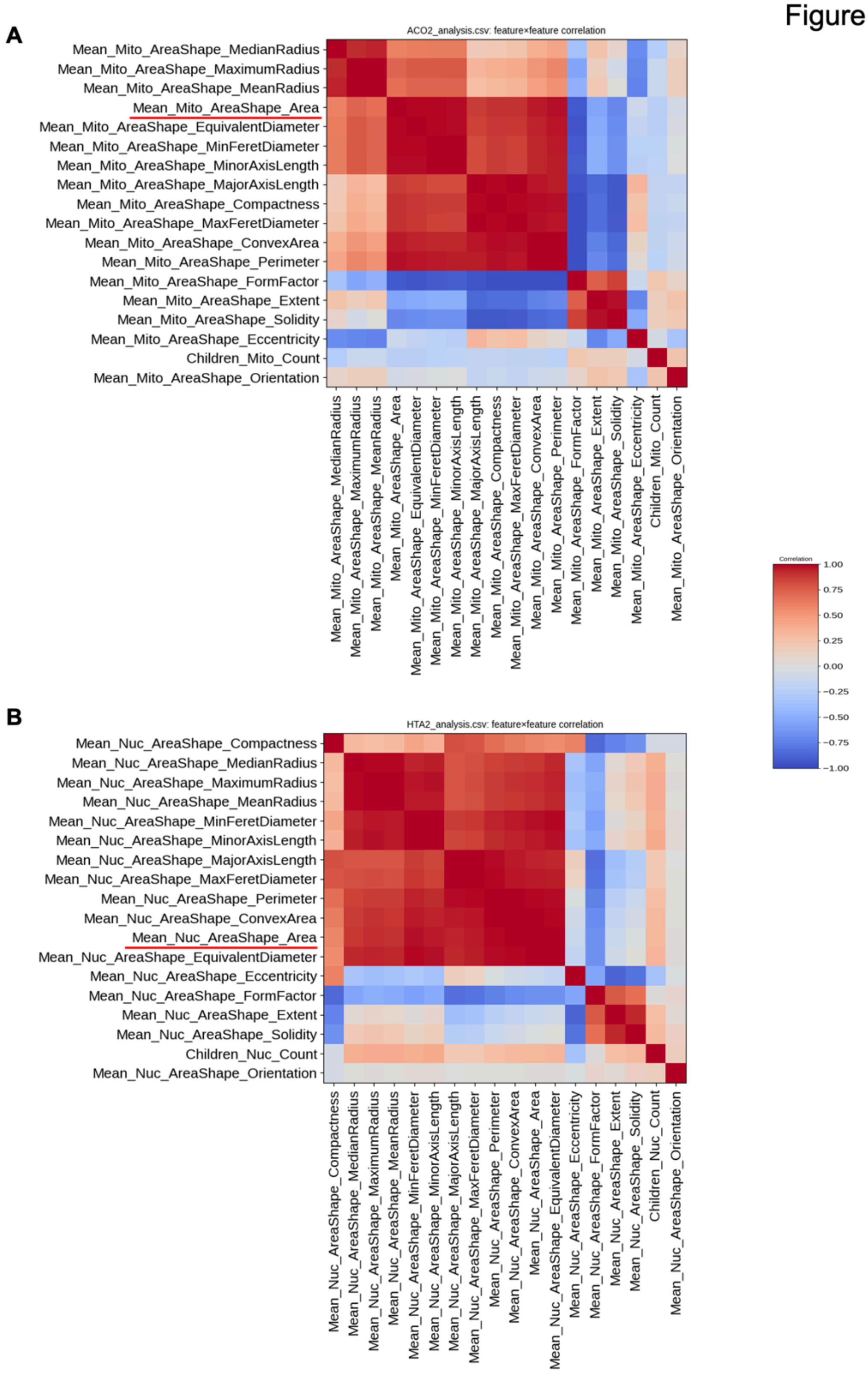


**Figure S31. Correlations among morphological features used for quantification of mitochondrial and nuclear morphology.**

Correlation matrices of morphological features extracted from Aco2-RFP-labeled mitochondria (top) and Hta2-RFP-labeled nuclei (bottom) by high-content image analysis. Matrix entries represent pairwise correlation coefficients ranging from −1 to 1. `Mean_Mito_AreaShape_Area` and `Mean_Nuc_AreaShape_Area`, used to quantify mitochondrial and nuclear size in **Figure 4G**, are underlined.

Figure S32

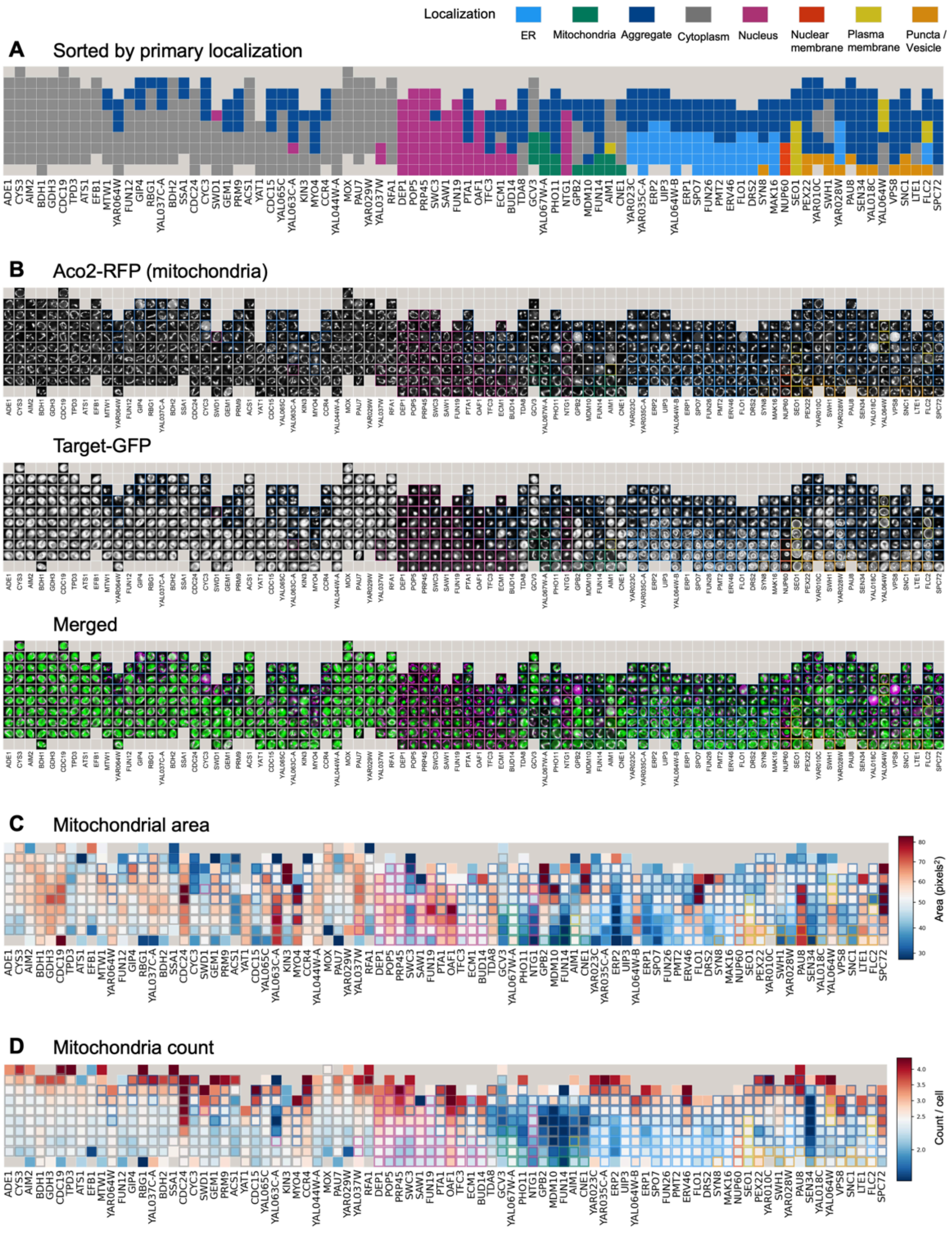


**Figure S32. Effects of protein overexpression on mitochondrial morphology.**

(**A**) Summary of the subcellular localization patterns of the overexpressed proteins. Proteins are ordered according to their primary localization, as in **Figure 4**. Boxes indicate the localization patterns observed at different expression levels.

(**B**) Representative fluorescence microscopy images of cells expressing the indicated GFP-tagged proteins together with Aco2-RFP as a mitochondrial marker. Aco2-RFP, Target-GFP, and merged images are shown. Proteins are arranged in the same order as in (**A**).

(**C**, **D**) Single-cell quantification of mitochondrial morphology using combined data from cells cultured for 6 h under standard multicopy (–Ura) and gTOW high-copy (–LeuUra) conditions. Heat maps show the mean mitochondrial area per cell (**C**) and the number of mitochondria per cell (**D**). Area values in (C) are shown in pixels. Each square shows the mean value of the indicated morphological parameter across cells within a GFP fluorescence intensity bin for the corresponding target protein. The mitochondrial area measurements shown in (C) were used for the analysis in **Figure 4G**.

Figure S33

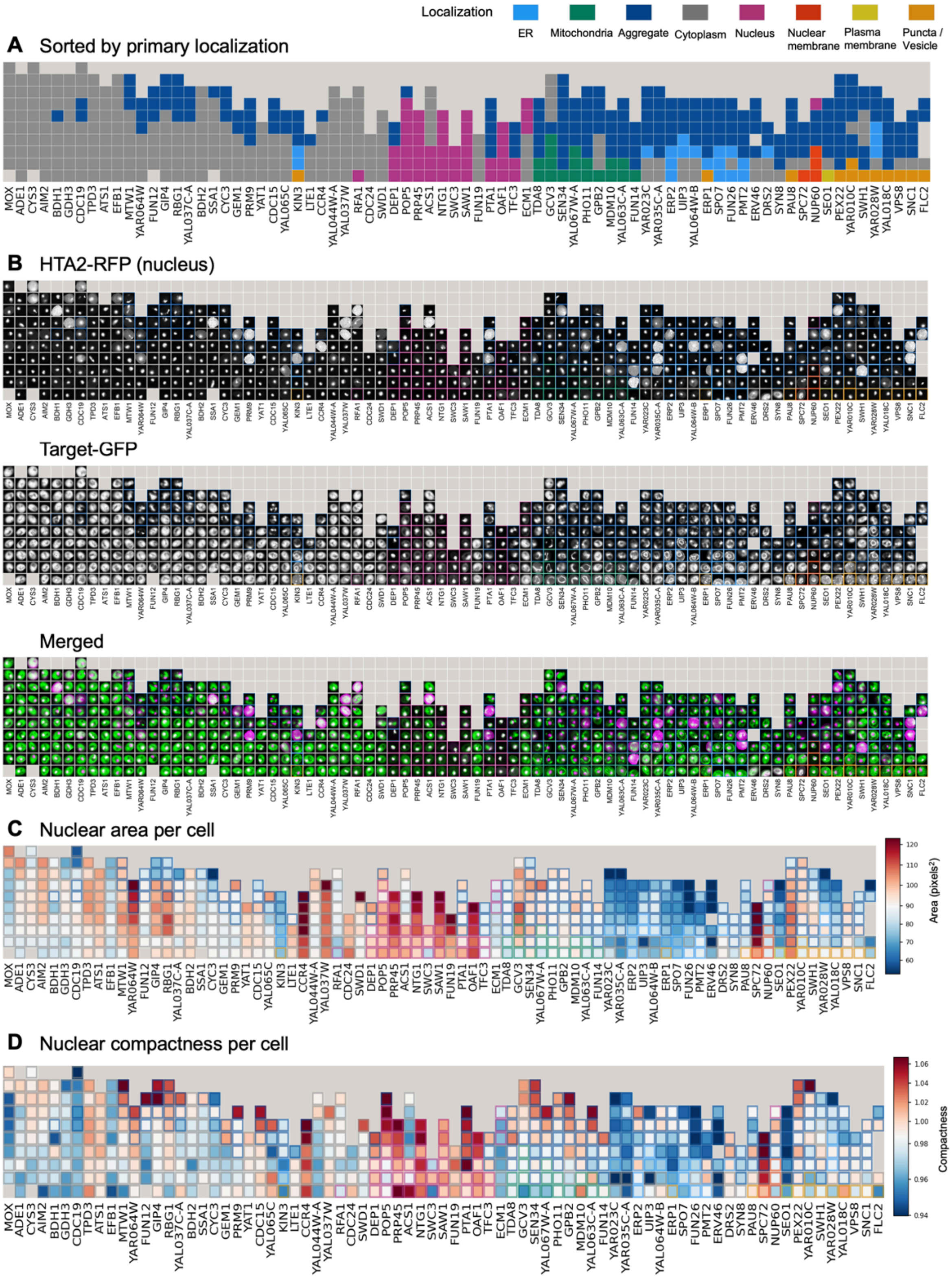
Localization
ER
Mitochondria
Aggregate
Cytoplasm
Nucleus
Nuclear membrane
Plasma membrane
Puncta / Vesicle
A Sorted by primary localization
B HTA2-RFP (nucleus)
Target-GFP
Merged
C Nuclear area per cell
Area (pixels²)
D Nuclear compactness per cell
Compactness
MOX
ADE1
CYS3
AIM2
BDH1
GDH3
CDC19
TPD3
ATS1
EFB1
MTW1
YAR064W
FUN12
GIP4
RBG1
YAL037C-A
BDH2
SSA1
CYC3
GEM1
PRM9
YAT1
CDC15
YAL065C
KIN3
LTE1
CCR4
YAL044W-A
YAL037W
RFA1
CDC24
SWD1
DEP1
POP5
PRP45
ACS1
NTG1
SWC3
SAW1
FUN19
PTA1
OAF1
TFC3
ECM1
TDA8
GCV3
SEN34
YAL067W-A
PHO11
GPB2
MDM10
YAL063C-A
FUN14
YAR023C
YAR035C-A
ERP2
UIP3
YAL064W-B
ERP1
SPO7
FUN26
PMT2
ERV46
DRS2
SYN8
PAU8
SPC72
NUP60
SEO1
PEX22
YAR010C
SWH1
YAR028W
YAL018C
VPS8
SNC1
FLC2

#### Figure S33. Effects of protein overexpression on nuclear morphology.

**(A)** Summary of the subcellular localization patterns of the overexpressed proteins. Proteins are ordered according to their primary localization, as in **Figure 4**. Boxes indicate the localization patterns observed at different expression levels.

**(B)** Representative fluorescence microscopy images of cells expressing the indicated GFP-tagged proteins together with Hta2-RFP as a nuclear marker. Hta2-RFP, Target-GFP, and merged images are shown. Proteins are arranged in the same order as in **(A)**.

**(C, D)** Single-cell quantification of nuclear morphology using combined data from cells cultured for 6 h under standard multicopy (–Ura) and gTOW high-copy (–LeuUra) conditions. Heat maps show the nuclear area per cell **(C)** and nuclear compactness per cell **(D)**. Area values in (C) are shown in pixels$^2$. Each square shows the mean value of the indicated morphological parameter across cells within a GFP fluorescence intensity bin for the corresponding target protein. The nuclear area measurements shown in (C) were used for the analysis in **Figure 4G**.

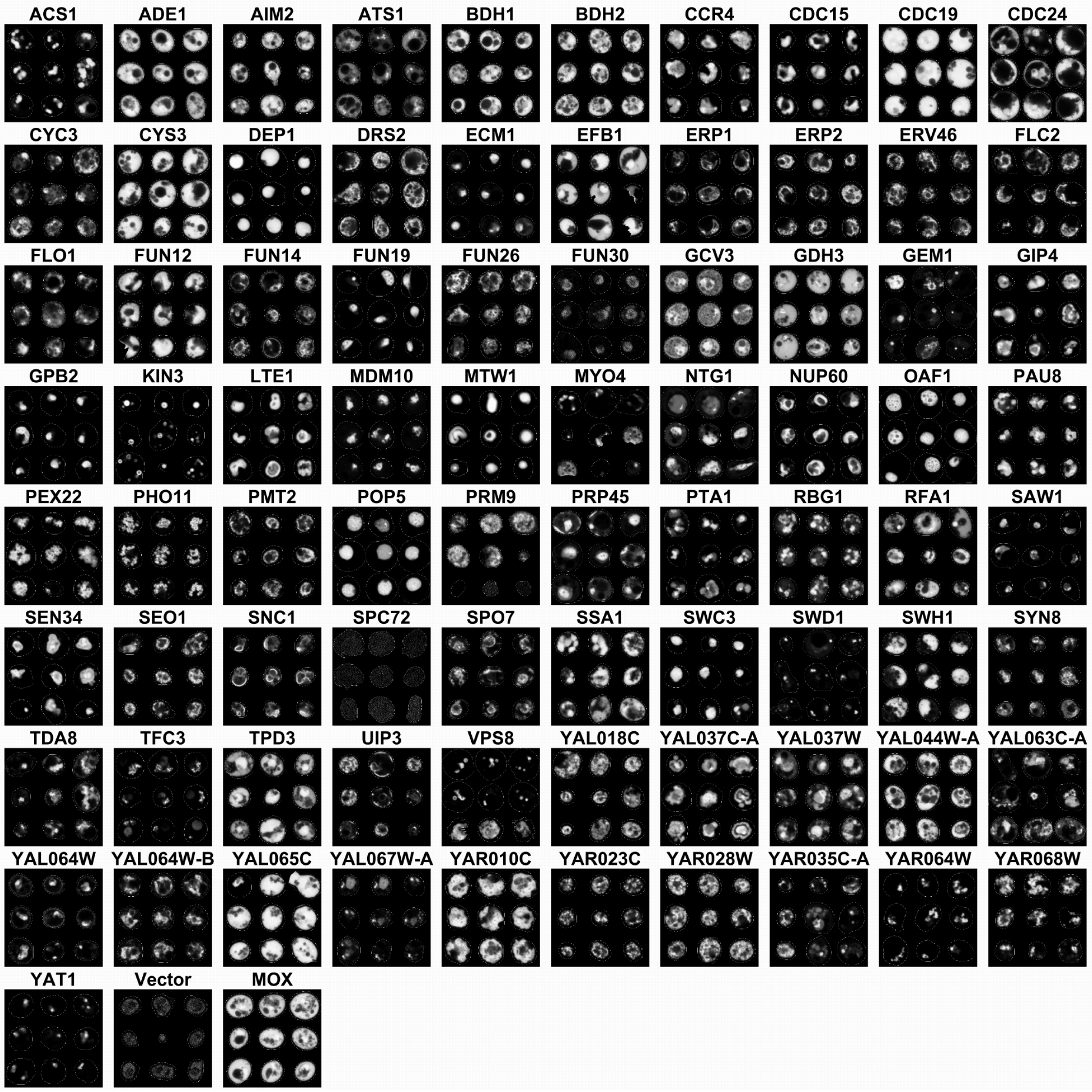


#### Figure S34. Subcellular localization patterns of target proteins under strong overexpression.

Representative GFP fluorescence images of cells strongly overexpressing the indicated target proteins. Cells expressing GFP-fused target proteins were cultured under gTOW high-copy conditions in the presence of 500 nM aTc and observed by fluorescence microscopy. Multiple cells are shown for each target protein to illustrate the characteristic localization patterns and cell-to-cell variation observed under strong overexpression. Vector and MOX controls are shown at the bottom. Representative single-cell images selected from these data are shown in **Figure 5A**.

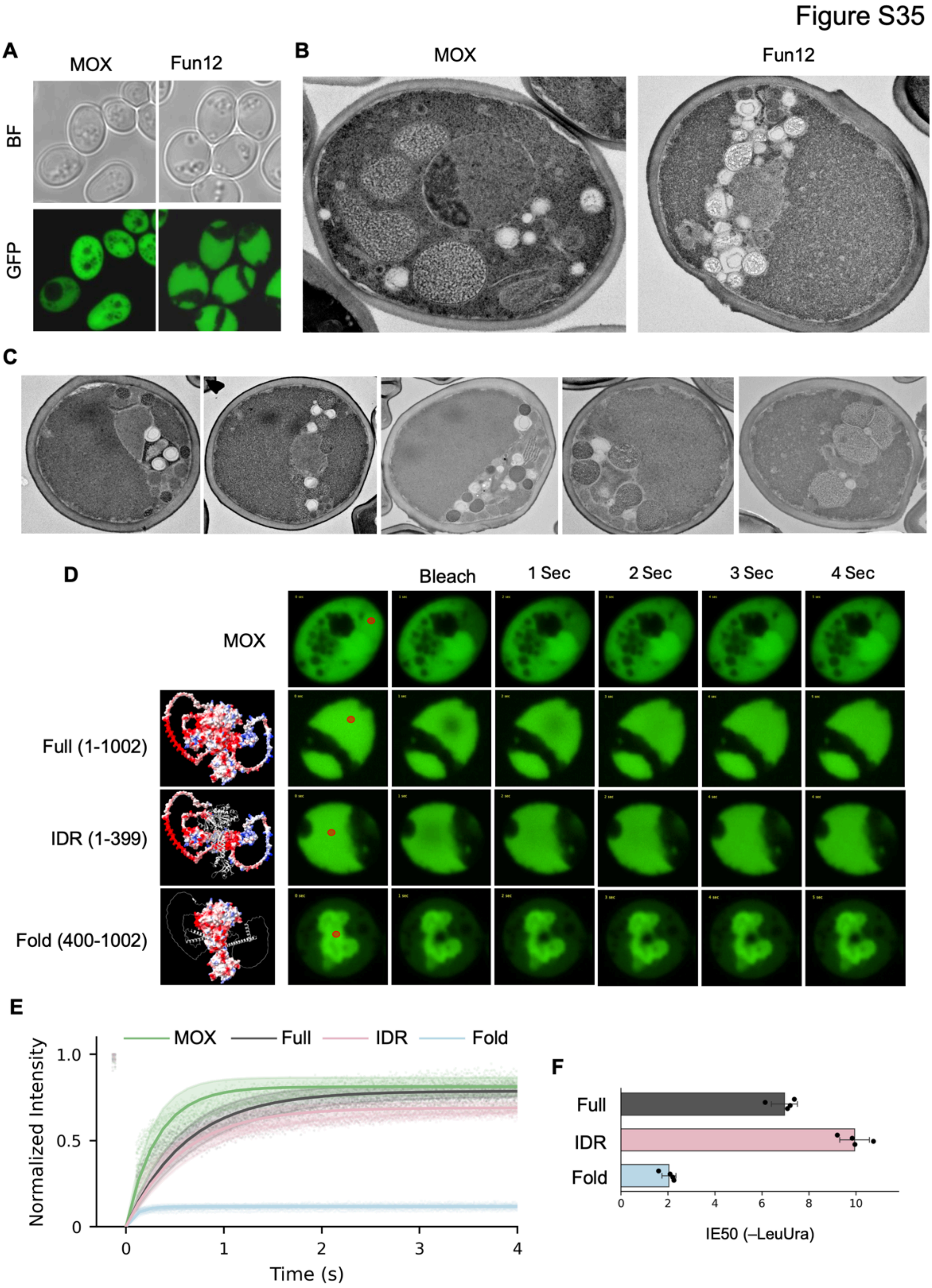


**Figure S35. The N-terminal IDR of Fun12 drives formation of a dynamic cytoplasmic state under strong overexpression.**

**(A)** Representative bright-field (BF) and GFP fluorescence images of cells expressing MOX or full-length Fun12-GFP under strong overexpression conditions. Fun12 overexpression produces a characteristic intracellular organization in which GFP fluorescence occupies a large region of the cytoplasm.

**(B)** Representative transmission electron microscopy images of cells overexpressing MOX or Fun12. For TEM analysis, cells were cultured in –LeuUra medium containing 500 nM aTc at 30°C for 12 h. Fun12-overexpressing cells show displacement and clustering of intracellular organelles, leaving a large, relatively homogeneous cytoplasmic region.

**(C)** Additional transmission electron microscopy images of Fun12-overexpressing cells showing representative variations of the characteristic intracellular morphology.

**(D)** FRAP analysis of MOX, full-length Fun12 (residues 1–1002), the N-terminal intrinsically disordered region (IDR; residues 1–399), and the C-terminal folded region (Fold; residues 400–1002). Representative fluorescence images before and after photobleaching are shown. Circles indicate the bleached regions. Predicted structures of the corresponding Fun12 constructs by AlphaFold2 are shown on the left.

**(E)** Quantification of fluorescence recovery after photobleaching for the constructs shown in **(D)**. Fluorescence intensity in the bleached region was background-corrected, normalized to the maximum corrected intensity of each measurement, and plotted as a function of time after bleaching. Individual measurements and fitted recovery curves are shown. Ten cells were analyzed for each condition.

**(F)** IE50 values of full-length Fun12, the IDR, and the folded region (Fold) under –LeuUra conditions. Bars show mean ± SD, and points represent biological replicates. IE50 values are plotted on a linear scale.

Figure S36

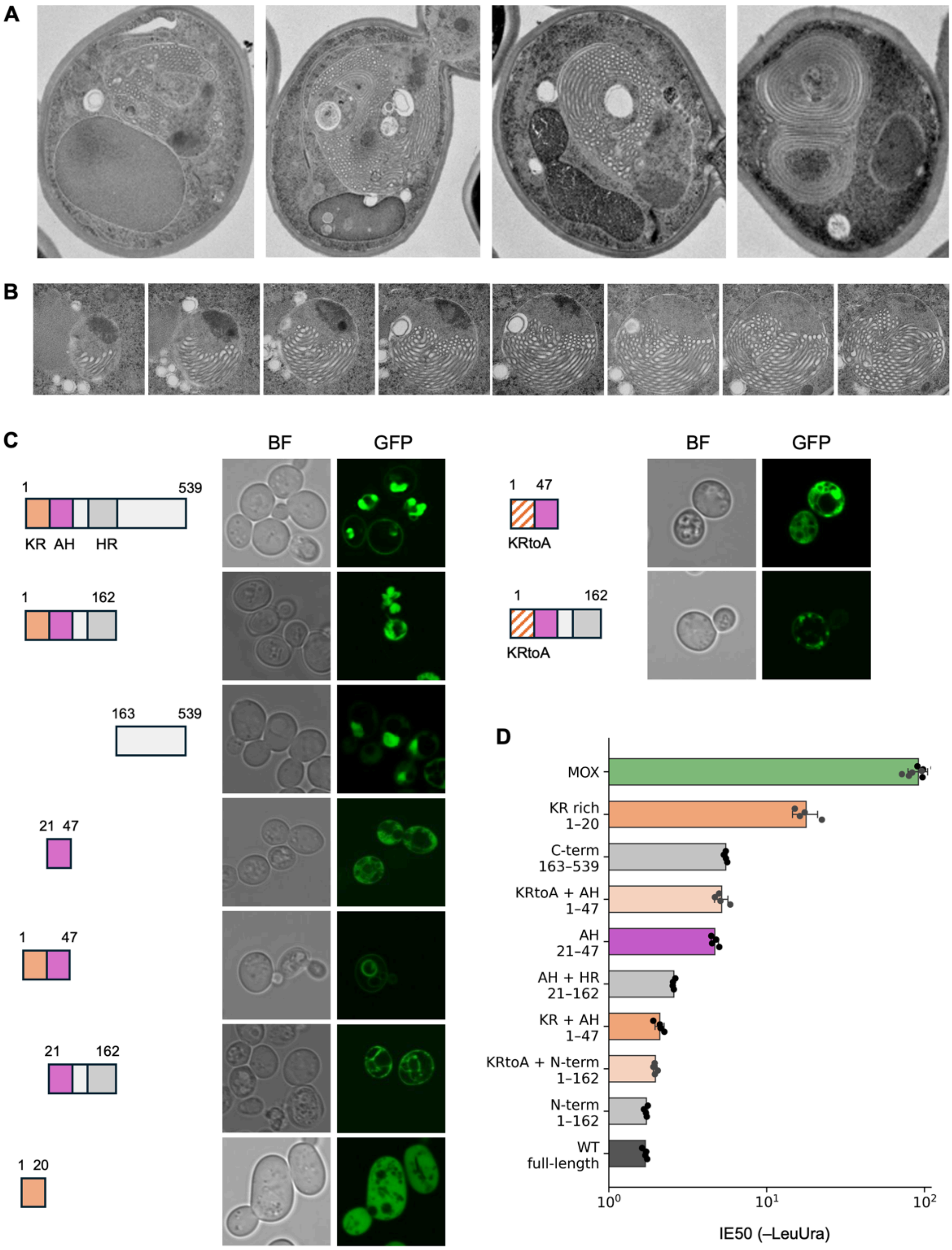
A
B
C
BF
GFP
1
539
KR
AH
HR
162
163
21
47
20
KRtoA
D
MOX
KR rich
1–20
C-term
163–539
KRtoA + AH
1–47
AH
21–47
AH + HR
21–162
KR + AH
1–47
KRtoA + N-term
1–162
N-term
1–162
WT
full-length
IE50 (–LeuUra)

#### Figure S36. Structural and sequence determinants of membrane remodeling induced by Nup60 overexpression.

**(A)** Representative transmission electron microscopy images of cells overexpressing full-length Nup60. For TEM analysis, cells were cultured in –LeuUra medium containing 500 nM aTc at 30°C for 12 h**.** Extensive membrane proliferation was observed around the nucleus, forming large multilayered and concentric membrane structures.

**(B)** Serial-section transmission electron microscopy of the membrane structure induced by Nup60 overexpression. Consecutive sections through the same structure show that the membranes form a multilayered lamellar organization rather than a single continuous tubular membrane.

**(C)** Subcellular localization and membrane-remodeling phenotypes of Nup60 fragments and mutants. Schematics indicate the Nup60 regions included in each construct, including the N-terminal KR-rich region (KR), amphipathic helix (AH), and adjacent helical region (HR). Representative bright-field (BF) and GFP fluorescence images are shown. Constructs containing both the KR-rich region and AH reproduced the characteristic nuclear-envelope-associated localization and membrane structures observed with full-length Nup60, whereas removal or mutation of the KR-rich region altered localization and membrane morphology. KRtoA indicates substitution of Lys and Arg residues in the KR-rich region with Ala.

**(D)** IE50 values of full-length Nup60 and the indicated fragments and mutants under –LeuUra conditions. Bars show mean ± SD, and points represent biological replicates. IE50 values are plotted on a logarithmic scale.

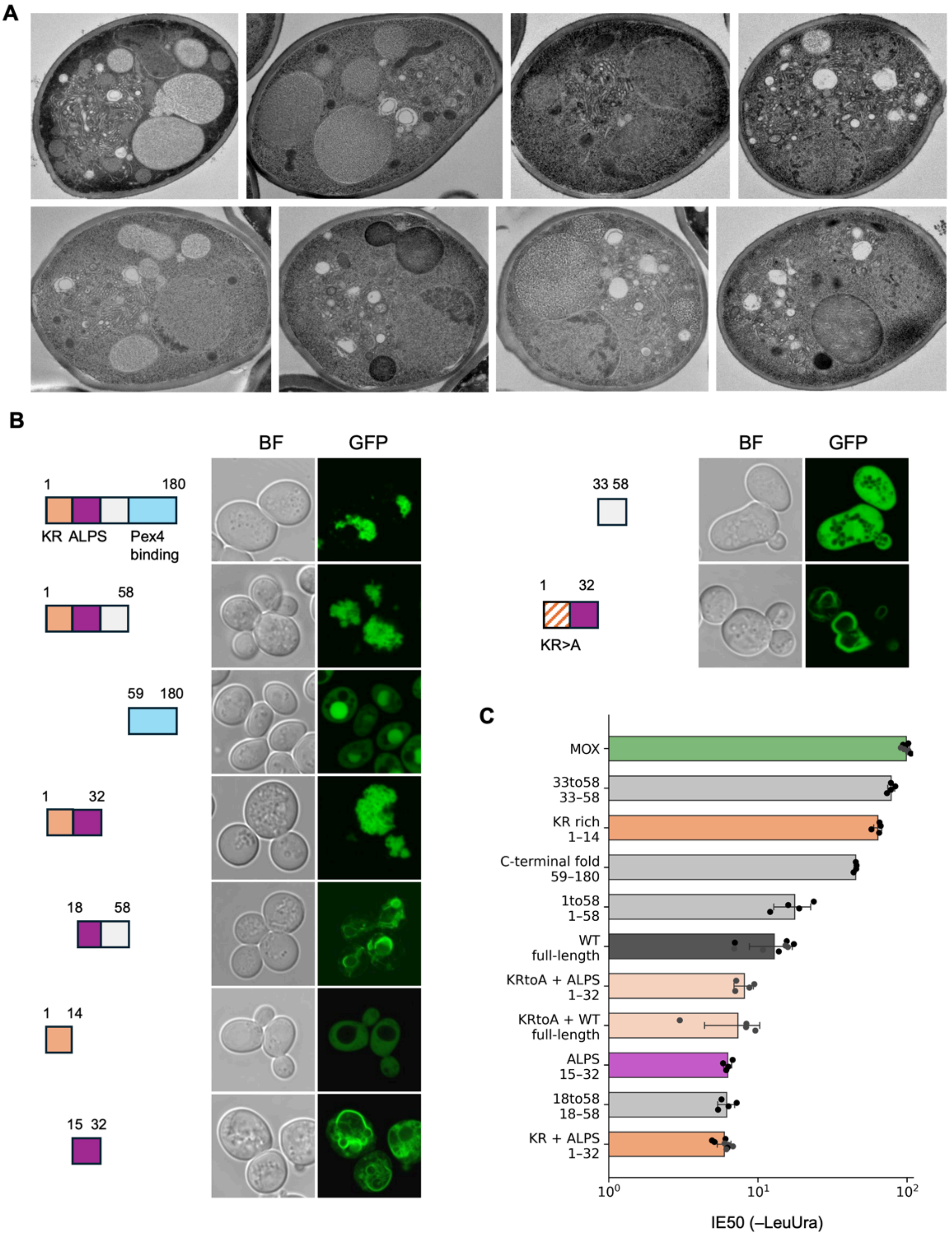


**Figure S37. Structural and sequence determinants of membrane remodeling induced by Pex22 overexpression.**

**(A)** Representative transmission electron microscopy images of cells overexpressing full-length Pex22. For TEM analysis, cells were cultured in –LeuUra medium containing 500 nM aTc at 30°C for 12 h. Extensive proliferation of intracellular membranes was observed, including large structures composed of densely packed tubular membrane profiles that occupied substantial portions of the cell.

**(B)** Subcellular localization and membrane-remodeling phenotypes of Pex22 fragments and mutants. Schematics indicate the Pex22 regions included in each construct, including the N-terminal KR-rich region (KR), putative amphipathic lipid packing sensor (ALPS) motif, and Pex4-binding region. Representative bright-field (BF) and GFP fluorescence images are shown. KR>A indicates substitution of Lys and Arg residues in the KR-rich region with Ala.

**(C)** IE50 values of full-length Pex22 and the indicated fragments and mutants under –LeuUra conditions. Bars show mean ± SD, and points represent biological replicates. IE50 values are plotted on a logarithmic scale.
